# Genomic diversity of elm trees for future treescapes

**DOI:** 10.64898/2026.04.28.720590

**Authors:** Mohammad Vatanparast, Joan F. Webber, Clive Brasier, Juan A. Martín, Richard Buggs

## Abstract

- Dutch elm disease has devastated the treescapes of Europe and North America, causing large elm trees to become largely absent. Several decades of work are now coming to fruition with the availability of potentially resistant cultivars for restoration.
- We analysed genome-wide single-nucleotide polymorphisms derived from whole-genome sequences of more than 200 trees, including elm species, British morphological forms, disease-tolerant Spanish selections and cultivars from several breeding programmes.
- While confirming many specimen labels and multigenerational crossing records, our results also revealed unexpected misidentifications and origins of some samples. This included past hybridisation in some resistant selections thought to be pure species. We find that within our samples, field elm (*Ulmus minor*) is more genetically diverse than wych elm (*U. glabra*). We show that over ten British elm forms sometimes assigned species epithets fall within the genetic range of *U. minor*, *U. glabra* or hybrids.
- With their exotic parentage, many of the elm cultivars introduced into Britain post-epidemic represent a considerable increase in nucleotide diversity over the still numerically large, native *U. minor* and *U. glabra* populations. Currently, the cultivars are being planted mainly as ornamentals. Their long-term arboricultural, adaptive and landscape suitability remains to be demonstrated.

## Introduction

Trees are crucial elements of our culture, nature and economy. They enhance the aesthetics of rural and urban landscapes, accumulate carbon, and provide livelihoods (Chapin III *et al*., 2000; Falkowski *et al*., 2000; Lutz *et al*., 2018). Trees also provide ecosystem services and contribute to biodiversity via interactions with numerous animals and microorganisms (Brockerhoff *et al*., 2017; Chakravarty *et al*., 2019). However, many tree species are threatened by excessive land use, climate change (e.g. extreme droughts, floods and fires and heat waves), overexploitation, and introduced pests and pathogens (Brasier, 2008; Pautasso, 2009; Fisher *et al*., 2012). Pests and pathogens alone killed hundreds of millions of trees worldwide in the last century (Boyd *et al*., 2013). Accelerated international trade in the global economy and worldwide travel of humans continue to transfer novel pests and pathogens from their native range to new continents and landscapes and sometimes challenge indigenous trees to their limits of survival or extinction (Roy *et al*., 2014; Tsouvalis, 2019; Williams *et al*., 2023). Dutch elm disease (DED) (Brasier, 1979), Port Orford cedar root disease (Hansen, 2008), Sudden oak and larch death (Rizzo *et al*., 2002; Brasier & Webber, 2010), ash dieback (Kowalski & Holdenrieder, 2009), emerald ash borer mortality (Herms & McCullough, 2014), chestnut blight (Anagnostakis, 1987), and pine wilt (Hunt, 2009) are all examples of major epidemics in the northern hemisphere caused by introductions.

Dutch elm disease is a notoriously destructive disease caused by Ascomycota fungi of the genus *Ophiostoma*. Two pandemics have occurred across Europe and North America since the early 20th century, the first caused by *Ophiostoma ulmi* (Buisman) Melin & Nannf., the second by the far more aggressive pathogen *O. novo-ulmi* Brasier (Brasier, 2001). Both probably originated from eastern Asia and are capable of infecting and killing European and North American elms (*Ulmaceae*) under field conditions, including common European field elm or English elm (*Ulmus minor* Mill.), European wych elm (*U. glabra* Huds.), European white elm (*U. laevis* Pall.), and American elm (*U. americana* L.). Elm bark beetles (*Scolytus* spp. and *Hylurgopinus rufipes* (Eichhoff)) are the main vectors of DED, as they can transfer *O. ulmi* and *O. novo-ulmi* spores while feeding from the phloem of healthy elms (Fransen & Buisman, 1935; Webber & Brasier, 1984). In Britain, elm is largely a landscape and woodland tree (Clouston & Stansfield, 1979; Richens, 1983), and by 1980 the second DED pandemic had killed more than 25 million elm trees in southern Britain alone (Gibbs, 1978; Brasier, 2001) causing iconic tree losses and landscape transformation (Wilkinson, 1978; Clouston & Stansfield, 1979; Russell & Buggs, 2019; Meiners, 2023).

While European and North American elms tend to be susceptible to *O. novo-ulmi* on inoculation, some Asian species such as *U. pumila* L., *U. davidiana* var. *japonica* Rehder and the Himalayan *U. wallichiana* Planch. are resistant (Heybroek, 1993a) and have contributed to resistance breeding programmes in several countries over the last few decades (Heybroek, 1993b; Smalley & Guries, 1993; Mittempergher & Santini, 2004). So far, over 100 elm hybrids or cultivars, often with very different forms and climatic tolerances, have been released in Europe and North America through breeding programmes, largely with the aim of bringing elms back into the urban landscape (Santini *et al*., 2004). Some of these have been commercially marketed in the UK as DED-resistant such as *Ulmus* ‘Nanguen’ (commercial name Lutece), ‘New Horizon’, ‘Rebona’, ‘Fiorente’ and ‘Sapporo Autumn Gold’ (Walter *et al*., 2026).

A small suite of apparently DED-resistant elm cultivars is available commercially for planting in Britain today. They have been tested to varying extents for DED-resistance but very little for their suitability for the UK environment and for environmental effects on resistance (Domínguez *et al*., 2024). Some, most notably from the Italian and Dutch breeding programmes, have come from hybridisation of *U. minor* with resistant Asiatic elm species (Mittempergher & Santini, 2004). Others are derived from rare *U. minor* trees that have survived DED in the field, most notably from Spain where seven selected genotypes showed good resistance to *O. novo-ulmi* when rigorously tested by inoculation (Martín *et al*., 2015). Similar selections of surviving *U. minor* have been made in Britain, for example by The Conservation Foundation, Natural England and Dr Alec Gunner, but have not so far been subjected to rigorous tests. Some trial deployments of resistant elm cultivars are already underway in Britain on a modest scale by Butterfly Conservation in Hampshire (Brookes, 2024), and on a small scale by various landowners and arboreta (i.e. Coleman, 2025).

In the British countryside, an enormous natural archive of the genetic diversity of elm trees prior to the recent pandemic still survives, since millions of suckers regenerating from the root systems of dying elms often grow until above eight years of age before being killed by DED (Brasier, 1983; Brasier & Webber, 2019). Some small *U. glabra* trees produce viable seeds before succumbing to the disease, again allowing successive generations of small trees (Peterken & Mountford, 1998; Brasier & Webber, 2019). Apart from their susceptibility to *O. novo-ulmi* these trees probably remain well adapted to British conditions. There are also relict populations of large elm trees in refuge areas such as East Sussex and northern Scotland, and rare survivors or disease escapes scattered across Britain (e.g. 624 mature trees recorded in ‘The Great British Elm Search’ (Seddon *et al*., 2025)). *Ulmus minor* is highly morphologically variable and there has also been a debate about the formal taxonomy of elm populations within the ‘*U. minor* group’ in the UK, in particular whether or not different morphological types are clonal cultivars or ‘microspecies’ (cf. Richens, 1983; Hollingsworth *et al*., 2000; Sell & Murrell, 2018). In addition, there is a small, parallel, but more heterogeneous elm population comprising native elms, exotic elms and hybrid cultivars in Britain’s cities, parks, arboreta and some specialist situations.

Including the three European species (*U. glabra, U. minor* and *U. laevis*) there are over 30 species of elms across the northern hemisphere, distributed across six major evolutionary lineages or clades (Whittemore *et al*., 2021). In terms of the genetic structure of the genus *Ulmus* as a whole, Whittemore et al (2021) applied a nuclear approach using RAD-seq loci, complementing earlier molecular phylogenetic studies based on chloroplast DNA (e.g. Wiegrefe *et al*., 1994). Whittemore et al. (2021) resolved the genus into six main evolutionary lineages, which were assigned to six generic sections and three subgenera. The lineages are inferred to have spread to different continents over ca. 10-40 Myr and the European elms, *U. glabra, U. minor* and *U. laevis* each fall within a different evolutionary lineage (Whittemore *et al*., 2021). Indeed, the closest known relatives of *U. glabra* in Section *Ulmus* are *U. rubra* (North America), *U. wallichiana* (Himalaya), *U. laciniata* (Herder) Mayr ex Schwapp. and *U. uyematsui* Hayata (Taiwan and Japan). *Ulmus minor* is nested within a clade comprising section *Foliaceae* along with *U. pumila, U. davidiana, U. castaneifolia* Hemsl.*, U. chenmoui* W.C.Cheng and others (Whittemore *et al*., 2021). *Ulmus laevis* belongs to predominantly American subgenus *Oreoptelea*, closely related to *U. americana* in section *Blepharocarpus* (Whittemore *et al*., 2021).

Genetic diversity within European elms in Europe has been studied using low numbers of markers (e.g. Machon *et al*., 1995; Čurn *et al*., 2014), and has demonstrated widespread clones in *U. minor* (Buiteveld *et al*., 2016). Within Britain, RAPD markers have also revealed widespread clones within *U. minor*, such as *U. minor* ‘Plotii’ (Hollingsworth *et al*., 2000; Coleman, 2000). Use of microsatellite and RFLP markers has suggested that the ‘English elm’, an elm of exceptional architectural form traditionally designated *U. procera or U. minor* var *vulgaris* (Wilkinson, 1978; Clouston & Stansfield, 1979; Richens, 1983) and generally accepted to be the commonest non-woodland elm in England, is a single introduced *U. minor* clone of Roman origin (Gil *et al*., 2004) which is usually described as *U. minor* ‘Atinia’. Whittemore et al (2021) provided evidence that some British elm types recently ascribed specific epithets (Sell and Murrell, 2018) are clones of *U. minor,* and confirmed long suspected interspecific hybridisation in Europe especially involving *U. pumila*, and hybridisation between *U. minor* and *U. glabra*.

Despite developments cited above, genomic analysis of elms is currently at an early stage. A whole genome reference assembly of ‘Ademuz’ elm from Spain was released recently (Pallares Zazo *et al*., 2025), and assemblies for *U. americana*, *U. elongata*, *U. glabra* and *U. parvifolia* were also recently released (Lyu *et al*., 2025; Coleman & Ruhsam, 2024; Wang *et al*., 2025). There have been studies of gene expression differences between susceptible and non-susceptible elm species, and gene expression changes during DED infection (Altmann *et al*., 2018; Chano *et al*., 2025). However, there has been no broad-scale genomic analysis of extant elm species and varieties present in one country, nor of recently released commercial hybrid cultivars. In the present study, we sampled over 200 elms via whole-genome sequencing to explore the genomic composition of British elms and elms potentially available for treescape restoration in Britain.

## Materials and Methods

### Sampling

Leaf materials of elm species and known DED-resistant elm cultivars were collected from the living collections of Royal Botanic Gardens, Kew, Sir Harold Hillier Gardens, Grange Farm Arboretum (Matthew Ellis), the Frank P. Matthews Ltd nursery, and field-collected materials across the UK from 2015-2024 (Table S1). We also included 49 *Ulmus* samples from the clonal bank at the Puerta de Hierro Forest Breeding Centre in Madrid, Spain, and 21 samples representing the Sell and Murrell elm microspecies (Sell & Murrell, 2018) from Kew arboretum nursery, from East Anglia collected by Brian Eversham and Alex Prendergast and sent to us by Cicely Marshall (University of Cambridge), and from Wakehurst collected and sent to us by Christopher P. Cockel (Kew Gardens).

### Whole Genome Sequencing

Collected leaf materials were preserved in paper or plastic bags with silica gel and stored at room temperature. Genomic DNA was extracted using a modified CTAB method with a Sorbitol pre-wash step (Inglis *et al*., 2018). The quality and quantity of extracted genomic DNA were checked with 1% agarose gel, Quantus Fluorometer (Promega) and NanoDrop 2000 (Thermo Scientific). Due to the viscosity of some extracts, we used Agencourt AMPure XP (Beckman) beads to purify DNA before library preparation. Library preparation was conducted using the egSEQ Enzymatic DNA protocol. Initially, DNA fragments were generated and selected to achieve an approximate insert size of 300-400 bp. Subsequently, end repair, A-tailing, and adaptor ligation steps were performed. Then purification and PCR amplification were carried out to enrich the library. Further purification was conducted to ensure high-quality libraries. Prior to sequencing on the MGI DNBSEQ-T7 platform with 150-bp paired-end reads, circularisation steps were implemented. The library preparation and sequencing were carried out by Edinburgh Genetics Ltd (Edinburgh, UK).

Raw reads were quality-filtered using Trimmomatic v0.33 (Bolger et al., 2014), applying a four-base sliding window with a mean Phred-quality threshold of 15 and assuming Phred+33 quality encoding. Reads shorter than 70 bp after trimming and reads containing residual adapter sequences were discarded. Read quality before and after trimming was assessed using FastQC v0.11.9 to confirm the removal of adapter contamination and overrepresented sequences.

### SNP Calling

We used haplotype 1 of the *U. americana* genome (NA87034HAP1_v1_1) as a reference for SNP calling which was the only chromosome-level reference genome available for elms (Spring 2023). The filtered sequencing reads were mapped to the *U. americana* reference genome using BWA MEM2 v2.2.1 (Vasimuddin *et al*., 2019). For each sample, the read group information was appended and alignments were sorted using SAMtools v1.6 (Danecek *et al*., 2021). PCR duplicates were removed using the MarkDuplicates option and REMOVE_DUPLICATES=true of the Picard v3.3.0 (https://github.com/broadinstitute/picard). Sorted BAM files were filtered to keep only reads with a mapping quality score of at least 30. The filtered BAM files were then indexed and quality metrics were generated using SAMtools flagstat option. Variant calling was performed using GATK v4.5.0.0 HaplotypeCaller for each of the 14 chromosome intervals; the three unplaced scaffolds in the reference assembly were excluded. Joint genotyping was performed using GATK GenotypeGVCFs after chromosome-interval genomic databases had been generated using GenomicsDBImport (Van der Auwera & O’Connor, 2020). Variants were hard-filtered using GATK VariantFiltration. Sites were flagged if they met any of the following criteria: quality by depth (QD) < 2.0, Fisher strand bias (FS) > 60.0, root mean square mapping quality (MQ) < 50.0, mapping-quality rank-sum test (MQRankSum) < −2.0, or read-position rank-sum test (ReadPosRankSum) < −2.0. Individual genotype calls with genotype quality (GQ) < 20 were flagged as lowGQ. GATK SelectVariants was then used to retain SNPs passing all site-level filters. The filtered VCFs were then processed with BCFtools v1.21 (Danecek *et al*., 2021) to keep only biallelic SNPs. We used PLINK v2.0.0-a.6.20LM (Chang *et al*., 2015) to remove duplicate variants, exclude variants with any missing genotypes (--geno 0), apply a minor-allele-frequency threshold of 0.05 and a Hardy–Weinberg equilibrium P-value threshold of 1 × 10, and perform LD pruning using --indep-pairwise 50 5 0.2. These filters were applied to the datasets used for PCA, kinship and ADMIXTURE analyses. The genome-wide diversity analyses used the unpruned biallelic SNP dataset without a minor-allele-frequency filter, as described below.

### Kinship analysis

In order to detect the clonal replicates, we built a KING relatedness matrix using PLINK v2.0.0-a.6.20LM --make-king algorithm (Manichaikul *et al*., 2010) on LD-pruned SNPs and plotted individuals and kinship coefficients using the R package igraph v2.0.3 (Csárdi *et al*., 2025). We identified individuals as clones if they had an estimated kinship coefficient of 0.30 or above (Manichaikul *et al*., 2010), as first-degree relatives if they had an estimated kinship coefficient from 0.177 to less than 0.30, and less than 0.177 were interpreted as second-degree relationships.

### Principal component analysis

We used the --pca option implemented in the PLINK v2.0.0-a.6.20LM (Chang et al., 2015) to obtain the eigenvalues and eigenvectors for the principal component analysis (PCA). PCA plots were constructed using the Bokeh Python library, and the variance explained by each principal component was visualized using the factoextra (https://github.com/kassambara/factoextra) and ggplot2 R packages (Wickham, 2011).

### Phylogenetic analysis

To place our accessions within a genus-wide phylogenetic framework, we assembled a combined RADseq dataset comprising 81 accessions generated in this study and 95 accessions from the RADseq phylogeny of Whittemore et al. (2021), for a total of 176 taxa. The combined dataset spanned 36 species of *Ulmus*, five species of *Zelkova* Spach*, Hemiptelea davidii* (Hance) Planch., and *Planera aquatica* J.F.Gmel. (the latter three genera used as outgroups). Artificial hybrid cultivars were excluded because of their known reticulate ancestry. Raw sequence reads from Whittemore et al. (2021) were downloaded from the NCBI Sequence Read Archive (BioProject PRJNA672171), trimmed as above using Trimmomatic v0.33 and processed jointly with our trimmed reads through the ipyrad v0.9.105 pipeline (Eaton & Overcast, 2020), using default parameters with 12,213 RAD loci as reference (Whittemore et al. 2021). The final alignment comprised 400,461 sites (200,643 unique site patterns) after removal of 255 gap-only columns. The alignment contained 26.85% gaps and 89.06% invariant sites. Maximum-likelihood (ML) phylogeny was inferred using RAxML-NG v1.2.2 (Kozlov *et al*., 2019) under the GTR+G substitution model. Twenty independent ML tree searches were conducted from 20 parsimony starting trees. Branch support was assessed using 100 non-parametric Felsenstein bootstrap replicates (Felsenstein, 1985). The best ML tree (log-likelihood = −1,126,230.33; AIC = 2,253,176.66; BIC = 2,257,078.99) was used, with bootstrap support values mapped onto its branches. The tree was rooted using *Planera aquatica*, *Hemiptelea davidii*, and *Zelkova* species as the outgroup, following Zhang et al (2022), and Gao et al. (2023). To obtain a Split Network visualisation, we used SplitsTree App v6.0.0 (Huson & Bryant, 2024) using the NeighbourNet method (Bryant, 2003).

### Admixture analysis

To estimate ancestry proportions across the sampled elms we conducted model-based clustering using ADMIXTURE v1.3.0 (Alexander *et al*., 2009). Multiple ADMIXTURE analyses were performed on overlapping subsets of the dataset, each enabling ancestry estimates of certain elms. For each dataset, ADMIXTURE was run on LD-pruned SNPs with clusters (K) ranging from 2 to 20, using 10-fold cross-validation (--cv=10), random seeds (-s), and 500 bootstrap replicates (-B) for standard error estimation. The optimal number of ancestral populations (K) was identified as the value minimising cross-validation (CV) error. The best-supported value of K for each analysis was selected using the plot function in R (R Core Team 2025) using cross-validation estimates. Because CV error is known to be an imperfect guide when data are hierarchically structured (Lawson *et al*., 2018), we examined results across multiple K values and interpreted ancestry proportions in the context of our PCA (Figs 1 and 2), and phylogenetic results (Fig. 3). Briefly, the first dataset comprises elms originally collected from trees present in the natural environment, excluding all complex hybrids (Elms-120), but including widespread clones like ‘Atinia’ that have been in Britain for many centuries. The second dataset comprised the Elms-120 accessions plus one representative from each selected clonal group of complex hybrids, giving 132 accessions in total. To test the admixture ancestry of DED-resistant Spanish *U. minor* clones (i.e. ‘Ademuz’), we used the f2 statistics from the AdmixTools 2 R package (Patterson *et al*., 2012). Using LD-pruned SNPs data from PLINK, we tested Spanish clones individually (E081, E126, E129, E132, E200 and E209) using qpAdm as target, European *U. minor* and *U. glabra* as left-hand sources, and four North American taxa (*U. mexicana, U. crassifolia, U. ismaelis* and *U. serotina*) as right-hand outgroups.

**Fig. 1.**
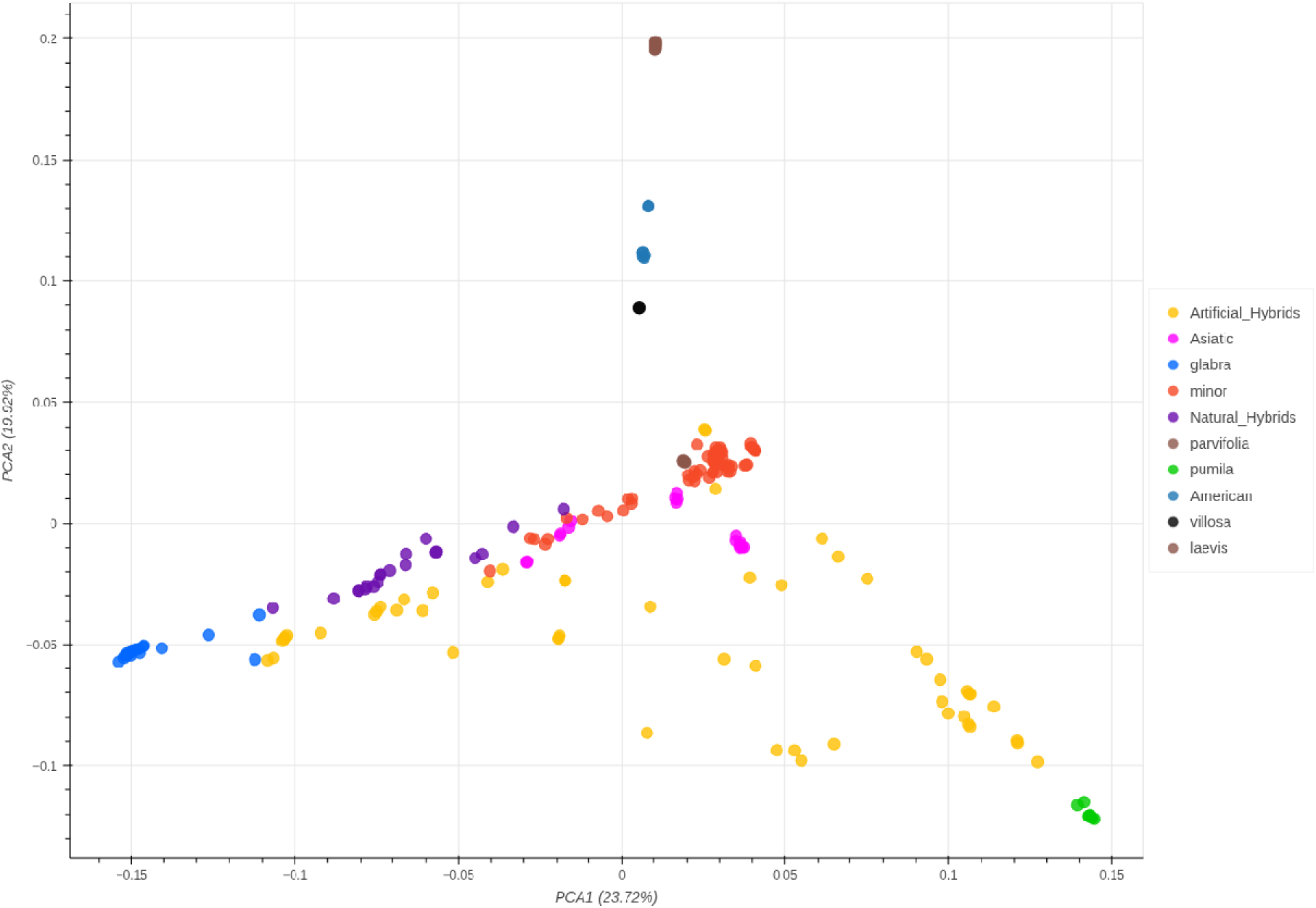
Principal component analysis (PCA) of whole-genome sequence data from all elms sequenced (204 elms), including representatives from all three subgenera, *Ulmus*, *Indoptelea* and *Oreoptelea* (Whittemore *et al*., 2021). Principal components 1 and 2 are shown. Other principal components are shown in Fig. S2. Colours represent major groupings, based on living collection labels for each sample (some of which we later re-identified - see main text). A zoomable HTML version of this figure with labels by hovering is provided as a Supplementary File.

**Fig. 2.**
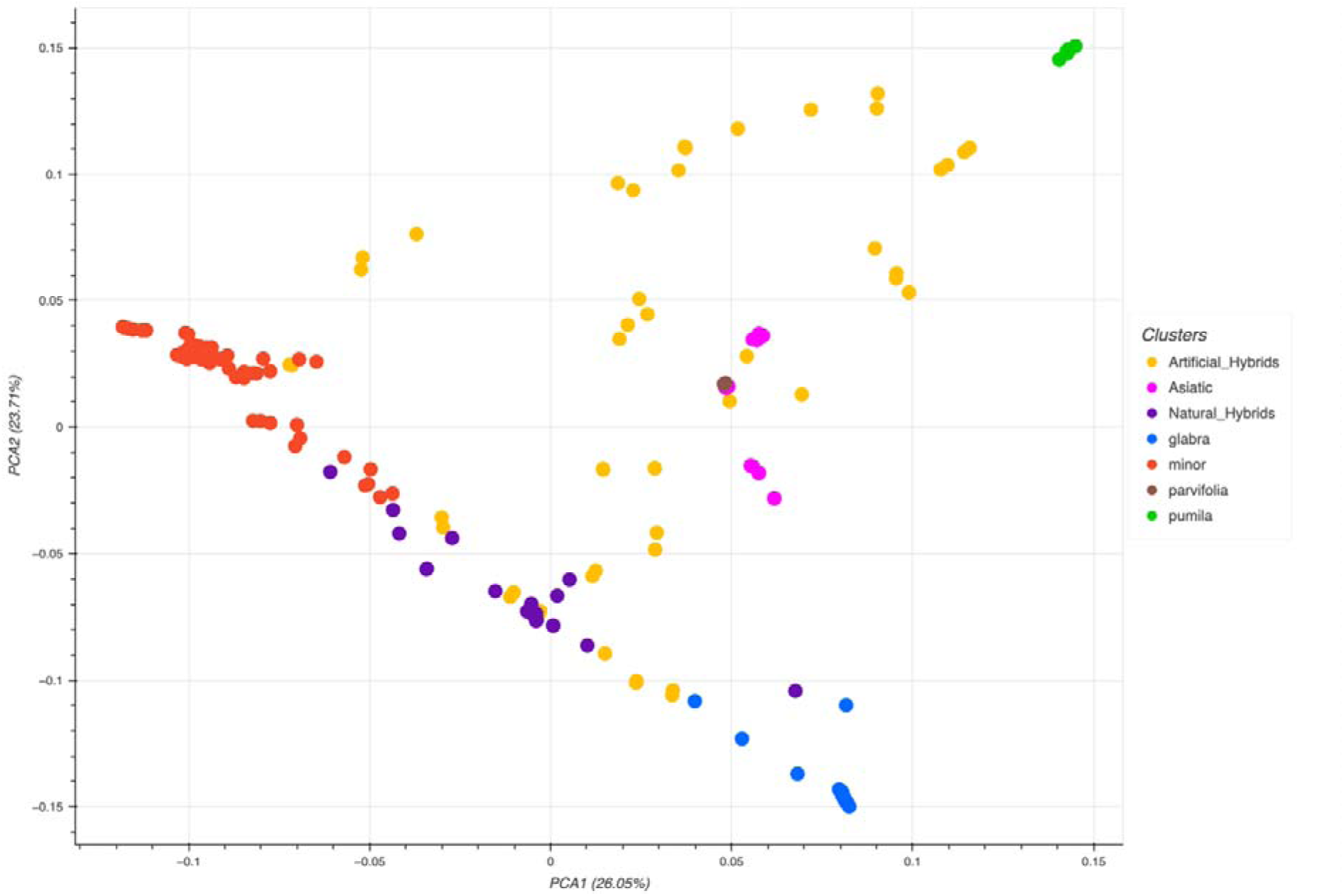
Principal component analysis of whole-genome sequence data from all elms sequenced from *Ulmus* subgenus *Ulmus* (180 elms). Principal components 1 and 2 are shown. Other principal components are shown in Fig. S3. A zoomable HTML version of this figure is provided as a Supplementary File.

**Fig. 3.**
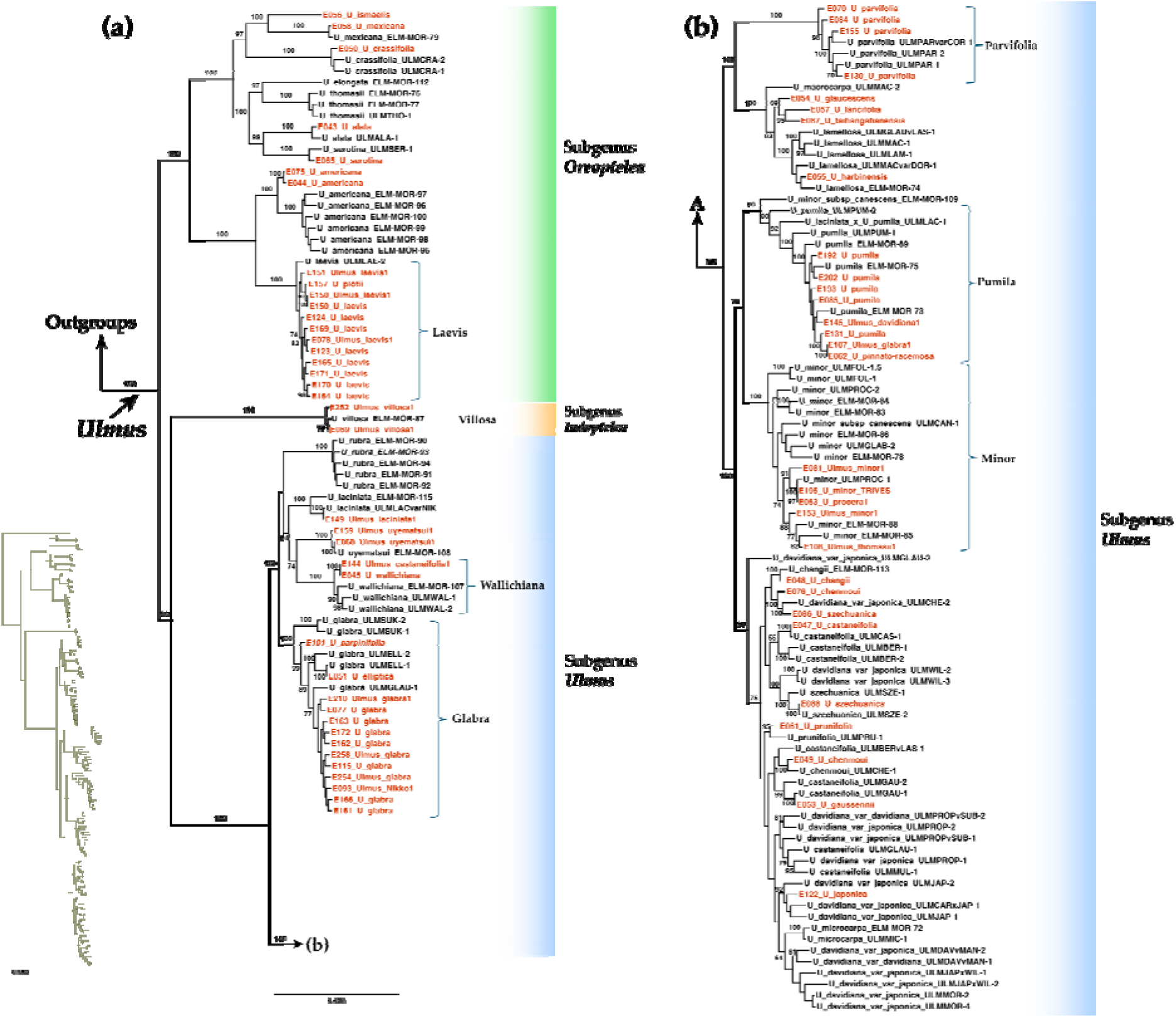
Maximum-likelihood phylogeny of elms inferred from 4,651 concatenated RADseq loci. The dataset comprised 81 accessions newly sequenced in this study, shown in red, and 95 accessions from Whittemore et al. (2021), shown in black. The complete tree was divided between panels (a) and (b) for readability; an overview is shown in the inset. Numbers beside branches indicate bootstrap support values ≥70%. Subgeneric classifications follow Whittemore et al. (2021).

### Genome-wide patterns of diversity within clusters

To compare genetic diversity between *U. minor* and *U. glabra*, we selected a balanced subset of 14 non-admixed individuals per species based on the ADMIXTURE analysis (Fig. 5a). For each species, eight British and six Spanish individuals were selected. Genetic diversity was calculated from the filtered biallelic SNP dataset for both species, without LD pruning and minor allele frequency filtering. Individual observed heterozygosity (*Ho*) was estimated from the PLINK --het option, following Anderson et al. (2010), as (N − O)/N, where N is the number of nonmissing genotypes, and O is the observed number of homozygous genotypes. Expected heterozygosity (He) was estimated from allele frequencies using PLINK --freq counts, with an unbiased correction for sample size. Chromosome-level and 1-Mb window summaries of Ho were calculated from genotype counts using PLINK --freqx output. We also estimated nucleotide diversity (π), chromosome-level and 1-Mb window π using VCFtools v0.1.17 (Danecek *et al*., 2011) from the same biallelic SNPs. We treated the estimated π from VCFtools as a relative comparison between species rather than an absolute genome-wide comparison, since the invariant sites are not included in the VCFtools analysis.

We also analysed the genome-wide distribution of biallelic SNP density without LD pruning and minor allele frequency filtering in elm groups using the CMplot R package (Yin *et al*., 2021). We did this for the following groups identified using the PCA analysis: (A) 14 individuals of *U. minor*; (B) 14 individuals of *U. glabra*; (C) 17 hybrids between *U. minor* and *U. glabra*; (D) 12 trees of Dutch hybrid cultivars (i.e. ‘Dodoens’, ‘Fagel’, ‘Lutece’, ‘Plantijn’/‘Plantyn’, and ‘Wanoux’); (E) 32 complex hybrids (i.e. *U. canescens*, ‘Fiorente’, FL493, FL610, ‘Lobel’, ‘Den Haag’, ‘Morfeo’, ‘Patriot’, ‘Plinio’, ‘San Zanobi’), and Spain hybrids; (F) 12 individuals of *U. laevis*; and (G) four individuals of *U. parvifolia*.

## Results

In this study, we generated whole-genome sequencing data for 204 elms using short-read paired-end reads (150 bp). Photographs of selected sequenced elm trees are in Fig. S1. Sequencing generated approximately 159 million clean paired-end read pairs per accession, corresponding to approximately 47.75 Gb of clean sequence data per accession. Sample-specific read counts and quality metrics are provided in Table S1. Over 2.7 million final high-quality LD-pruned SNPs were retrieved across 204 elms, although the final SNP sets varied among the datasets and analyses involving 204, 180, 132 or 120 elm accessions. The average percentage of reads mapped to the *U. americana* genome was 95.2% across all samples.

### Kinship analysis

Kinship analysis revealed numerous clones in our sequenced samples, some of which correspond with the original labels of the samples and some of which do not (Table 1). All four samples of the complex hybrid FL493, also known as ‘Wingham’ are clones. Three samples labelled ‘Fiorente’ are clones, as are three samples labelled ‘Wanoux’, two samples labelled ‘Lutece’, two samples labelled ‘Plantyn’ and two samples labelled ‘Ademuz’. Three samples labelled ‘Dodoens’ are clones, but one sample labelled ‘Plantyn’ is the same clone, suggesting that the latter is mislabelled. Two samples labelled ‘New Horizon’ are clones, but one labelled ‘Rebona’ is the same clone, suggesting a mislabelling of the latter, as these two cultivars should be full siblings, not clones. Two samples from the Spanish programme labelled with the synonyms *U. procera* and *U. minor* var *vulgaris* from Puente de San Fernando (BCC-F5C4; E207 and BCC-F5C10; E196) were clones. Some sets of trees or clones are also suggestive of mislabelling in living collections, especially in the Melville hedge of elms at Wakehurst Place (UK). These sets include:

- *Ulmus* ‘Christine Buisman’ (identified with confidence) from Alice Holt (E125) and *U. coritana* from Wakehurst (1975-2700)
- *U. plotii* from Wakehurst (1975-6178) and *U. carpinifolia* Wakehurst (1975-6183)
- *U. plotii* × *carpinifolia* from Wakehurst (1977-1491) & *U. coritana* var. *angustifolia* from Wakehurst (1975-2704)
- *U. minor* from Cambridge (E231) and *U. serratifrons* from Cambridge (E234) and *U. minor* ‘Keyston’ Kew (2015-69*1)
- *U. coritana* × *plotii* (Wakehurst 277-75.02710) and *U.* × *diversifolia* (Wakehurst 1975-6176)
- *U. minor* from Kew (1973-11710*1) and *U. laciniata nikkoensis* from Kew 1973-11708*1
- *Ulmus* ‘San Zanobi’ from Kew (2019-2712*2; E158), and Dutch hybrid ‘Lobel’ from Alice Holt (E135)
- *U. plotii* from Kew (2010-1842*1) is *U. laevis* based on PCA, phylogenetic and expert inspection.

**Table 1.**
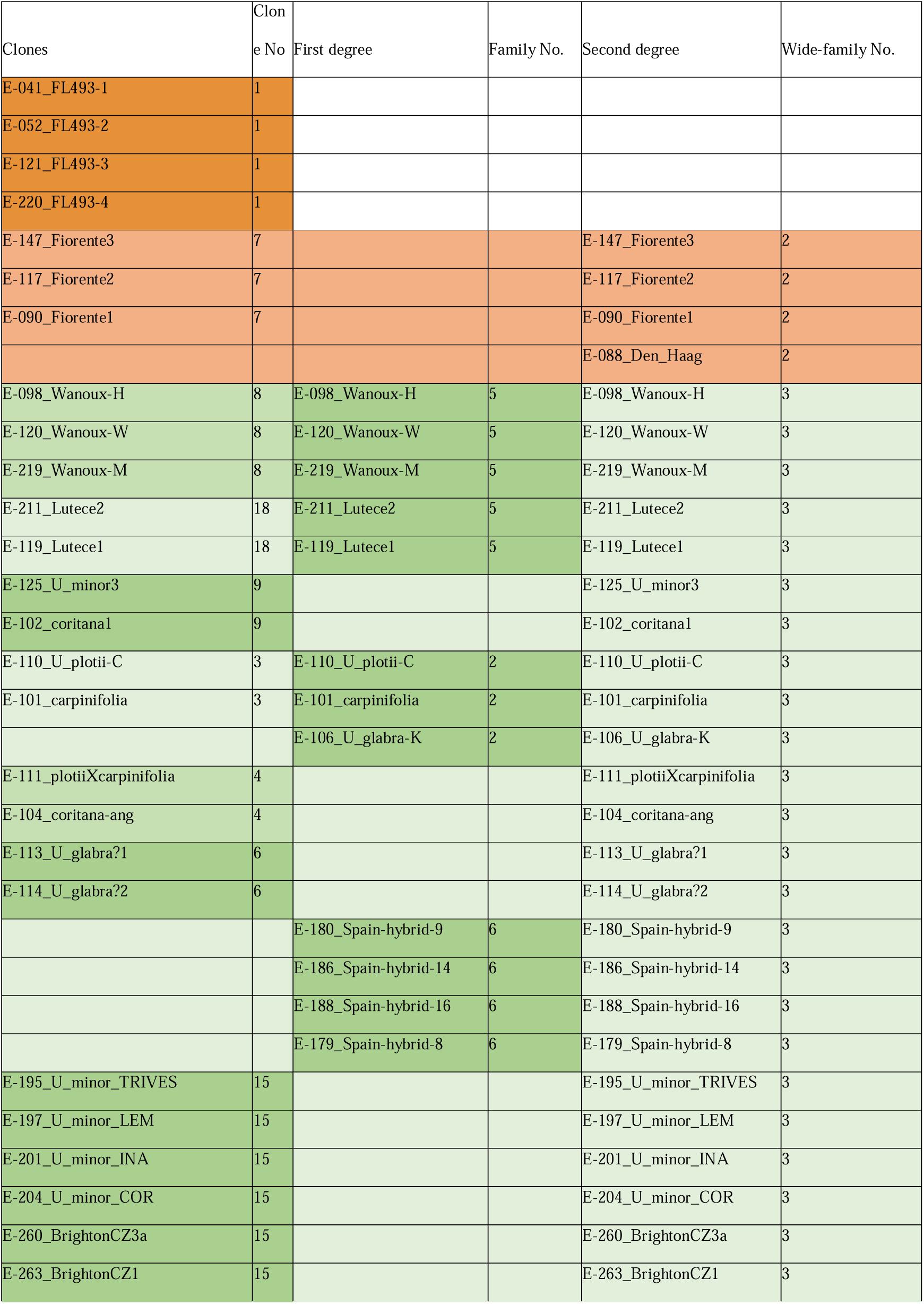
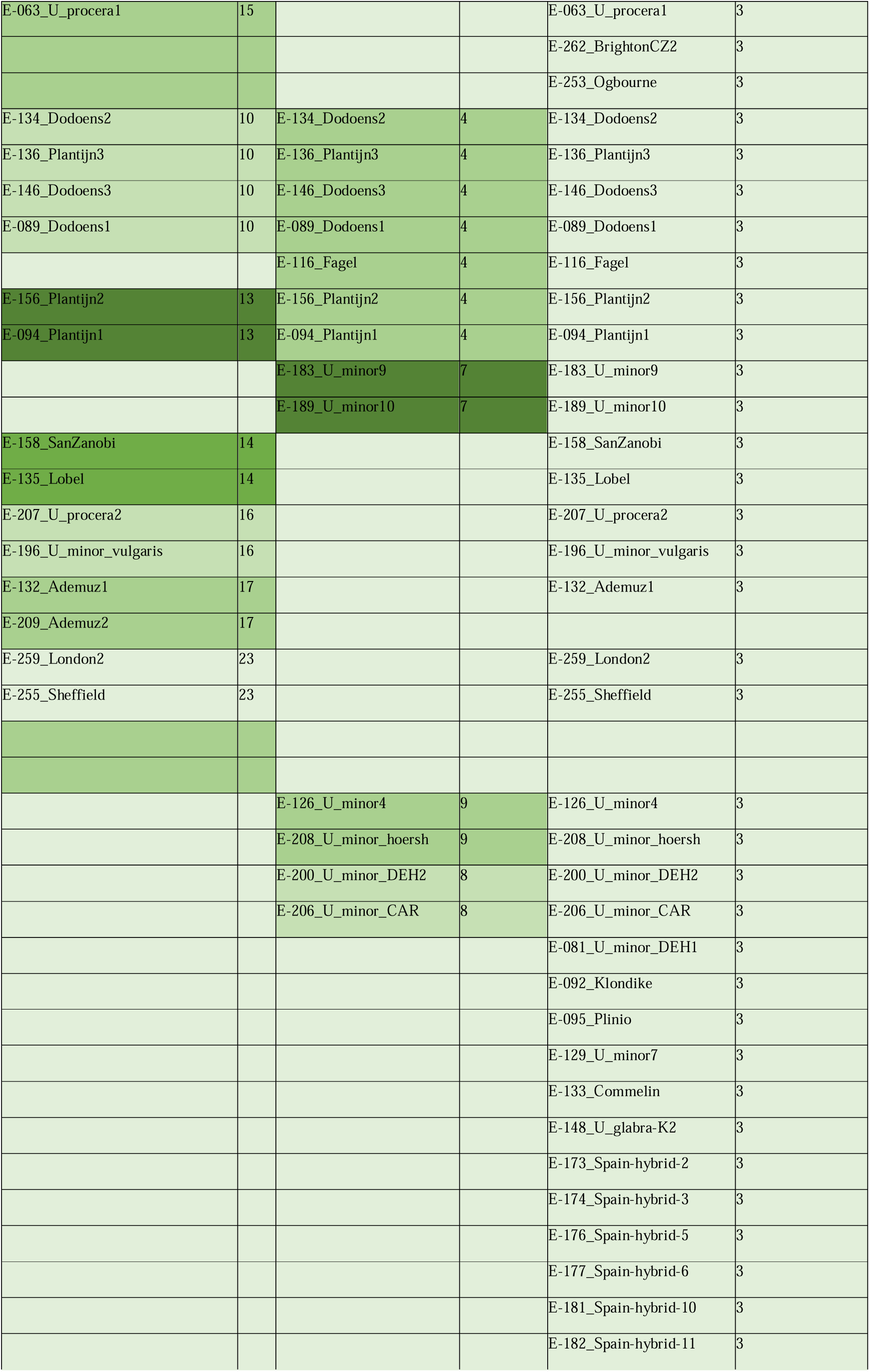
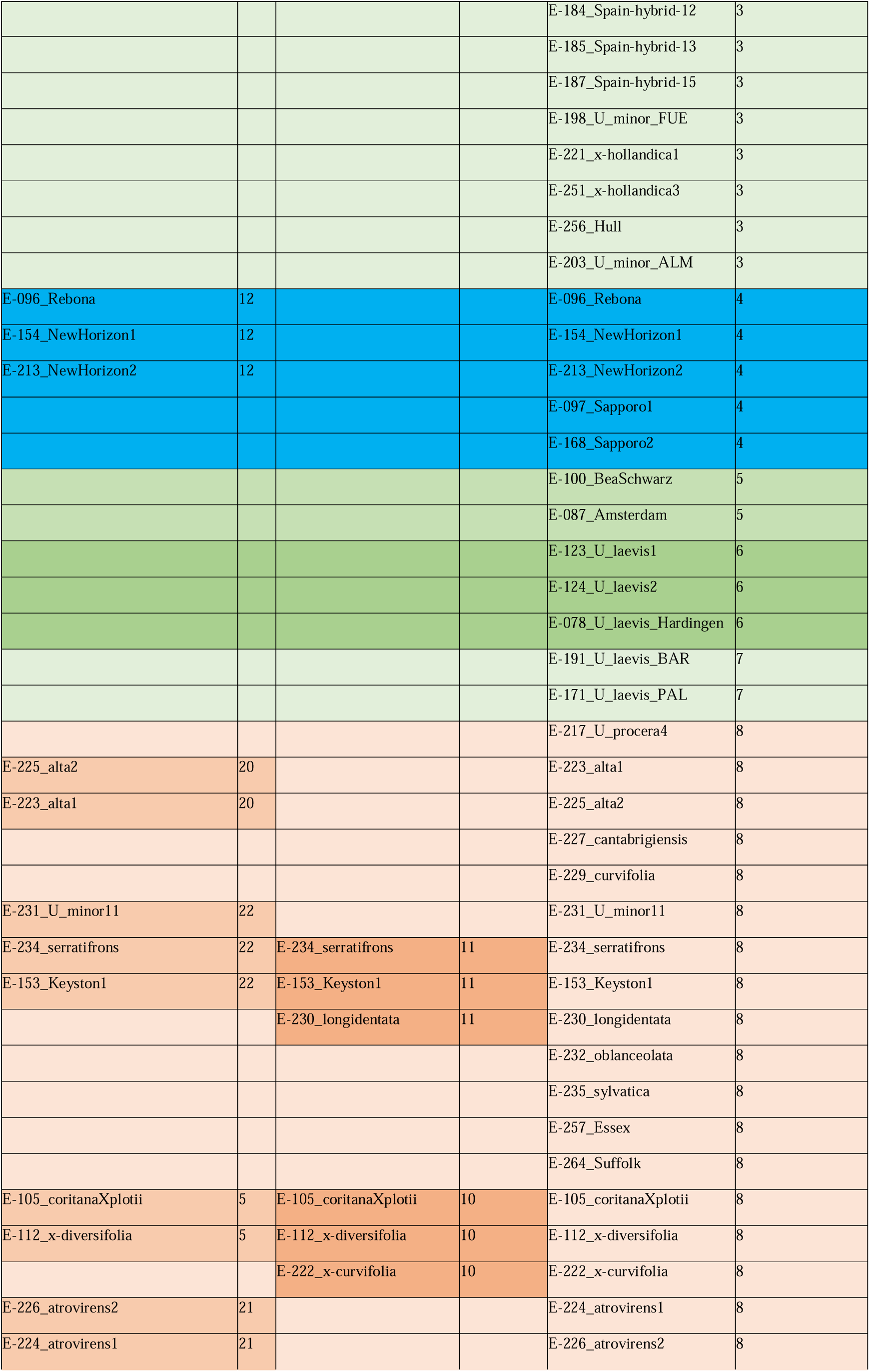
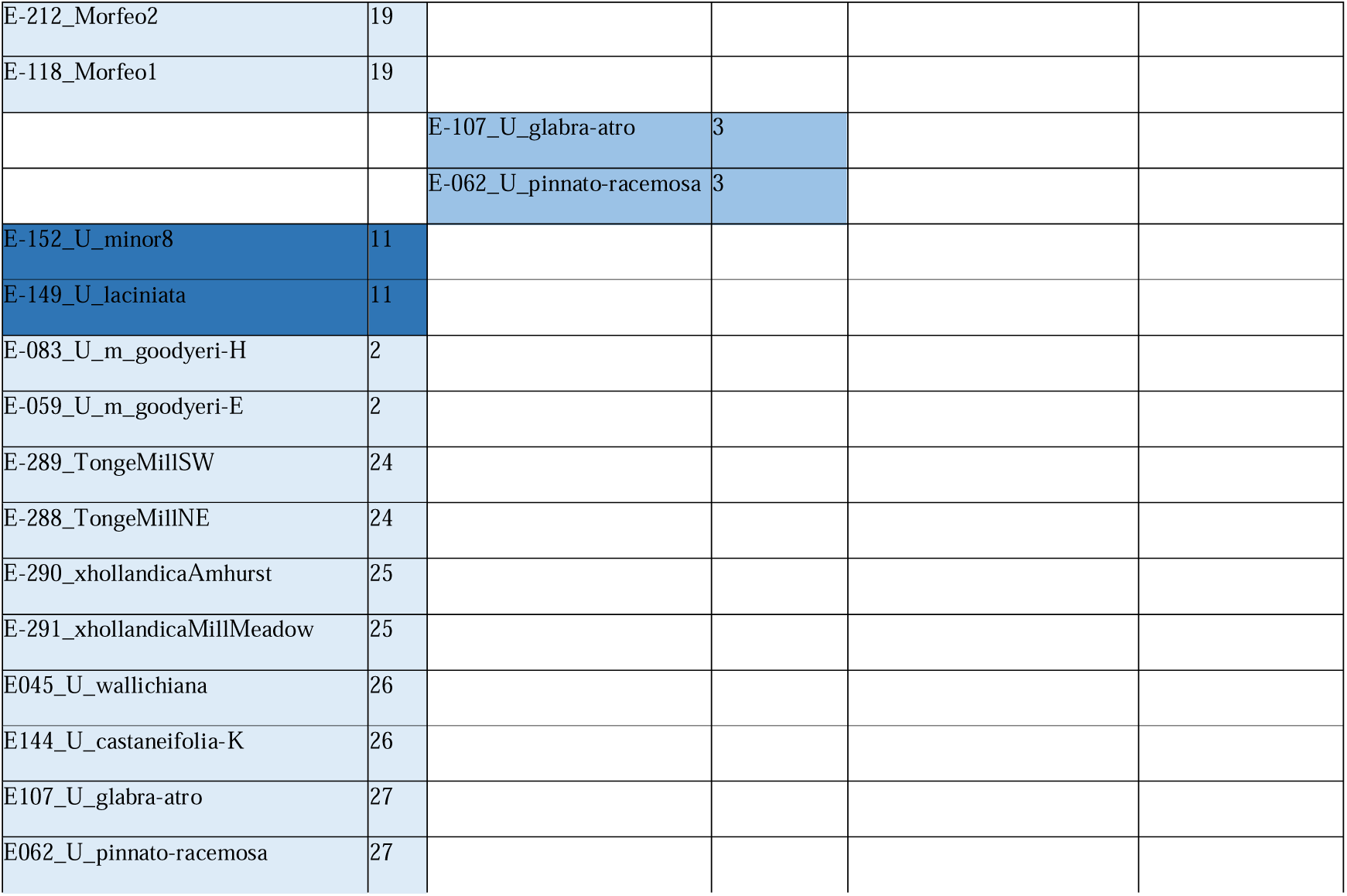
Relatedness among sequenced samples, calculated using the KING algorithm.

Other clonal sets suggest widespread historical planting of single clones in the wider environment:

- Clonal Set 1. Four *U. minor* from Galicia in northwestern Spain labelled: P. Trives (BCC-F5C6), M. Lemos (BCC-F6C2), Iñas (BCC-F9C2), La Coruña (BCC-F11C6) and two elms from the Brighton control zone (E260 and E263) and an accession labelled *U. procera* from Grange Farm Arboretum (E063). This would appear to be the very widely planted and distinctive English elm clone, in Britain often historically designated *U. procera* or *U. minor* var *vulgaris*; but more recently redesignated *U. minor* ‘Atinia’ and suggested to be an Italian clone introduced into England by the Romans from Galicia (Gil *et al*., 2004). In the PCA analysis (see below) these samples cluster at one extreme of the overall distribution.
- Clonal Set 2. An elm sampled in 2015 by a roadside near Glynde, East Sussex (E262) and another by a farm track near Ogbourne St George in Wiltshire (E253). In the PCA (see below) these are the closest samples to Clonal Set 1.
- Clonal Set 3. Large surviving street trees collected from Islington, London (E259) and Nether Edge, Sheffield (E255). In the PCA (see below) these appear to be hybrids between *U. minor* and *U. glabra* and could therefore potentially be classed as *U. × hollandica*.

A large number of first and second-degree relatives were also found among the samples (see Table 1).

### Principal component analysis

In a PCA of all *Ulmus* samples sequenced (204 elms), the clustering of accessions fits well with most assigned species labels, the known parentage of some trees labelled as hybrids, and the above kinship analysis (Fig. 1). The first principal component (PC1) summarises ca. 23% of the variation and primarily separates *U. glabra* at strongly negative values from *U. minor* and, further along the same axis, many artificial hybrids and *U. pumila* at positive values. Natural hybrids occupy an intermediate position between the *U. glabra* and *U. minor* clusters. The broad *U. minor* cluster lies on the positive side of PC1, with Clonal Set 1 occupying one end of that wider distribution; Clonal Set 2 is close to Clonal Set 1. The separation of *U. glabra* and *U. minor* on PC1 is consistent with their placement in distinct evolutionary lineages, Section *Ulmus* and Section *Foliaceae*, in the molecular phylogeny of Whittemore et al. (2021) and in our phylogeny (Fig. 3).

The second principal component (PC2), which summarises ca. 20% of the variation, separates the American species, *U. villosa*, and *U. laevis* from the main European clusters. The American species (*U. alata, U. crassifolia, U. ismaelis, U. mexicana, U. serotina and U. americana*) form a compact cluster at positive PC2 values. *Ulmus villosa* lies close to this American cluster, while *U. laevis* is separated still further in the positive direction of PC2. *Ulmus parvifolia* forms a distinct cluster at slightly positive PC1 and PC2 values. The Asiatic species (*U. castaneifolia*, *U. changii*, *U. chenmoui*, *U. gaussenii*, *U. prunifolia*, *U. szechuanica*, *U. japonica*) are distributed among several groups near the centre of the plot, between the European species and the more divergent *U. pumila* cluster. A broad continuum of artificial hybrids extends from the vicinity of *U. minor* toward the Asiatic groups and *U. pumila*, reflecting the widespread use of Asiatic elms, especially *U. pumila*, in breeding programmes.

Interspersed among the groups of worldwide native elm species in the PCA are some anomalous samples. Accession E093 from Sir Harold Hillier Gardens, labelled as ‘Nikko’ (a name usually used for a cultivar of *U. laciniata* or a hybrid thereof) clusters with *U. glabra*. In contrast, Kew accession E149 *U. laciniata nikkoensis* is found in the Asian pacific coast cluster about half-way between *U. minor* and *U. pumila*, close to two accessions of *U. uyematsui* (E159 and E068). Accession E152 from Kew, labelled *U. minor*, was anomalously also in this Asian pacific cluster, suggesting it is mislabelled. Kew accession *U. plotii* (E157) (normally considered part of the *U. minor* complex) falls within a cluster of *U. laevis* samples, suggesting that it is in fact *U. laevis*; further examination of its morphology also suggests that it is *U. laevis*. Wakehurst accession E107 labelled *U. glabra* ‘atropurpurea’ clusters with *U. pumila*, suggesting that either the accession is mislabelled or the taxonomy of the cultivar ‘atropurpurea’ needs revision.

We sequenced two samples of the Chinese species *U. castaneifolia*: one being Kew accession E144 and the other from Grange Farm arboretum (E047). Of these, the Kew sample clustered with *U. wallichiana* (Himalayan elm) and the Grange Farm sample with other Chinese species including *U. szechuanica*. An accession from Grange Farm labelled as the Mediterranean species *U. canescens* (E046), is found close to a hybrid from the Spanish programme (E176), which is ((*U. minor* × *U. pumila*) × *U. minor*). This suggests either that *U. canescens* has a natural hybrid origin, or this specimen is mislabelled. In Spain, *U. minor* and *U. pumila* hybridise quite easily, and many interspecific hybrids are found in the landscape (Cogolludo-Agustín *et al*., 2000).

The PCA of genome-wide SNPs from only *Ulmus* subgenus *Ulmus* (180 elms) shows that *U. minor* forms a large cluster at the negative end of PC1, which explains ca. 26% of the variation, with samples collected from Britain and Spain interspersed throughout (Fig. 2). Also falling within the *U. minor* cluster are samples labelled with microspecies names in the Sell and Murrell (2018) taxonomy, including: *U. longidentata*, *U. serratifrons*, *U. coritana*, *U. alta*, *U. atrovirens*, *U. cantabrigiensis*, *U. oblanceolata*, *U. sylvatica* and *U. coritana × plotii. Ulmus glabra* forms a distinct cluster at the positive end of PC1 and at strongly negative values of PC2. Samples positioned between the *U. minor* and *U. glabra* clusters form an intermediate arc and are strongly suggestive of hybrids and introgressions between the two species, often known as *U. × hollandica*. One of the samples in this intermediate group was labelled *U. × hollandica* and another as ‘Wentworthii’, a known cultivar of *U. × hollandica*. However, other samples in this intermediate group had a variety of different labels in the source collections, including *U. carpinifolia, U. coritana, U. plotii, U. glabra, U. minor, U. procera and U. plotii × carpinifolia*. Clonal Set 3 is also found in this group. Their positions in the PCA support the hypothesis that these samples are of hybrid origin between *U. minor* and *U. glabra*, and the admixture analysis presented below is consistent with this interpretation.

The second principal component (PC2 = ca. 23%) further separates the European groups from the more Asiatic elms. *Ulmus glabra* occupies the most negative PC2 values, whereas artificial hybrids, Asiatic taxa, and especially *U. pumila* occupy increasingly positive PC2 values. *Ulmus parvifolia* and the other Asiatic samples lie between the European clusters and the strongly separated *U. pumila* cluster. A broad spread of artificial hybrids extends from the vicinity of *U. minor* toward the Asiatic groups and *U. pumila*, consistent with the extensive use of *U. pumila* and other Asiatic elms in breeding programmes. Selections attributed to *U. minor* with some resistance to DED, such as ‘Ademuz’, ‘Klondike’ and ‘Bea Schwarz’, occur on the edge of the *U. minor* cluster in the direction of *U. glabra* and the artificial/Asiatic material. A sample of *U. thomasii* from Wakehurst is found within the *U. minor* cluster, which again is likely to reflect mislabelling, as *U. thomasii* is native to North America and belongs to a different subgenus from U. minor.

We consider Clonal Set 1 to represent the “English elm” *U. minor* ‘Atinia’ (syn. *U. procera* and *U. minor* var. *vulgaris*). Among our samples were six with the above attributions that did not cluster with Clonal Set 1 in the PCA. A sample from Frank P. Matthews Ltd labelled *U. procera* “Dodwell large leaves wind Rd 16” clustered with *U. glabra*; and samples from the same nursery labelled *U. procera* “Dodwell normal leaves” and “Staffordshire” were in the putative *U. × hollandica* cluster. A sample from Grange Farm arboretum labelled *U. procera* was in the *U. minor* cluster, but two from Spain labelled as *U. procera* and *U. minor* var. *vulgaris* “Puente de San Fernando” (BCC-F5C4; E207 and BCC-F5C10; E196) were in a different region of the *U. minor* cluster. The latter two Spanish samples are known to be the same clone in the Spanish collection, but their classification as *U. procera* was based solely on leaf morphology and re-examination in the light of these molecular results suggests that though they have a leaf shape similar to ‘Atinia’, they are slightly smoother in texture (Juan A. Martín, pers. obs.). If Clonal Set 1 is true ‘English elm’, in agreement with the claim by Gil et al. (2004), based on eight samples, that all English elms are a single clone, then the six samples labelled as such but not falling within Clonal Set 1 are most probably mislabelled, although the possibility that English elm is a convergent *U. minor* phenotype expressed by several different genotypes showing only subtle morphological differences cannot yet be ruled out. Unfortunately, we have not yet sequenced Forestry Commission *U. procera* clone SR4, which has been widely used in the UK as a standard for testing the aggressiveness of DED *Ophiostoma* genotypes (e.g. Sutherland et al., 1997) and in elm genetic transformation (e.g. Gartland et al., 2000).

### Phylogenetic analysis

The initial assembly contained 12,213 prefiltered loci. Sequential filtering (max_SNPs) excluded 7,529 loci that exceeded the maximum number of SNPs, 12 additional loci that exceeded the maximum shared-heterozygosity threshold (max_shared_Hs_locus) and eight additional loci that failed the minimum sample-occupancy criterion (minimum-sample), resulting in a final alignment of 4,651 loci (38.1%). Most samples recovered approximately 3,700–4,000 loci, with mean depths typically around 22–30x. Our WGS dataset retained more loci per sample than the RADseq dataset (mean 3865.7 vs 3266.9; median 3944 vs 3378, respectively), and also recovered more consensus reads (mean 9920.3 vs 8074.5) and aligned sites (mean 803,842 vs 683,495). Mean depth across retained loci was similar between the WGS-derived and published RADseq datasets (25.62× and 25.82×, respectively). Concatenated local alignments for 176 taxa were 400,461 bp long.

The maximum likelihood phylogenetic analysis recovered all major clades within *Ulmus* previously identified in the RADseq phylogeny of Whittemore et al. (2021), with strong bootstrap support at most nodes (Fig. 3). The analysis recovered *Ulmus* as monophyletic and supported the recognition of three subgenera: *Ulmus* subg. *Oreoptelea*, *Ulmus* subg. *Indoptelea* and *Ulmus* subg. *Ulmus* (Whittemore et al. 2021) (Fig. 3). Our whole-genome sampling added several taxa not represented in the Whittemore et al. (2021) dataset, including *U. harbinensis, U. glaucescens, U. gaussenii, U. lancifolia, U. taihangshanensis, U. pinnato-racemosa, U. elliptica,* and *U. ismaelis* (Fig. 3). The two *U. chenmoui* accessions (E076 and E049) are not monophyletic. E076 may have been misidentified, but E049 is closely related to *U. chenmoui_ULMCHE-1* in Whittemore et al. (2021), which could be a correct identification. The *U. szechuanica* (E086) accession may also be misidentified based on our phylogenetic analysis, though the backbone of the tree is not well-supported. The IDs for *U. changii* (E048), *U. prunifolia* (E061), *U. chenmoui* (E049), *U. szechuanica* (E066), and *U. castaneifolia* (E047) appear correct. The *U. harbinensis* (E055) might actually be *U. lamellosa* based on Fig. 3b. The phylogenetic relationships of four species (*U. glaucescens, U. gaussenii, U. lancifolia,* and *U. taihangshanensis*) are difficult to confirm due to the lack of data in Whittemore et al. (2021), therefore multiple accessions are required to confirm their identification. Additionally, Whittemore et al. (2021) phylogenetic tree contains several misidentified species (e.g., *U. castaneifolia*, *U. davidiana*), which are not addressed here due to being out of scope.

The new phylogenetic tree reveals very short internal branches among the sampled *U. glabra* accessions from Europe, indicating low divergence in this dataset (Fig. 3). It also detects multiple cases of mislabelling in UK living collections, when compared with the Whittemore et al. (2021) reference accession, as also shown by the PCA analysis (see above) and ADMIXTURE analysis (Table S2). For example, E144 (labelled as *U. castaneifolia,* Kew, accession 1973-11726*1) is sister to *U. wallichiana* (E045, Kew), and both are closely related to three Whittemore accessions that are *U. wallichiana*. The tree also recovered *U. glabra* and *U. wallichiana* as closely related lineages, and *U. minor* and U. pumila as closely related lineages (Fig. 3). The phylogenetic relationships among the *U. davidiana* accessions are not resolved, which has been termed the *U. davidiana* complex by Whittemore et al. (2021). The network relationships of the same data matrix are shown in Fig. S7.

### Admixture analysis

In presenting ADMIXTURE results, we note that our sampling is not ideally suited to the software as it assumes random sampling of populations and may be sensitive to uneven sampling and hierarchical population structure (Puechmaille, 2016). These results might reflect the sensitivity of clustering algorithms to uneven lineage representation and the complex hybrid backgrounds present in some elm samples. We therefore interpret ancestry components as summaries of genomic affinity rather than as exact reconstructions of pedigree, particularly for cultivars and complex hybrids with multigenerational breeding histories.

The ADMIXTURE analysis of the Elms-120 dataset (i.e. mainly natural species and elms not artificially crossed - see Methods), based on 923,700 SNPs had a best-supported value of K=5, which identified the following groups: (1) several American species and *U. villosa,* (2) *U. laevis*, (3) Asiatic species, (4) *U. glabra*, and (5) *U. minor* (Fig. 4a). In this analysis several natural species from East Asia, and also *U. americana* appear to contain allele frequencies suggestive of several groups and do not form a group of their own. This is probably due to insufficient sampling of these species. Similarly, *U. wallichiana* (E045) from the Himalaya, appears to incorporate alleles suggestive of all five groups in K=5, having a majority from the Asiatic cluster (52%), but also a large *U. glabra* component (30%). *Ulmus elliptica* (E051) from the Caucasus seems to be mainly *U. glabra* (87%) with some allele frequency shared with Asiatic species (10%) and the Himalayan *U. villosa*.

**Fig. 4.**
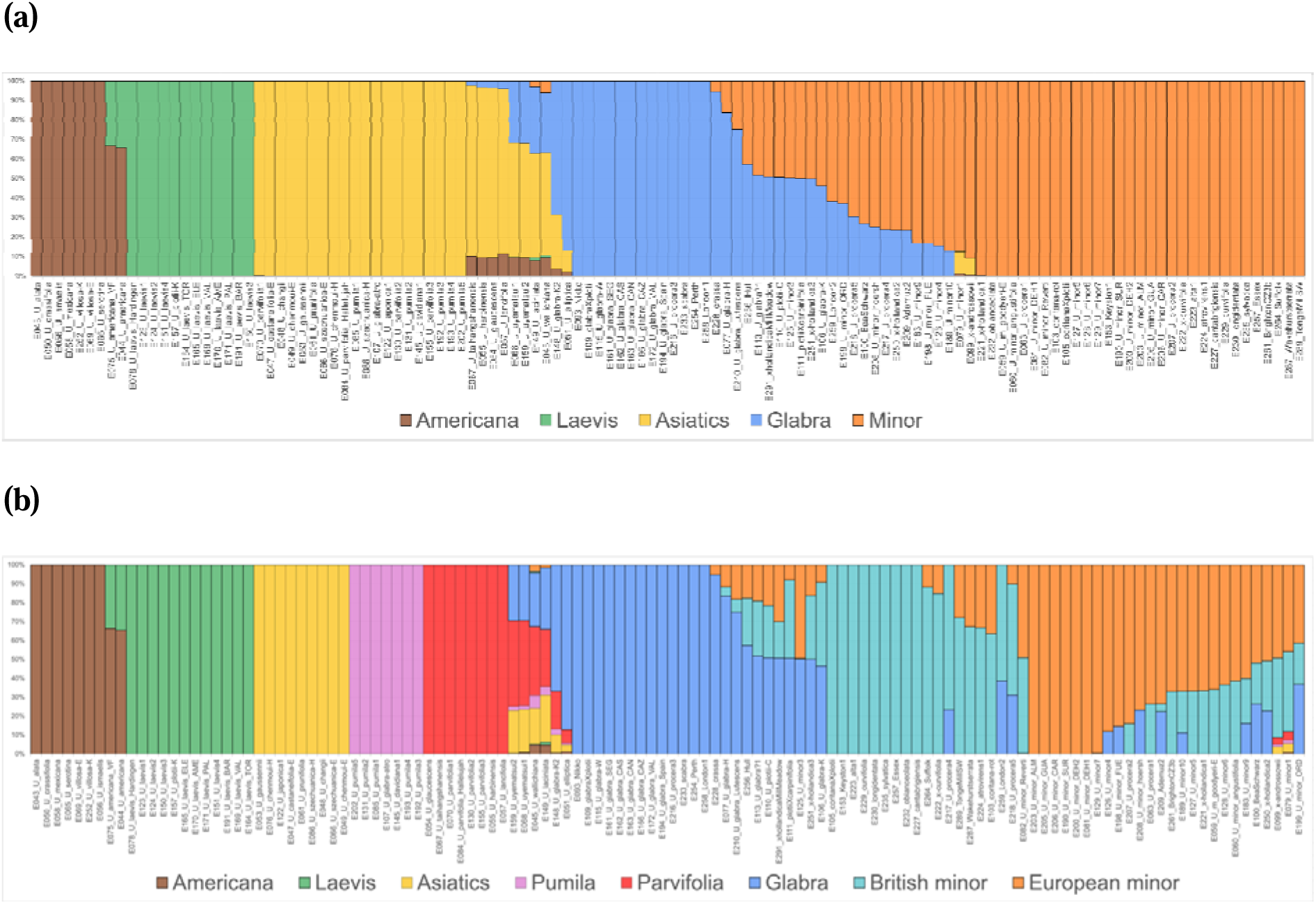
ADMIXTURE analysis of the Elms-120 dataset, comprising mainly natural species and accessions not identified as complex artificial hybrids. (a) K=5 (best K), the colours can be interpreted as representing the following gene pools from left to right: Brown: Several American species and *U. villosa*, Green: *U. laevis*, Yellow: Asiatic species, Blue: *U. glabra*, and Orange: *U. minor.* (b) K=8, the colours can be interpreted as representing the following gene pools: Brown: Several American species and *U. villosa*, Green: *U. laevis*, Yellow: Asiatic species including *U. japonica*, Purple: *U. pumila*, Red: *U. parvifolia*, Blue: *U. glabra*, Cyan: British *U. minor*, Orange: European *U. minor*. Analyses with other values of K, and a best cross-validation plot are shown in Fig. S4.

An Asiatic component (27%) is detected in a *U. glabra* accession from Kew (E148), an *U. minor* (10% Asiatic) from Sir Harold Hillier Gardens (E079) and *U.* × *androssowii* (8.6% Asiatic) from Sir Harold Hillier Gardens (E099). This signal was consistently found from K=3 to K=10, suggesting some exotic admixture in these samples. The *U.* × *androssowii* accession originated from Uzbekistan and was suspected to be an old hybrid of unknown origin between *U. minor* × *U. pumila*. Our results are consistent with this but suggest that the Asiatic component is small and 90% of its genome comprises *U. minor* (Fig. 4a).

From K=8, Asiatic species clusters were split into three groups, and the *U. minor* group was split into a mainly European *U. minor* and a mainly British *U. minor* cluster, with a great deal of admixture between them. “Pure” examples of the European *U. minor* cluster were E200 *U. minor* DEH2 (Dehesa de la Villa), E081 *U. minor* DEH1 (Dehesa de la Villa), E206 *U. minor* CAR (Villanueva Del Carazo), E190 *U. minor* SUR, E203 *U. minor* ALM (Almarcha) and E205 *U. minor* GUA (Guadalajara). “Pure” examples of the British minor group were E105 *U. coritana* × *plotii*, E153 *U. minor* ‘Keyston’, E223 *U. alta*, E230 *U. longidentata*, E235 *U. sylvatica*, E229 *U. curvifolia* and E257 *U. minor* (from Essex), several of which are Sell and Murrell (2018) taxa. Many samples of *Ulmus* from the British landscape contain components of both British and European *U. minor* groups, with *U. procera* falling largely within the European group; and some also contain an *U. glabra* component (Fig. 4b). Analyses with other values of K, and the best K plot are shown in Fig. S4.

ADMIXTURE analysis of the Elms-132 dataset, comprising the Elms-120 accessions plus one representative from each selected clonal group of complex hybrids, was performed using 645,489 LD-pruned SNPs. Cross-validation was lowest at K=6 (CV = 0.40562), and therefore we interpret K=6 as the best-supported results, while K=8 provides complementary results that resolve Asian-derived ancestry. At K=6, ancestry components correspond to American/*U. parvifolia* (K1), *U. glabra* (K2), *U. minor* (K3), *U. laevis* (K4), *U. pumila*/*U. davidiana* (K5), and Asiatics (K6). At K=8, the *U. minor* cluster is further resolved into British *U. minor* (K4) and European *U. minor* (K8), similar to the Elms-120 dataset (Fig. 4b), while *U. laevis* (K1), *U. pumila*/*U. davidiana* (K2), *U. parvifolia* (K3), Asiatics (K5), *U. glabra* (K6), and American/*U. villosa* (K7) are recovered separately. The Spanish DED-tolerant clones (Martín *et al*., 2015) ranged from essentially pure *U. minor* genotypes (‘Retiro’; ‘Dehesa de la Villa’) to *U. glabra* admixed accessions (‘Dehesa de Amaniel’, and ‘Ademuz’) (Table 2). ‘Ademuz’ (E132), an elm with currently promising arboricultural and resistance properties (Martín *et al*., 2015), showed mainly *U. minor* ancestry (76.3%) but also, somewhat unexpectedly, significant *U. glabra* ancestry (23.6%). Retiro (E129) was nearly pure minor (98.7%) with only a small *U. glabra* proportion (1.3%), whereas E199 (*U. minor*, Ordes, Spain) was more admixed (K3=62.8%, K2=37.2%). E079 (labelled *U. minor*) and E082 (*U. minor* Reverti) were largely *U. minor*, while E125 (*U. minor* ‘Christine Buisman’, originally from Spain) was almost evenly split between *U. minor* (49.8%) and *U. glabra* (50.2%) ancestry, consistent with hybrid or *U. × hollandica* affinity. The Tonge Mill elm (E288) was recovered as essentially pure *U. minor*.

**Table 2.**
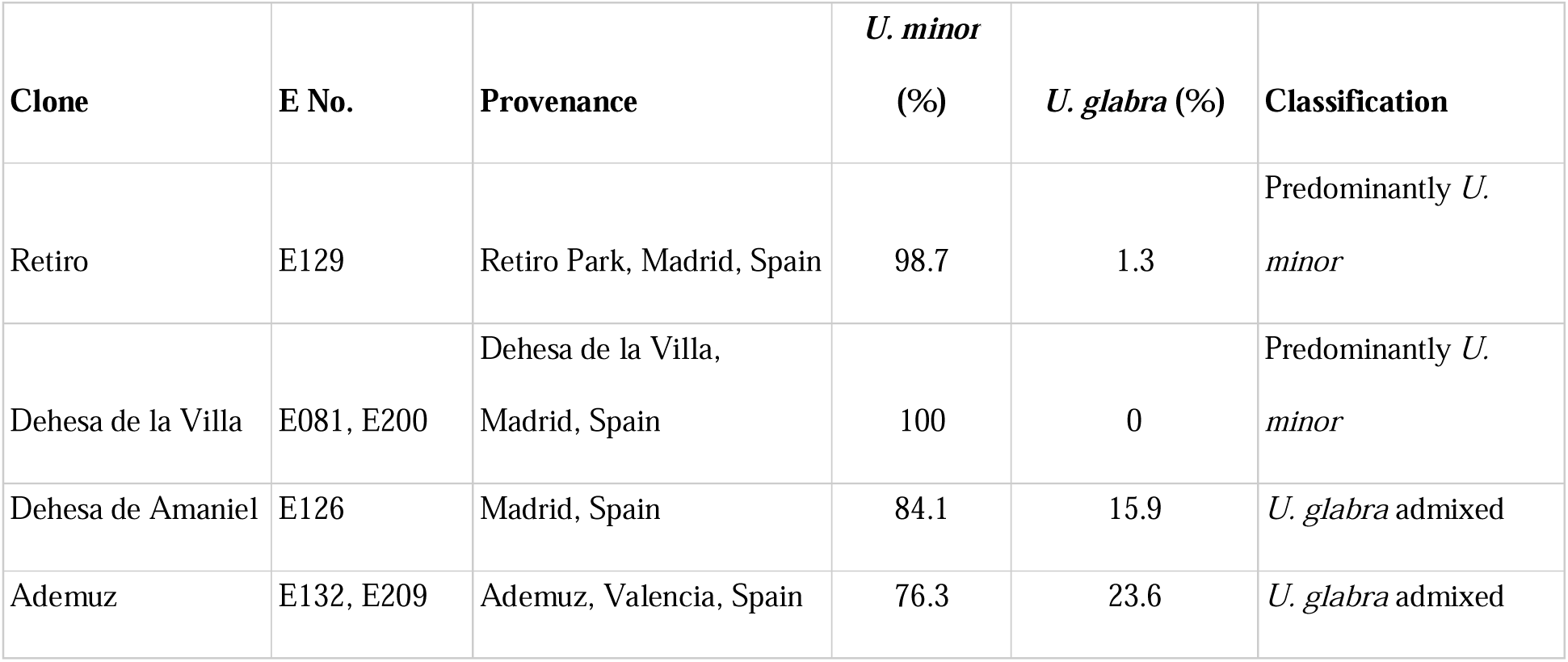
ADMIXTURE ancestry of Spanish DED-tolerant clones (Martín *et al*., 2015) derived from Elms-132, K=6 (Fig. S5).

**Table 3.** Hybrid cultivar parentage based on ADMIXTURE K=8 ancestry of Elms-132 dataset.

| E No. | Hybrid / cultivar | Recorded parentage | 'European'<br>*U. minor*<br>(K8) | 'British'<br>*U. minor*<br>(K4) | *U. glabra*<br>(K6) | Asiatics<br>(K5) | *U. pumila*<br>(K2) | *U. parvifolia*<br>(K3) | *U. laevis*<br>(K1) | American/<br>*U. villosa*<br>(K7) |
| --- | --- | --- | --- | --- | --- | --- | --- | --- | --- | --- |
| E087 | 'Amsterdam' | 'Bea Schwarz' × *U. minor* | 65.6 | 16.3 | 18.1 | 0 | 0 | 0 | 0 | 0 |
| E100 | 'Bea Schwarz' | *U. minor* derived clone | 64.8 | 9.8 | 25.5 | 0 | 0 | 0 | 0 | 0 |
| E251 | 'Wentworthi' | *U. × hollandica* cultivar | 18 | 32.5 | 49.5 | 0 | 0 | 0 | 0 | 0 |
| E133 | 'Commelin' | *U. × hollandica* 'Vegeta' × *U. minor* | 31.1 | 39.2 | 29.7 | 0 | 0 | 0 | 0 | 0 |
| E221 | 'Wredei' | *U. × hollandica* (*U. glabra* × *U. minor*) | 67.4 | 32.6 | 0 | 0 | 0 | 0 | 0 | 0 |
| E094 | 'Plantijn'/'Plantyn' | (*Exoniensis* × *U. wallichiana*) × (*U. minor* × *U. minor*) | 36 | 0 | 57.1 | 5.8 | 0 | 0.3 | 0 | 0.8 |
| E119 | 'Lutèce'/'Nanguen' | 'Plantyn' × ('Bea Schwarz' selfed) | 60.4 | 0 | 39.6 | 0 | 0 | 0 | 0 | 0 |
| E120 | 'Vada'/'Wanoux' | 'Plantyn' × 'Plantyn' selfed | 55.8 | 0 | 41.4 | 2.8 | 0 | 0 | 0 | 0 |
| E089 | 'Dodoens' | (*Exoniensis* × *U. wallichiana*) selfed | 17 | 20.6 | 55.3 | 5.5 | 0 | 0.6 | 0 | 0.9 |
| E158 | 'San Zanobi' | 'Plantyn' × *U. pumila* | 44.8 | 0 | 39.4 | 11.8 | 0 | 1.6 | 0 | 2.4 |
| E154 | 'New Horizon' | *U. davidiana* var. *japonica* × *U. pumila* | 0 | 0 | 0 | 48.8 | 51.2 | 0 | 0 | 0 |
| E118 | 'Morfeo' | (*U. × hollandica* × *U. minor*) × *U. chenmoui* | 27.3 | 0 | 22.2 | 50.5 | 0 | 0 | 0 | 0 |
| E090 | 'Fiorente' | *U. pumila* × *U. minor* | 31.3 | 19.2 | 0 | 0 | 49.5 | 0 | 0 | 0 |
| E099 | x-androssowii | often treated/suspected as *U. minor* × *U. pumila* | 49.1 | 41.4 | 0 | 5.7 | 1.1 | 0.9 | 0.1 | 1.6 |

Among complex hybrids, the Dutch cultivars (Heybroek, 1993a) were dominated by the *U. minor* and *U. glabra* clusters. ‘Plantyn’ (E094) showed 57.1% *U. glabra*, 33.0% *U. minor,* 4.4% Asiatics, and 5.5% American/*U. parvifolia* ancestry, while ‘Dodoens’ (E089) was 55.0% *U. glabra*, 36.9% *U. minor*, 3.7% Asiatics, and 4.4% American/*U. parvifolia* ancestry. ‘Lutèce’/’Nanguen’ (E119) and ‘Wanoux’/’Vada’ (E120), both derived from ‘Plantyn’, were composed mainly of *U. minor* and *U. glabra* ancestry, with ‘Lutèce’ containing 56.9% *U. minor* and 40.0% *U. glabra*, and ‘Wanoux’ 53.1% minor and 41.5% *U. glabra*, respectively. Amsterdam (E087) was mostly *U. minor* (80.6%) with 19.4% *U. glabra* ancestry, whereas ‘Commelin’ (E133) showed 69.7% *U. minor* and 30.3% *U. glabra* ancestry. Among the more clearly Asian-derived cultivars, ‘Morfeo’ (E118) differed from the Dutch hybrids in carrying a strong Asiatic component (50%), together with 26.6% *U. minor* and 23.4% *U. glabra* ancestry. FL493/’Wingham’ (E041) comprised 49.3% *U. minor*, 30.7% *U. glabra*, and 15.5% *U. pumila*/*U. davidiana* ancestry. *Ulmus* ‘San Zanobi’ (E158) which is hybrid of *Ulmus* ‘Plantyn’ and *U. pumila* was mainly *U. minor* (43.0%) and *U. glabra* (39.6%) with only 9.3% Asiatics, but this accession may be mislabelled as according to our Kinship analysis (see above) Ulmus ‘San Zanobi’ (E158) is identical to the Dutch hybrid ‘Lobel’ (E135). As expected, ‘New Horizon’ (E154) was recovered as approximately equal *U. pumila/U. davidiana* (51.6%) and Asiatics (48.4%) ancestry, and ‘Fiorente’ (E090) showed an approximately equal split between *U. minor* (50.4%) and *U. pumila/U. davidiana* (49.6%) ancestry. *Ulmus wallichiana* (E045) appears to contain all six groups in K=6, American/parvifolia/villosa (23.18%), *U. glabra* (34.6%), *U. minor* (3.12%), *U. laevis* (1.86%), *U. pumila/U. davidiana* (5.36%), and Asiatics (31.85%), probably due to incomplete lineage sorting. In comparing ADMIXTURE analyses, we found that occasional ancestry assignments varied depending on the individuals included in the analysis. For example, in the Elms-120 analysis, ADMIXTURE inferred Asiatic ancestry in the European *U. minor* ‘Ademuz’ (E132), at the K=6. However, when this sample was replaced with another clone of ‘Ademuz’ (E209), the Asiatic ancestry was not detected in the best model (K=5) (data not shown). The ADMIXTURE analysis of the Elms-132 dataset did not show this Asiatic component either for E132. Small components of unexpected ancestry that appear in some analyses and not others may be due to rare genetic variation that the algorithm struggles to classify.

At K=8, the previously merged *U. minor* cluster was divided into European and British *U. minor* as in the Elms-120 data of K=8, and the separation of the Asiatic species (Fig. 5b). Among the Dutch hybrids, the ‘Plantyn’ (E094) comprised 57.1% *U. glabra*, 36.0% European *U. minor* and 5.8% Asiatics, ‘Dodoens’ (E089) 55.3% *U. glabra*, 20.6% British *U. minor*, 17.0% European *U. minor* and 5.5% Asiatics, ‘Lutèce’/’Nanguen’ (E119) 60.4% European *U. minor* and 39.6% *U. glabra*, and ‘Wanoux’/’Vada’ (E120) 55.8% European *U. minor*, 41.4% *U. glabra* and 2.8% Asiatics. New Horizon (E154) was recovered as approximately equal *U. pumila* (51.2%) and Asiatics (48.8%) ancestry, ‘Fiorente’ (E090) as 49.5% *U. pumila*, 31.3% European *U. minor* and 19.2% British *U. minor*, and ‘Morfeo’ (E118) as 50.5% Asiatics, 27.3% European *U. minor* and 22.2% *U. glabra*. By contrast, ‘San Zanobi’ (E158) comprised European *U. minor* (44.8%) ancestry and *U. glabra* (39.4%), with only 11.8% Asiatics. The clusters above K=8 resolved the Dutch hybrids into a cluster of their own, giving no information about their hybrid origin (Fig. S5).

**Fig. 5.**
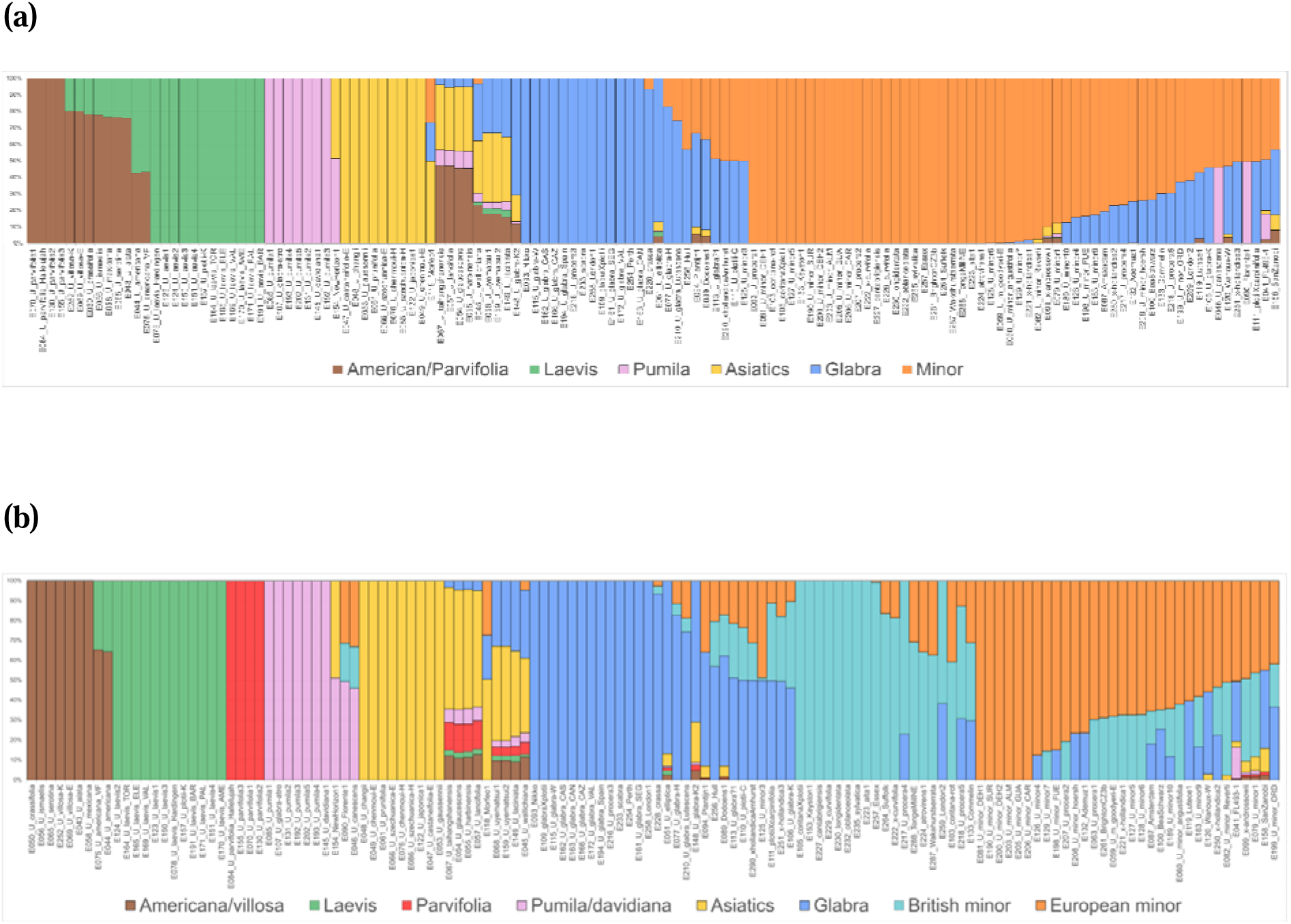
ADMIXTURE analysis of 132 elm accessions, comprising the Elms-120 dataset plus one representative from each selected clonal group of complex hybrids. (a) K=6 (best K), showing the six inferred ancestry components: Brown: American/*U. parvifolia*, Green: *U. laevis*, Purple: *U. pumila/U. davidiana*, Yellow: Asiatics, Blue: *U. glabra*, and Orange: *U. minor*. (b) showing K=8 inferred ancestry components: Brown: American/ *U. villosa*, Green: *U. laevis*, Red: *U. parvifolia*, Purple: *U. pumila/U. davidiana*, Yellow: Asiatics, Blue: *U. glabra*, Cyan: ‘British’ *U. minor*, and Orange: ‘European’ *U. minor*. Samples are ordered by the number of ancestry components they contain. Analyses at other values of K and cross-validation results are shown in Fig. S5.

Allele-sharing tests of AdmixTools 2 analysis supported the observation that the two sequenced ‘Ademuz’ samples are composed of ca. 75–76% European *U. minor* and 24–25% *U. glabra*, and in both samples the European *U. minor*-only model was rejected, whereas the full two-source model (*U. minor* and *U. glabra*) fitted well. By contrast, the putatively pure *U. minor* (E081, E129 and E200) did not require a *U. glabra* source, and E126 remained European *U. minor* alone (Table S3).

### Genome-wide patterns of diversity within clusters

A balanced comparison of 14 individuals per species showed higher genomic diversity in *U. minor* than in *U. glabra* (Fig. 6). Mean observed heterozygosity was 0.114 in *U. minor* and 0.093 in *U. glabra* (Fig. 6a), while mean unbiased expected heterozygosity was 0.171 and 0.148, respectively (Fig. 6b). We obtained a consistent pattern of *Ho* and *He* across 14 chromosomes (Fig. 6c) and genomic windows with *U. minor* showing higher *Ho* in 82.4% of paired 1-Mb windows (Fig. 6d). A complementary nucleotide diversity analysis also supported this pattern, with higher mean π in *U. minor* than in *U. glabra* (0.00133 vs 0.00115; 81.7% of windows higher in *U. minor*) (Fig. S8). Because this estimate was based on a SNP-only VCF, it was interpreted as a relative comparison rather than an absolute genome-wide estimate of nucleotide diversity. The levels of genetic diversity within some taxa are shown by SNP density plots in different groups of sequenced individuals in Fig. S9. The SNPs were broadly distributed across the genome in all groups. The 14 *U. minor* individuals contained 7,018,578 SNPs, approximately 0.53 million more than the 6,485,481 SNPs detected in the 14 *U. glabra* individuals (Fig. S9).

**Fig. 6.**
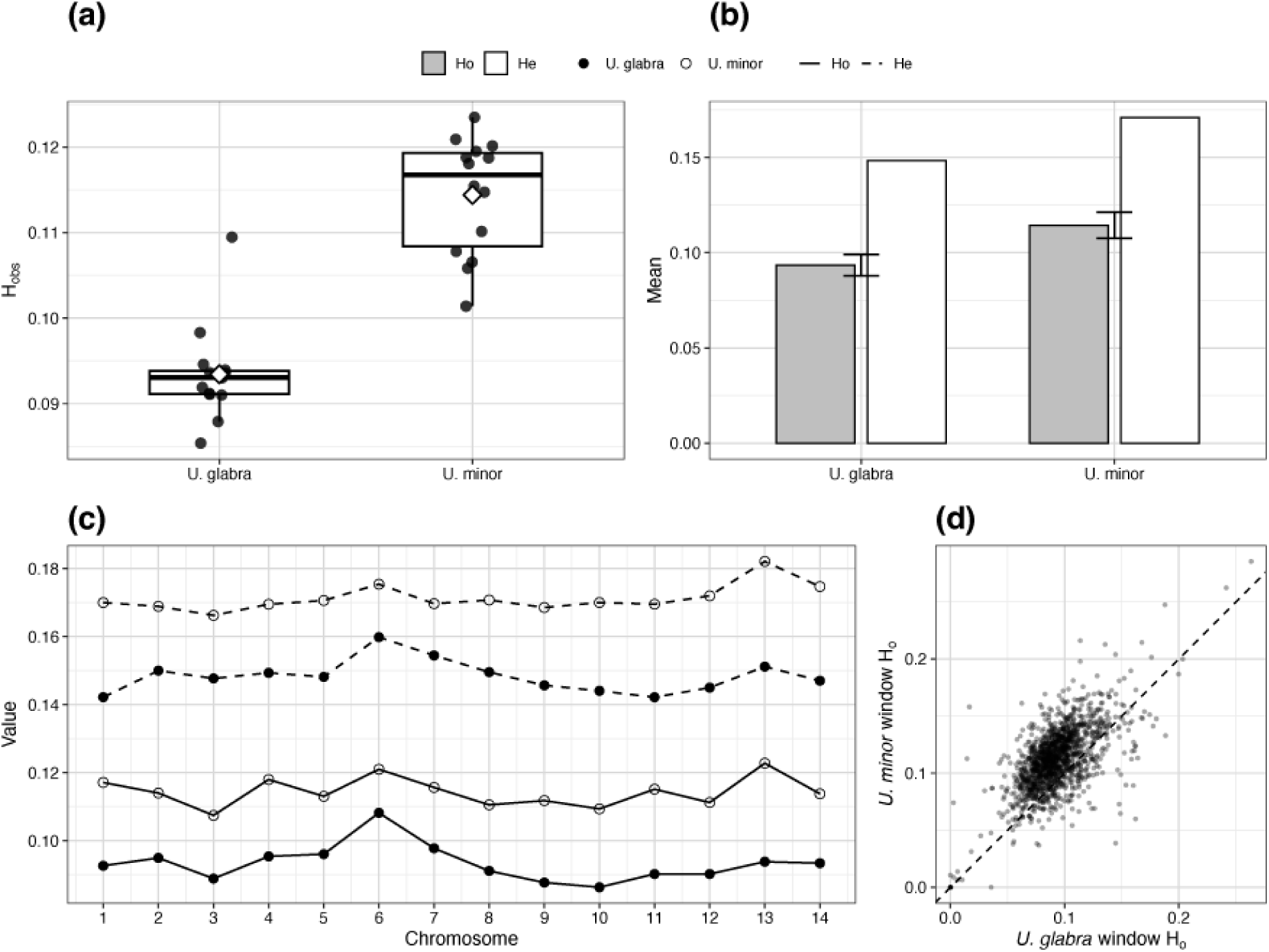
(a) Individual observed heterozygosity (*Ho*) for 14 *U. glabra* and 14 *U. minor* individuals (eight British and six Spanish individuals of each species). Points represent individuals and diamonds indicate group means. (b) Mean observed heterozygosity (*Ho*) and unbiased expected heterozygosity (*He*) are shown for each species. Grey bars show *Ho*, white bars show *He*; error bars indicate the standard deviation of individual *Ho*. (c) Chromosome-level *Ho* and *He* across the 14 elm chromosomes. Line type distinguishes *Ho* and *He*, and point symbols distinguish species. (d) Paired 1-Mb window comparison of observed heterozygosity, with *U. glabra* on the x-axis and *U. minor* on the y-axis. The dashed diagonal indicates equal *Ho* between species; points above the line indicate windows with higher *Ho* in *U. minor*.

## Discussion

In this study, we have conducted an extensive genome-wide survey of elm trees growing in Britain, as well as other trees currently grown in Europe that could be planted in Britain in the future. As expected, we found extensive genomic diversity in many artificial cultivars, spanning species-level differences, compared with pure *U. minor*, *U. glabra*, and *U. pumila*. Our findings also confirm the accuracy of many tree labels and the recorded parentage for many artificial cultivars.

### Genomic diversity of elms

Based on genome-wide observed heterozygosity and nucleotide diversity, we find *U. minor* to be more diverse than *U. glabra* (Figs 6, S8 and S9). This is not surprising given the morphological variability observed in *U. minor* in Britain that led to the discrimination of numerous microspecies by Sell and Murrell (2018). This difference between the two species may be partly because *U. glabra* largely reproduces sexually (Thomas *et al*., 2018), meaning that variation will be gradually lost in small populations, whereas *U. minor* is known for clonal propagation (Cox *et al*., 2014) which maintains heterozygosity even in small populations. The high morphological diversity of *U. minor* may also reflect its adaptation to a heterogeneous array of habitats across Eurasia, in comparison to *U. glabra* which is predominantly a temperate, moist woodland tree (Thomas *et al*., 2018). Our ADMIXTURE analyses found two *U. minor* clusters (European and British), with extensive introgression between them. This pattern may reflect a geographic cline of genetic diversity within *U. minor* further mixed by anthropogenic movement of clones, some of which are believed to date back to the Roman Empire or even earlier. We also obtained further evidence for extensive hybridisation and backcrossing between *U. minor* and *U. glabra. S*ome examples of these had collection labels suggesting they were pure *U. minor* or pure *U. glabra* and would be hard to discriminate without genome-wide analysis (Figs 2 and 4).

Our PCA and ADMIXTURE results illustrate that artificial hybrids occupy a genomic space that spans and often complements that of natural species and their natural hybrids (Figs 1, 2, 4 and 5). It shows how since the 1970s, following the second pandemic of DED, artificial hybridisation through elm breeding programmes introduced novel genomic diversity into native European and North American elms from resistant Asian species (Heybroek, 1993a; Mittempergher & Santini, 2004; Martín *et al*., 2023). This expanded the available genomic diversity in breeding clones, as has also been seen in other tree species (Janes & Hamilton, 2017; Yan *et al*., 2024). Many of the cultivars analysed here represent a reservoir of often exotic genomic diversity, assembled with DED-resistance in mind. In one sense this significantly enhances the genomic diversity of elms in Britain today, since it takes it beyond the diversity still surviving in the post-DED native elm landscape. On the other hand, the number of planted individuals of these cultivars is currently insignificant compared to the millions of regenerating but currently early-dying *U. minor, U. glabra* and their hybrids.

Limited planting of a more genetically diverse elm-scape, especially for urban street planting, ornamental planting and other specialised uses, is already becoming a reality. However, in terms of planting at a landscape and woodland scale, the properties and suitability of most of these cultivars have yet to be critically evaluated, notably their disease resistance under UK conditions and their arboricultural and ecological suitability. The latter applies especially to cultivars with a significant proportion of Asiatic parentage. Furthermore, the consequences for UK natural habitats and biodiversity of any landscape-wide planting, especially in the context of a very large, surviving though frequently reinfected, but otherwise highly adapted endemic elm population, also need to be evaluated.

We provide strong genomic evidence that the 21 accessions included in our Sell and Murrell comparison, including accessions representing 11 named species or forms (Sell & Murrell, 2018), fall within the genomic variation sampled for *U. minor*, *U. glabra* or their hybrids (Figs 1-5 and S6). Our admixture analysis also suggests that *U. carpinifolia* and two individuals of the *U. coritana* of Sell and Murrell are a hybrid between *U. minor* and *U. glabra* (Fig. 4). The genome-wide *F_ST_* results (Methods S1) also confirm very little genetic differentiation between Sell and Murrell ‘species’ and *U. minor* than between *U. minor* and either *U. glabra* or *U. laevis* (Fig. S6). Other Sell and Murrell species, such as *U. alta, U. atrovirens, U. cantabrigiensis, U. longidentata,* and *U. sylvatica*, are also *U. minor* based on the PCA and ADMIXTURE analysis (Figs 4 and 5). Most probably they are mainly human-cultivated clones. Further consideration of whether or not these taxa deserve recognition at an infraspecific ranking level, such as microspecies, is beyond the scope of the present study. Our genomic results, together with the morphological diversity found within *U. minor* by the proponents of these microspecies, again suggests that *U. minor* has a long history in Britain (cf. Richens 1983).

We identify some cases of mislabelling in living collections. This is particularly common in the Melville collection, preserved as a laid hedge at Wakehurst, UK, perhaps because different species are entangled or because labels have been misplaced during hedge management. Mislabelling is common in biological collections (Goodwin *et al*., 2015), and careful genetic-based analysis is required to validate the ancestry of the lineage in hand (Migicovsky *et al*., 2019).

Our study also sheds light on the ancestry of some of the known natural and artificial hybrids using genome sequences. For example, ‘Bea Schwarz’ was an early selection made in France after the first DED pandemic and used in many artificial crosses (Santini *et al*., 2004). Its ancestry had long been debated (Santamour & Bentz, 1995), with some favouring *U. minor* and others *U. × hollandica*. Our K=5 analysis suggests it is about a quarter *U. glabra* and three quarters *U. minor*, and the K=8 analysis adds information suggesting that the *U. minor* component contains both ‘European’ and ‘British’ lineages. It therefore appears to be a cross between a pure ‘European’ *U. minor* and a *U. × hollandica* formed from a hybridisation between a ‘British’ *U. minor* and a *U. glabra* (Cox *et al*., 2014).

### Implications of the current study for future elm planting

Our discovery that two DED-tolerant Spanish selections previously considered *U. minor* are in fact admixed with 16–25% *U. glabra* ancestry (Table 2) was unexpected, but could be explained by the coexistence of these two species in glacial refugia in Spain (Gil *et al*., 2004) or subsequent natural hybridisation. This finding has practical implications for those who wish to plant only pure species. It also demonstrates the benefits of genomic verification of accessions. The possibility of a positive association between *U. glabra* introgression and tolerance to *O. novo-ulmi* among the Spanish selections (e.g. ‘Ademuz’), also indicated by early Dutch hybrid releases tested with *O. ulmi* (e.g., ‘Christine Buisman’ and ‘Commelin’), warrants further investigation. If transgressive segregation, or specific *U. glabra* alleles, contribute to tolerance, this could inform future crosses.

As mentioned above, our data do not support the recognition of several of the Sell and Murrell (2018) elm species. For conservation purposes, these forms should be treated as part of the natural variation of *U. minor* and *U. glabra* and their hybrids in Britain rather than as species of individual conservation concern. In our view they would be inappropriate for red listing. However, the genetic diversity within them undoubtedly merits conservation as a resource for restoration (West *et al*., 2025).

Although in recent years multiple reference genomes and annotations have been published for elms which were not available at the time this research started (Lyu *et al*., 2025; Coleman & Ruhsam, 2024; Wang *et al*., 2025; Pallares Zazo *et al*., 2025), genomic resources for elms are still limited, such that there is no genome annotation for *U. glabra* and other ancestral species of various hybrids (i.e. *U. wallichiana*, *U. japonica* and *U. pumila*). Functionally annotated elm genomes are necessary to track genomic-scale diversity, admixture, and traits related to Dutch elm disease resistance. This study is the first attempt to study genomic signatures of numerous widely known elm cultivars and hybrids. Nevertheless, we have sequenced fewer than half of the more than 100 known elm cultivars/hybrids worldwide. Therefore, we recommend further effort to sample as many of these cultivars as possible from living collections to increase the genomic diversity panel in future studies.

## Acknowledgements

This project was funded by the Department for Environment, Food & Rural Affairs (Defra) via the Centre for Forest Protection (Ref. 2967). We are grateful to Defra staff for their support and interest. We are also thankful to the Royal Botanic Gardens, Kew, and the staff at Sir Harold Hillier Gardens for their help and cooperation during sampling. We thank Matthew Charles Ellis from Grange Farm Arboretum, Nick Dunn from Frank P. Matthews Ltd, Barry Clarke from Sir Harold Hillier Gardens, Gemma Hampshire, Cicely Marshall, Alex Prendergast, Alberto Santini, Christopher P Cockel, and Chris Poynton for providing materials for sequencing. We are also grateful for the knowledge-sharing of Allison Oakes, David Shreeve from the Conservation Foundation, David Macaya Sanz, and Andrew Brookes from Butterfly Conservation. Special thanks to Laszlo Csiba, Robyn S Cowan at the Jodrell laboratory and the Plant Health and Adaptation team at Kew for valuable advice on this study. We thank Max Coleman for helpful conversations and comments on an earlier draft of this paper. The authors acknowledge resources and technical support of Queen Mary University of London, Apocrita cluster (https://doi.org/10.5281/zenodo.438045) and the UK’s Crop Diversity Bioinformatics cluster of the James Hutton Institute (https://doi.org/10.1002/ppp3.10607).

## Author contributions

Mohammad Vatanparast co-designed the project, collated the samples and conducted laboratory and bioinformatic analyses and co-wrote the manuscript, Joan F. Webber and Clive Brasier supplied samples and contributed to the manuscript, Juan A. Martín supplied samples and contributed to the manuscript, Richard Buggs supervised the project, raised funds, co-designed the project, co-wrote the manuscript.

## Data availability

The raw sequences generated in this project are available from GenBank under project number PRJNA982084. Scripts used to analyse data are available from https://github.com/vatanparast/Genomic_Diversity_Elms

