## Supplementary Material for "Genomic diversity of elm trees for future treescapes"

### **List of Supporting Information**

**Methods S1** Genome-wide differentiation analysis.

**Table S1** Sample metadata and sequencing quality statistics.

**Table S2** Summary of apparent mislabelling and unexpected genomic identifications.

**Table S3** qpAdm and ADMIXTURE analyses of Spanish DED-tolerant clones.

**Fig. S1** Photographs of selected sequenced elm trees.

**Fig. S2** Principal components 3–10 for all 204 sequenced elm accessions.

**Fig. S3** Principal components 3–10 for the 180 accessions of *Ulmus* subgenus *Ulmus*.

**Fig. S4** Best K plot of the ADMIXTURE analysis of 120 elms and different K plots.

**Fig. S5** Best K plot of the ADMIXTURE analysis of 132 elms and different K plots.

**Fig. S6** Genome-wide differentiation (*F_ST_*).

**Fig. S7** NeighborNet phylogenetic network.

**Fig. S8** Nucleotide diversity (𝜋) between *U. minor* and *U. glabra*.

**Fig. S9** Genome-wide SNP-density distributions in selected elm groups.

**Methods S1.** Genome-wide differentiation analysis.

#### Genome-wide differentiation (*F_ST_*)- methods

To compare genome-wide differentiation between *U. minor* and other sampled elm groups, we estimated Weir and Cockerham’s *F_ST_* (Weir & Cockerham, 1984) for three comparisons: *U. minor* versus *U. laevis*, *U. minor* versus *U. glabra*, and *U. minor* versus accessions bearing taxon names recognized by Sell and Murrell (2018). We included 30 samples of *U. minor* and 13 samples of *U. laevis*; 30 samples of *U. minor* and 15 samples of *U. glabra*; 30 samples of *U. minor* and 21 samples of Sell and Murrell taxa (*U. alta, U. atrovirens, U. cantabrigiensis, U. carpinifolia, U. coritana, U. crassa, U. curvifolia, U. longidentata, U. oblanceolata, U. procera, U. scabra, U. serratifrons, U. sylvatica, U.* × *curvifolia*). The *F_ST_* values across chromosomes were plotted using the qqman R package (Turner, 2018).

##### Genome-wide differentiation (*F_ST_*) - results

Genetic differentiation between *U. minor* and *U. laevis* is high (*F_ST_*=1.0) throughout the genome (Fig. S6a). The *F_ST_* is nearly equal to 0.5 between *U. minor* and *U. glabra*, with Chromosome 4 showing slightly higher differentiation (Fig. S6b). Genetic differentiation between the pooled Sell and Murrell accessions and the sampled *U. minor* accessions was substantially lower than that observed between *U. minor* and either *U. glabra* or *U. laevis* (*F_ST_* <0.2), showing that these are not well differentiated lineages and can be considered to fall within *U. minor* (Fig. S6c). This result is consistent with the PCA and ADMIXTURE analyses (Figs 4 and 5), although the pooled comparison does not assess differentiation among individual named forms.

### **Table S1** Sample identities, provenance, collection information, taxonomic assignments, PCA clusters and sequencing-quality statistics for the 204 elm accessions used in this study. Provided as a separate CSV file.

#### **Table S2** Summary of mislabelling and unexpected identifications.

| **Accession** | **Collection label** | **Provenance** | **Genomic identity** |
| --- | --- | --- | --- |
| E158 | ‘San Zanobi' | Kew, UK (Arboretum Nursery) | Clone-identical to ‘Lobel’; inconsistent with the recorded parentage ‘Plantyn’ × *U. pumila*. |
| E096 | ‘Rebona' | Sir Harold Hillier Gardens, UK (CK013) | ‘New Horizon' clone |
| E157 | *U. plotii* | Kew, UK (2010-1842*1) | *U. laevis* |
| E144 | *U. castaneifolia* | Kew, UK (1973-11726*1) | Clusters with *U. wallichiana* |
| E107 | *U. glabra* 'Atropurpurea' | Wakehurst, UK (014-D6.01411) | Clusters with *U. pumila* |
| E046 | *U. canescens* | Grange Farm, UK (ACC826) | *U. minor* *× U. pumila* hybrid (~50:50) |
| E148 | *U. glabra* | Kew, UK (1959-23702*1) | Approximately 27% affinity to the ADMIXTURE component predominant in sampled Asiatic accessions |
| E108 | *U. thomasii* (N. American) | Wakehurst, UK (486-68.48603) | 100% European *U. minor* in all analyses |
| E145 | *U. davidiana* | Kew, UK (1995-1305*1) | Groups with pumila cluster |

#### **Table S3** Summary of qpAdm and ADMIXTURE analyses for Spanish clones.

| **Clone** | **Sample(s)** | **ADMIXTURE *U. glabra* %** | **qpAdm *U. glabra* weight** |
| --- | --- | --- | --- |
| Dehesa de la Villa | E081, E200 | 0.0 | 7.7 / 13.1 |
| Retiro | E129 | 1.3 | 13.2 |
| Dehesa de Amaniel | E126 | 15.9 | 3.5 |
| Ademuz | E132, E209 | 25.0 | 25.3 / 24.0 |

#### **Fig. S1** Photographs of selected sequenced elm trees.

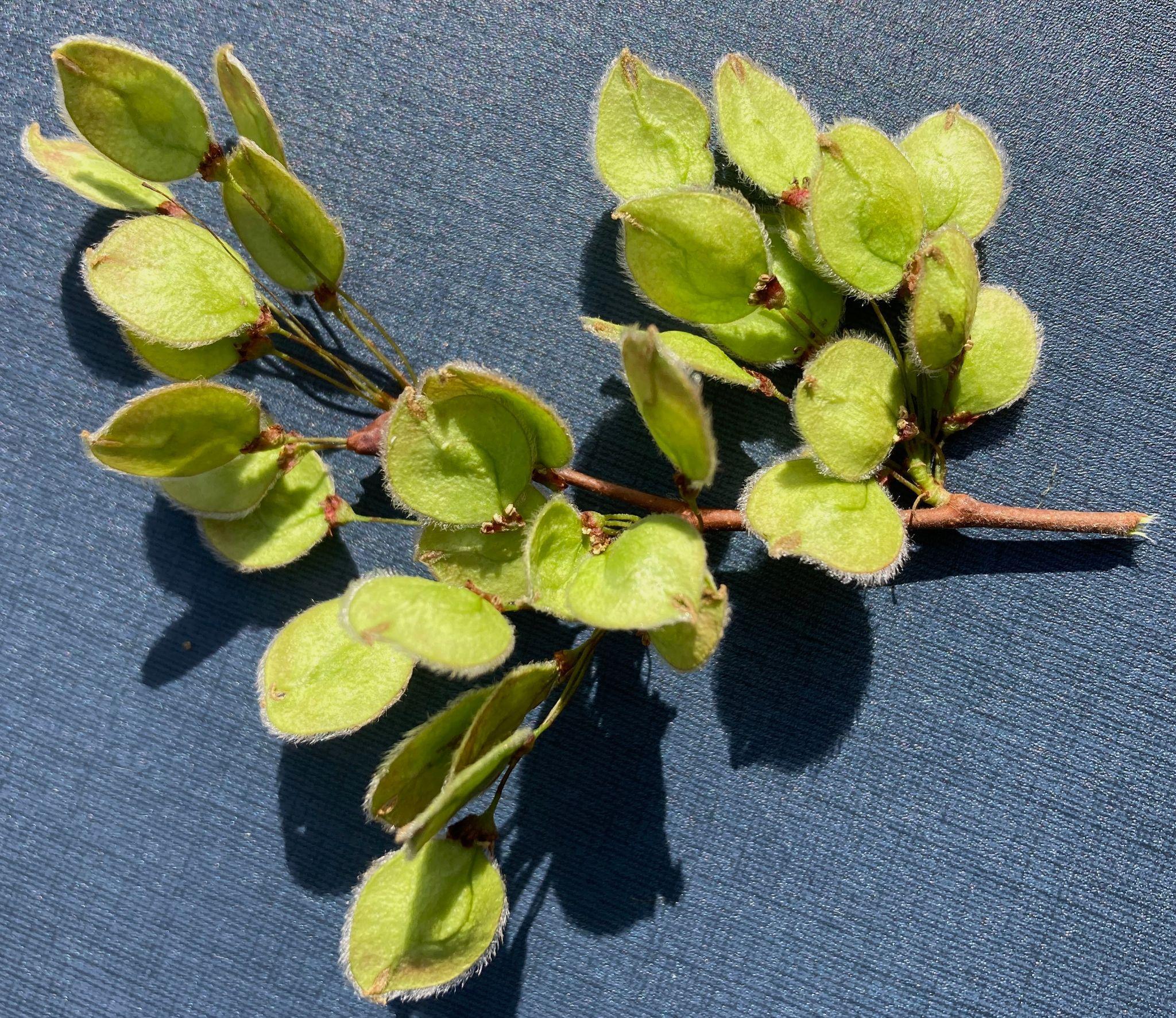

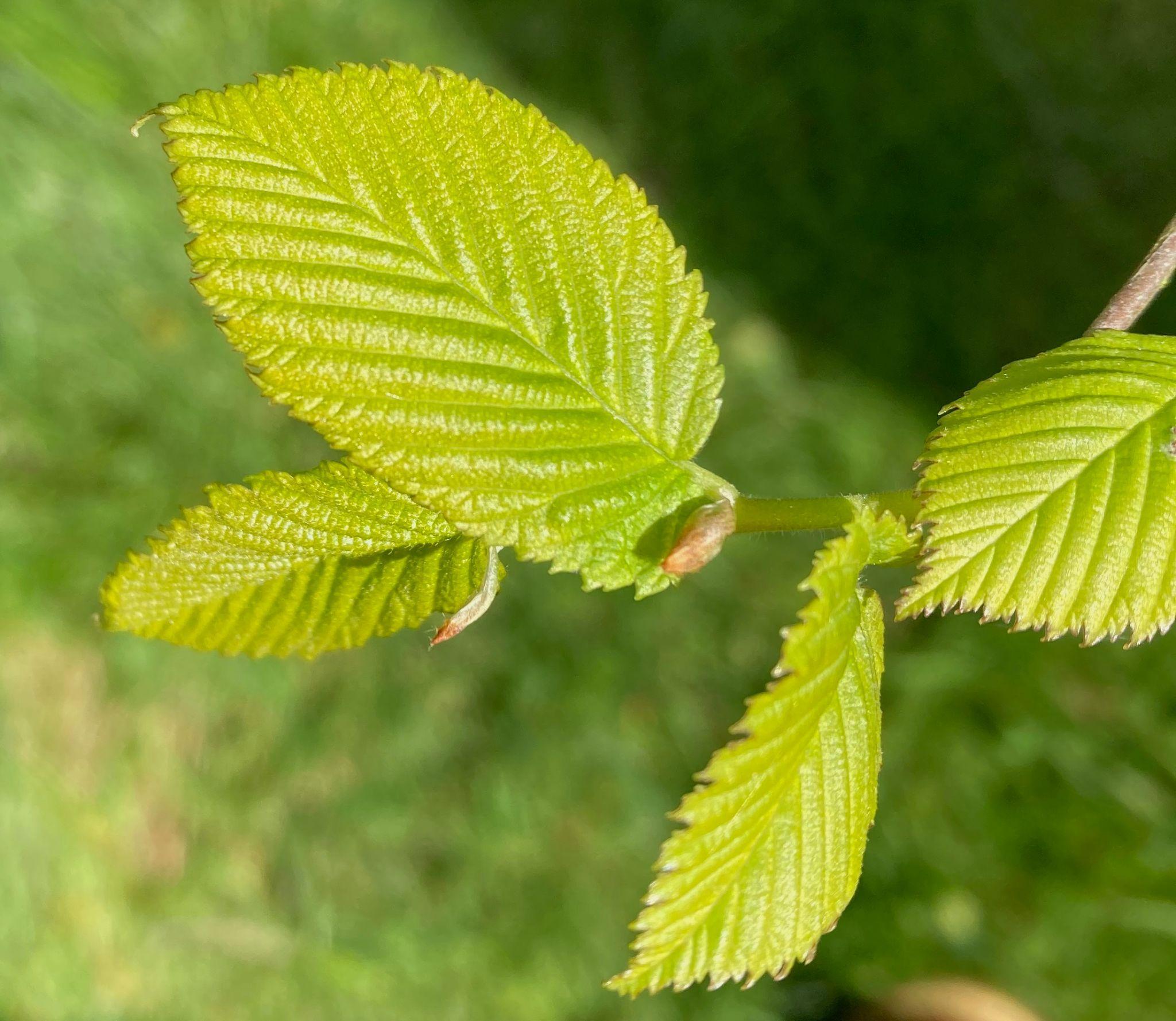

E157 RBG Kew accession 2010-1842*1 labelled as *U. plotii* appears to be *U. laevis*.

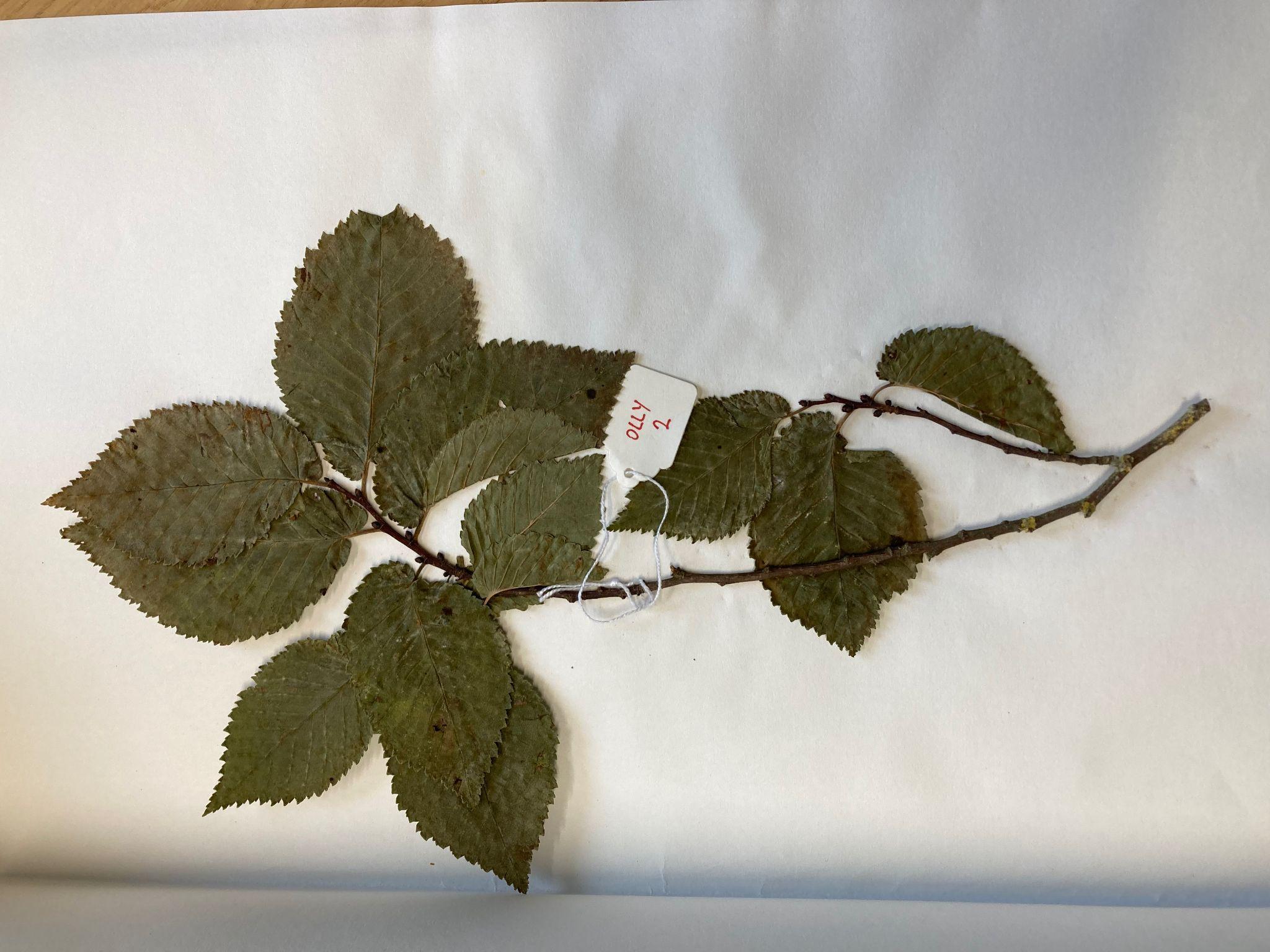

E253 *U. minor* Ogbourne St George, England, 51.458726, -1.71453 (Scale: Jeweller’s tag is 24 mm in width)

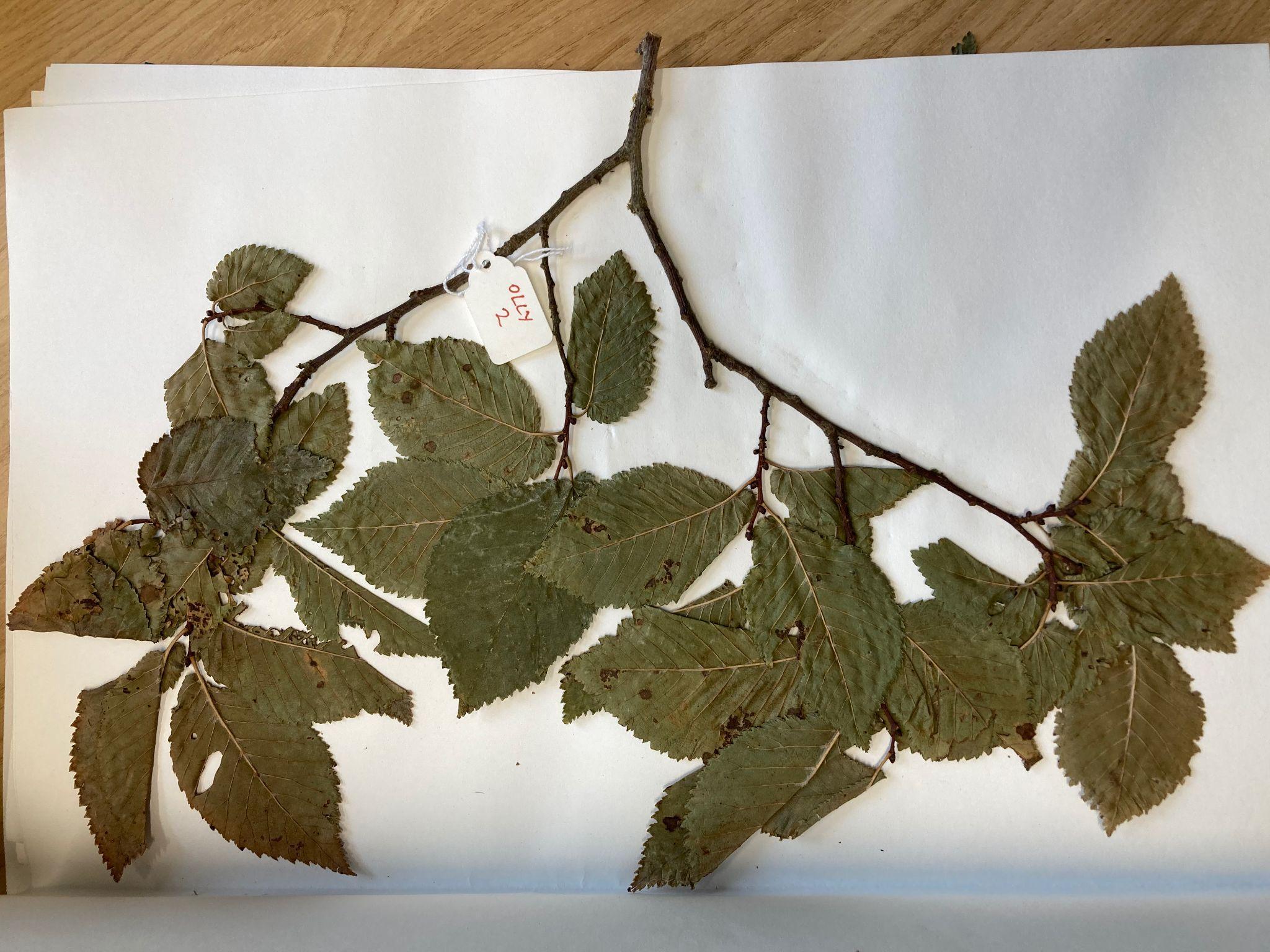

E253 *U. minor* Ogbourne St George, England, 51.458726, -1.71453 (Scale: Jeweller’s tag is 24 mm in width)

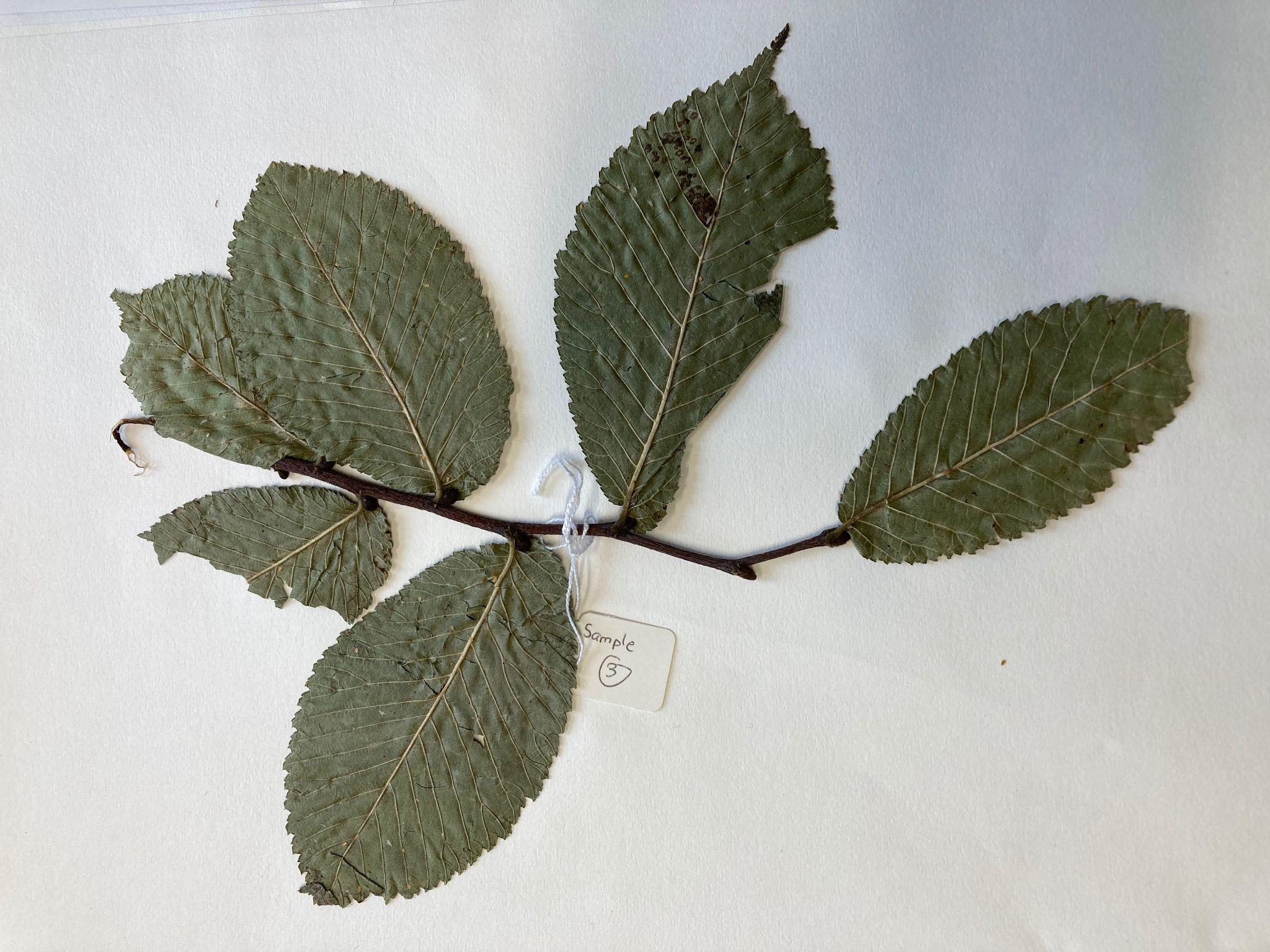

E254 *U. glabra* Perth, Scotland 56.397144, -3.4267545 (Scale: Jeweller’s tag is 24 mm in width)

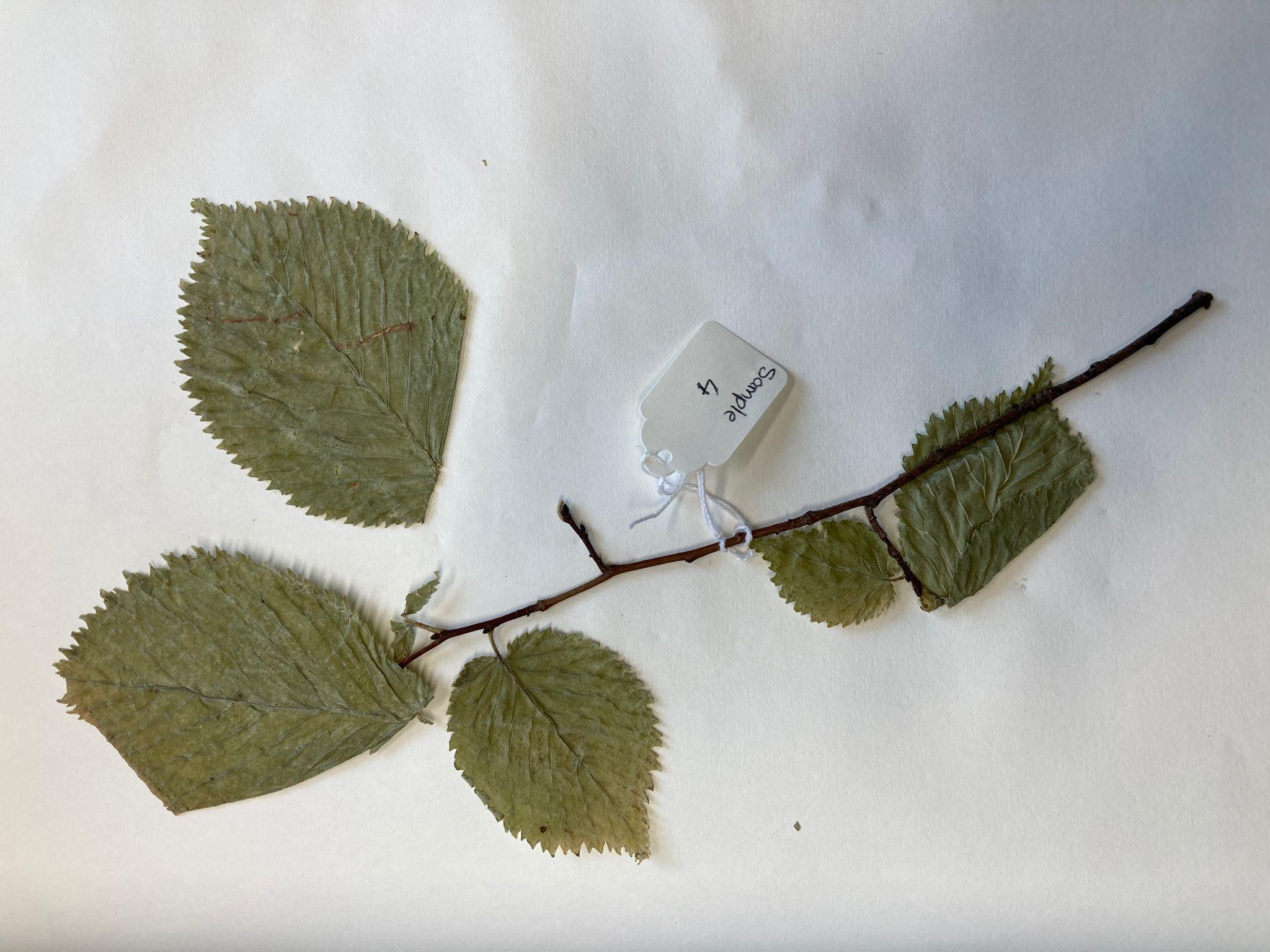

E255 *U. × hollandica s.l.* Sheffield 53.357627, -1.494873 (Scale: Jeweller’s tag is 24 mm in width)

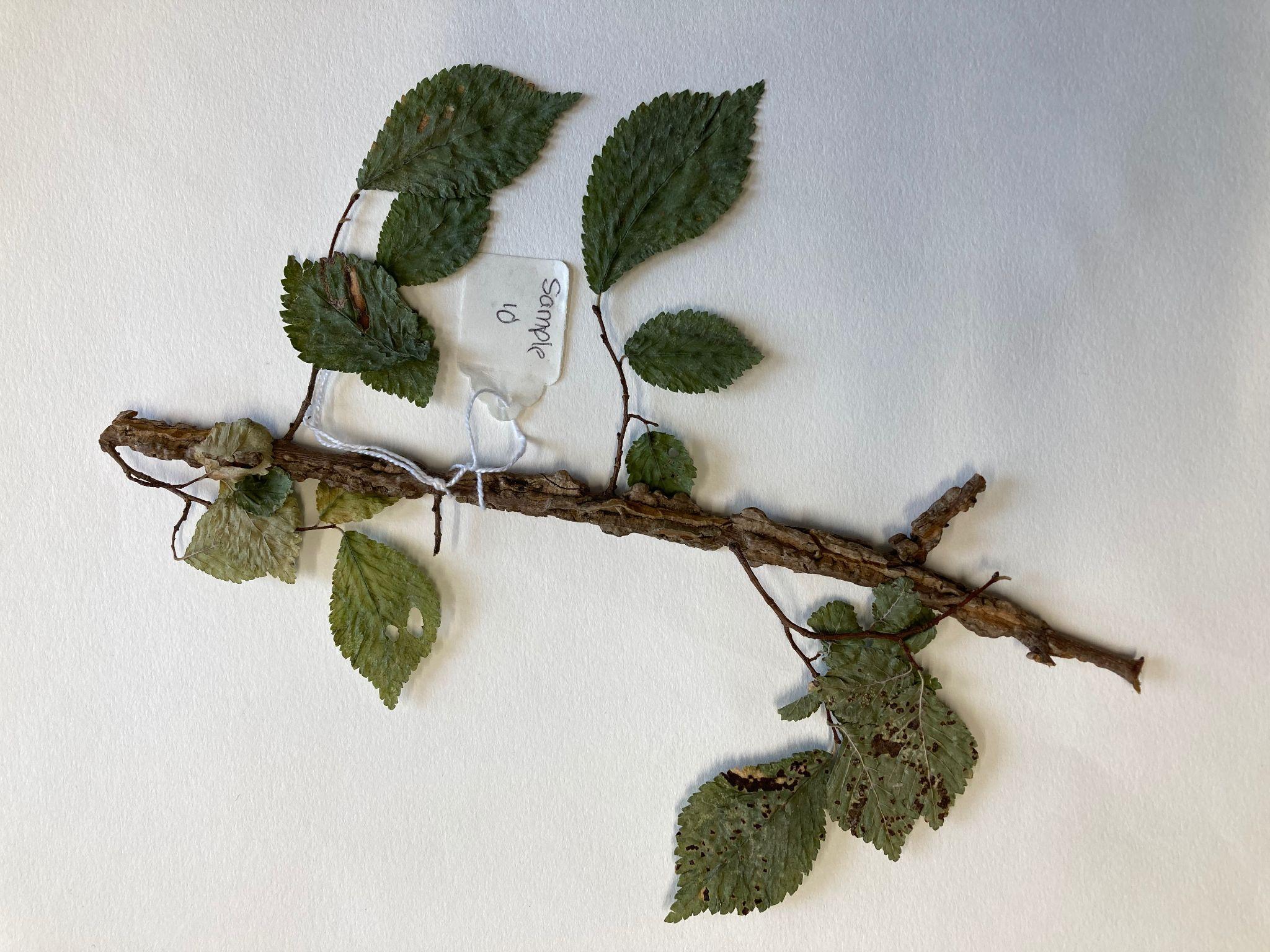

E257 *U. minor* Essex 52.018962, 0.2381907 (Scale: Jeweller’s tag is 24 mm in width)

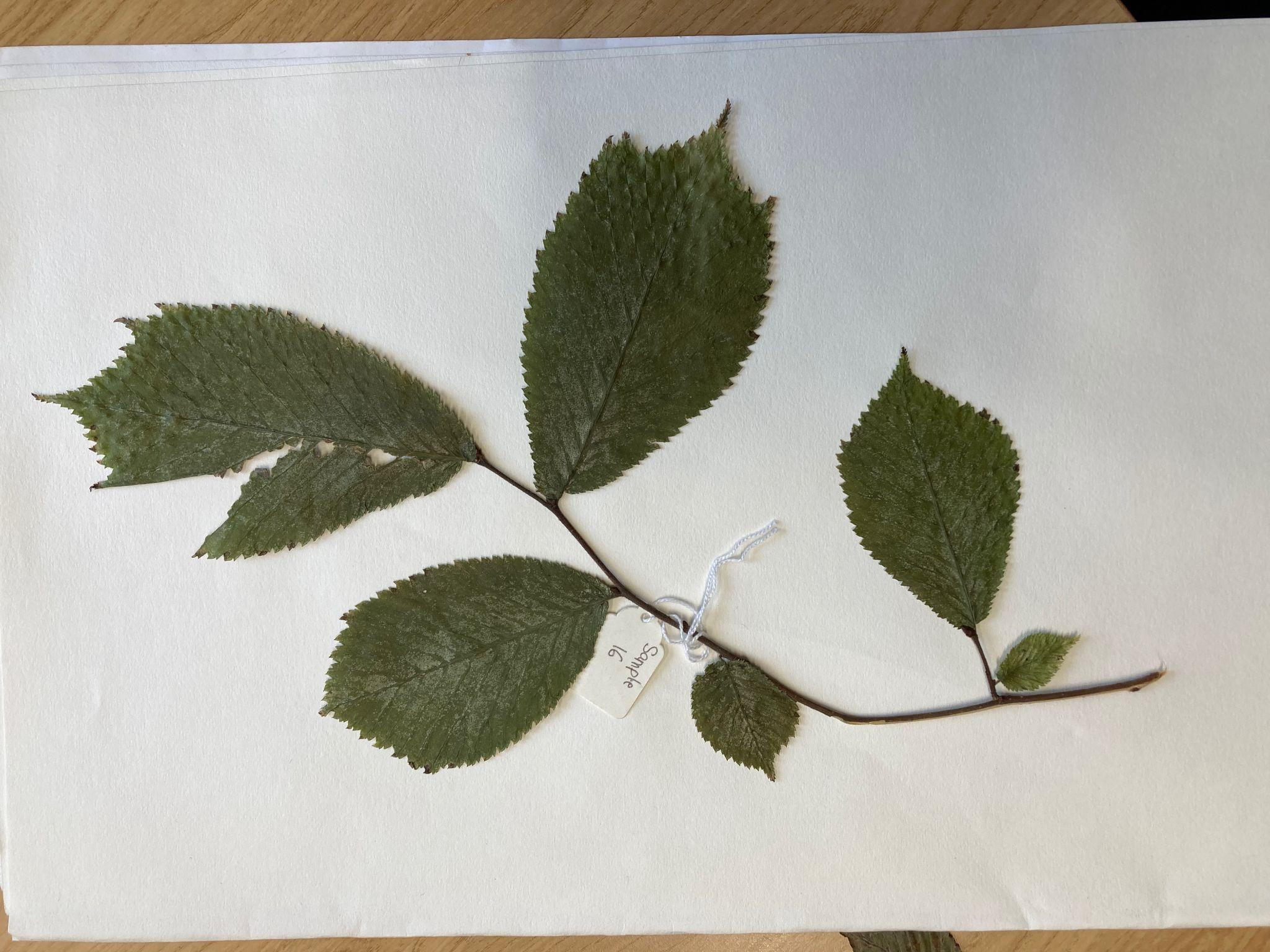

E258 *U. glabra* London 51.530204, -0.10889411 (Scale: Jeweller’s tag is 24 mm in width)

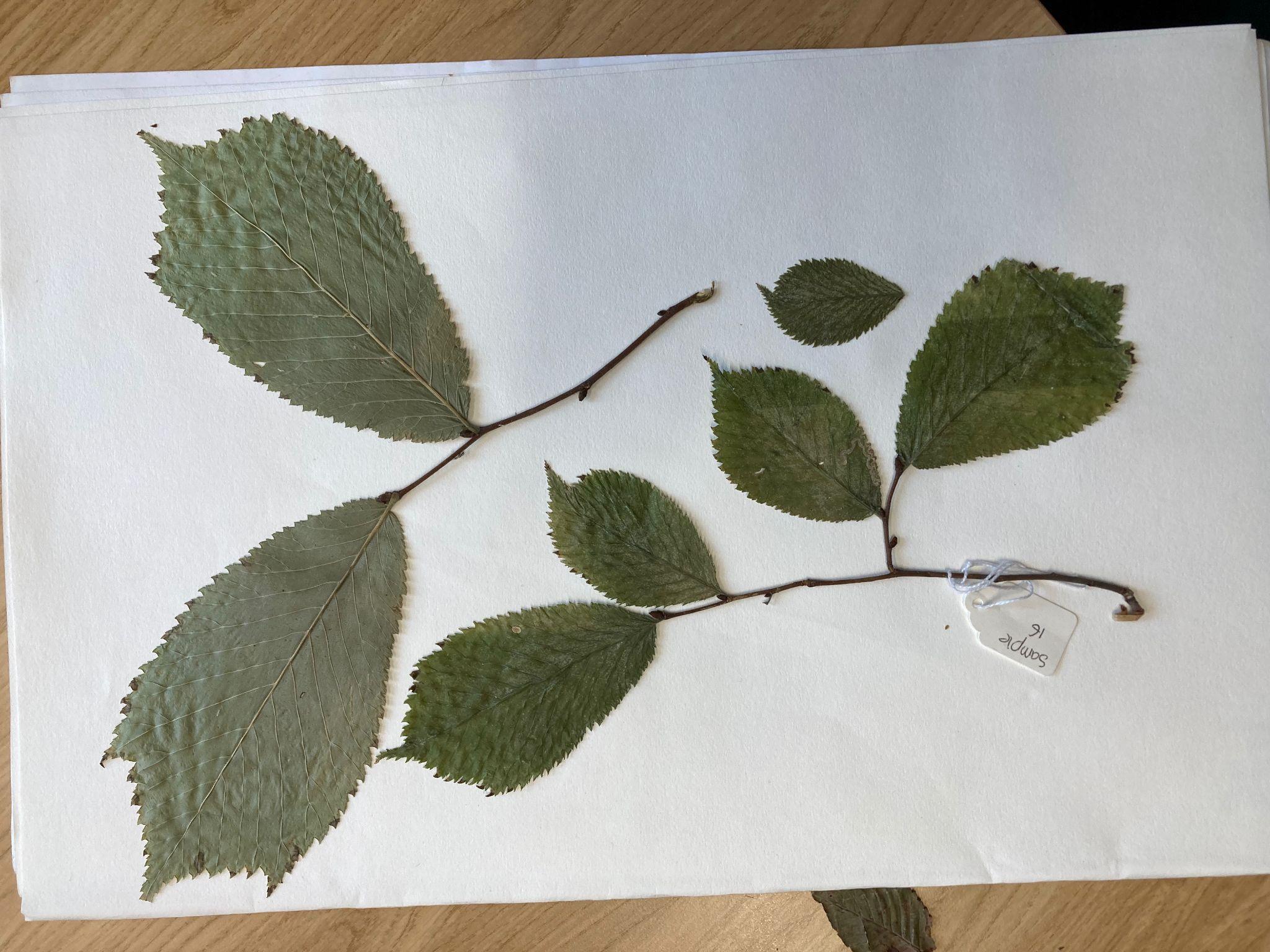

E258 *U. glabra* London 51.530204, -0.10889411 (Scale: Jeweller’s tag is 24 mm in width)

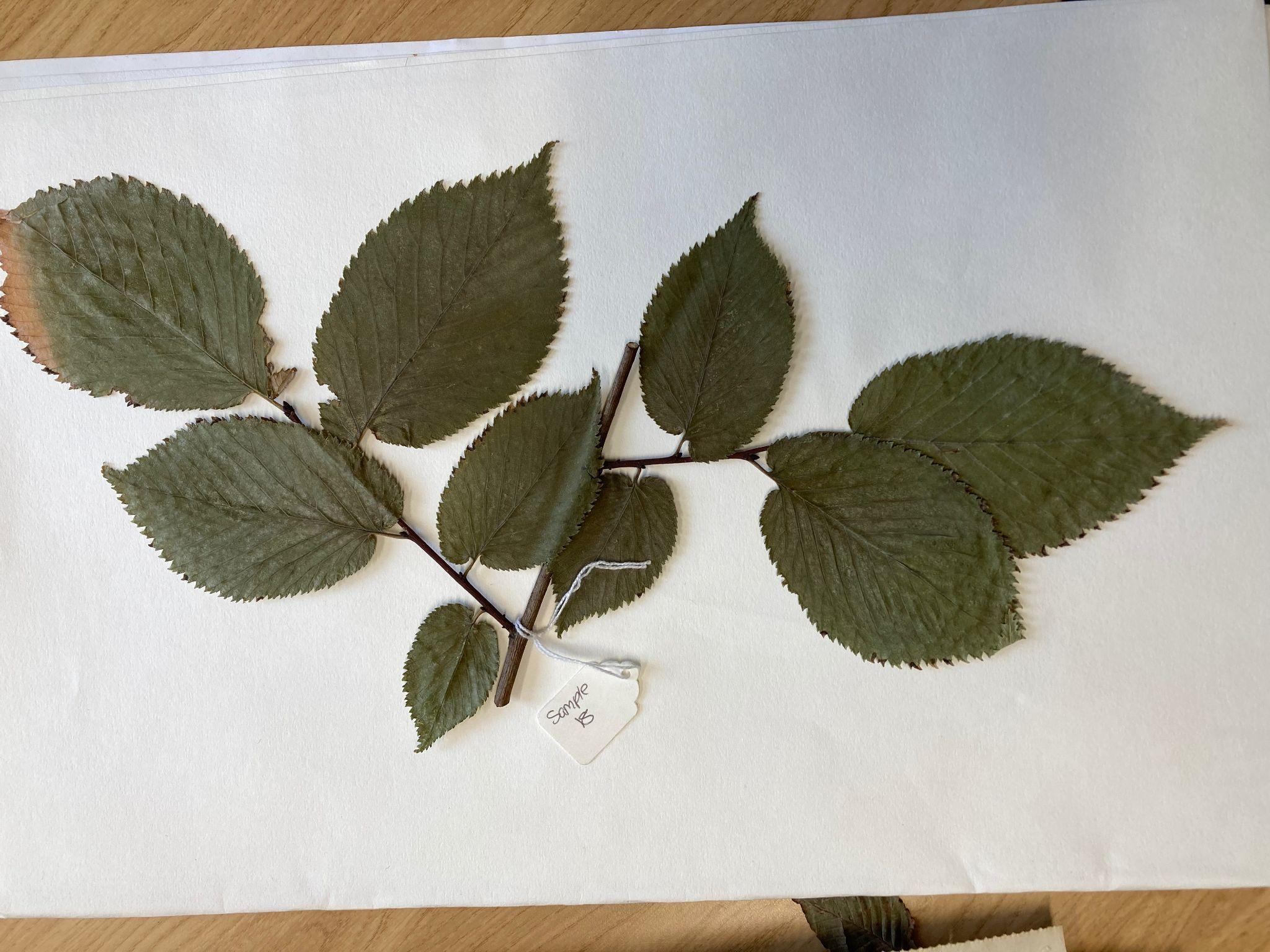

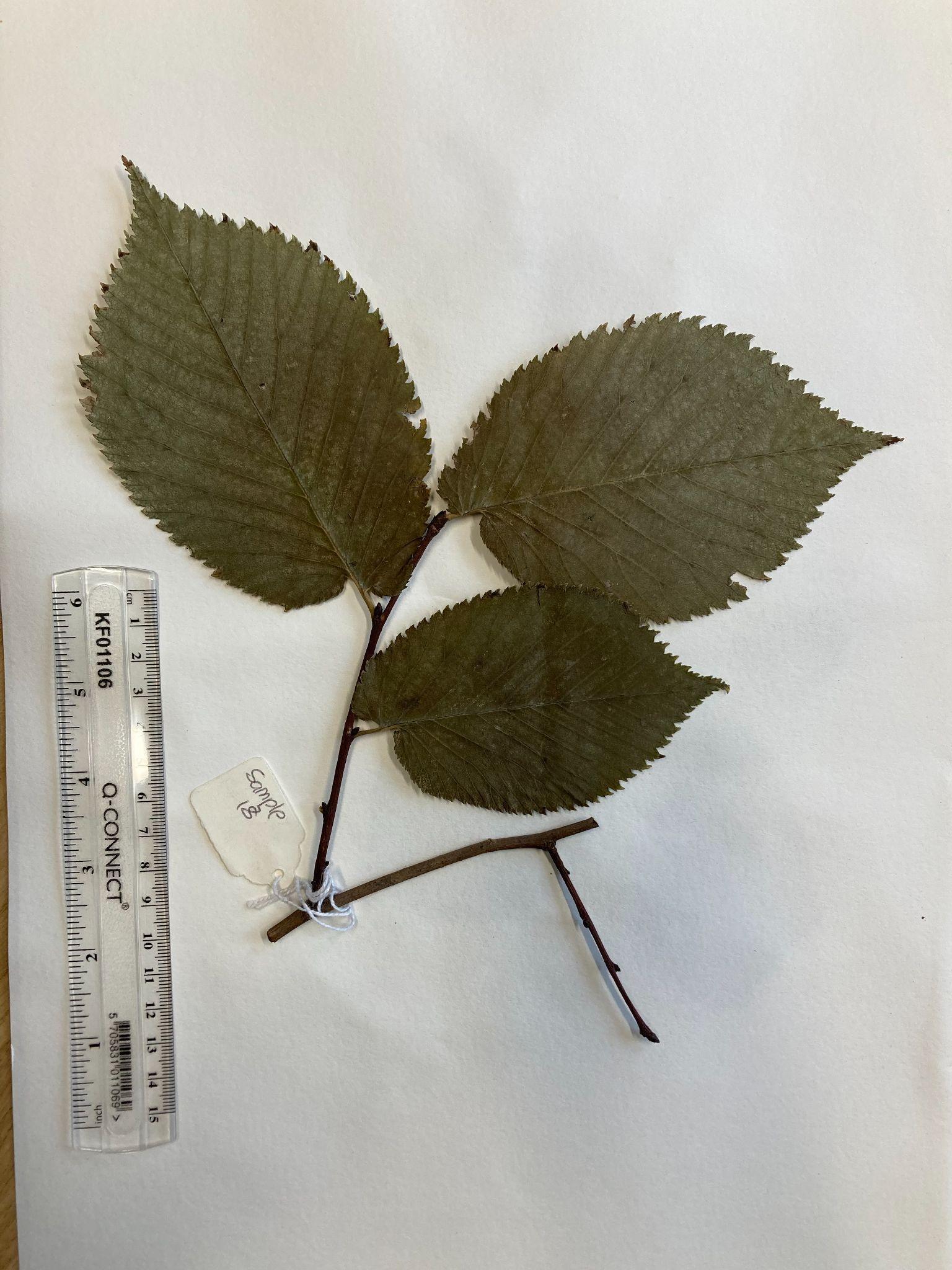

E259 *U. × hollandica s.l.* London 51.543226, -0.10208365 (Scale: Jeweller’s tag is 24 mm in width)

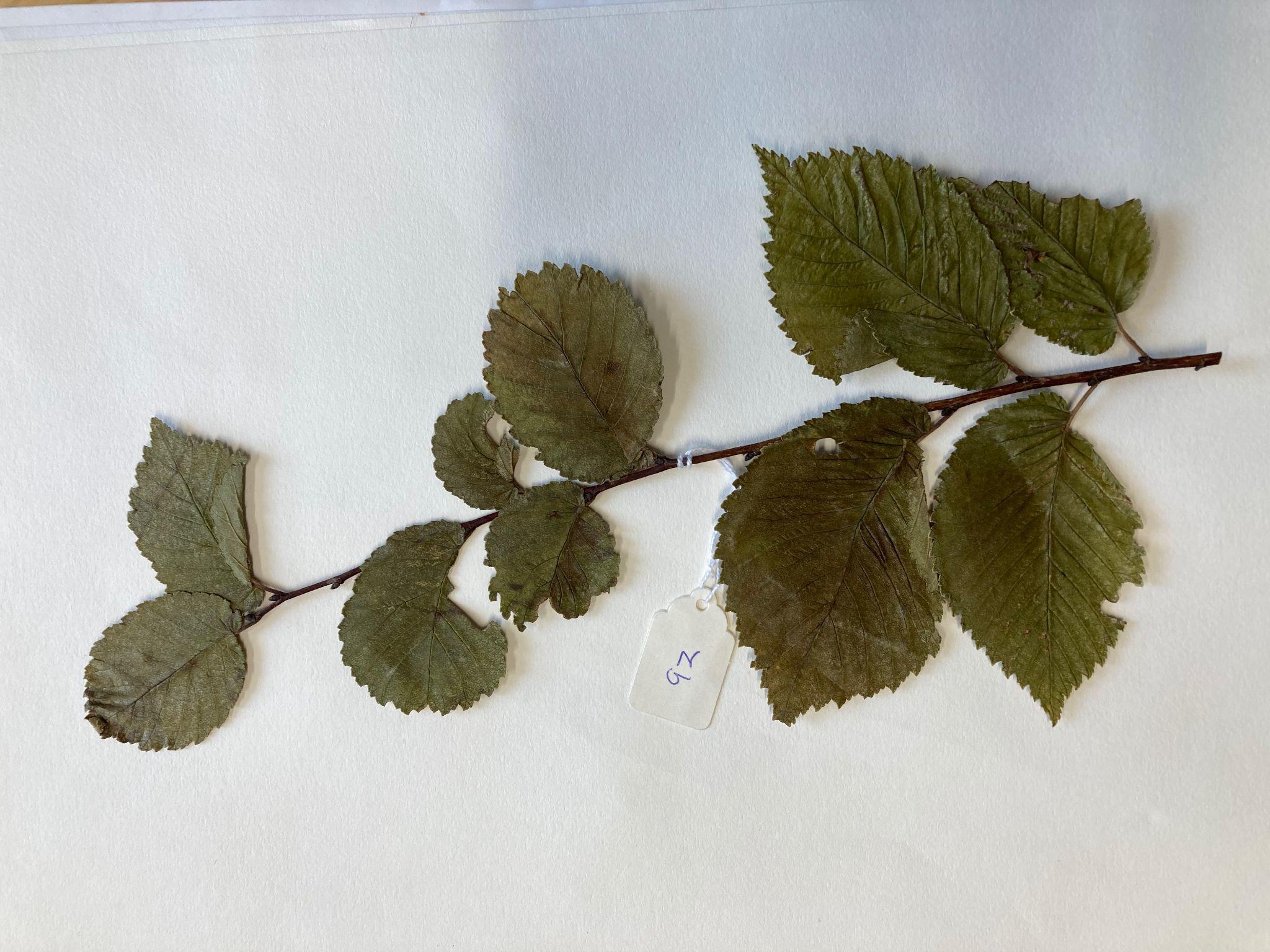

E261 *U. minor* Brighton 50.866669, -0.002134 (Scale: Jeweller’s tag is 24 mm in width)

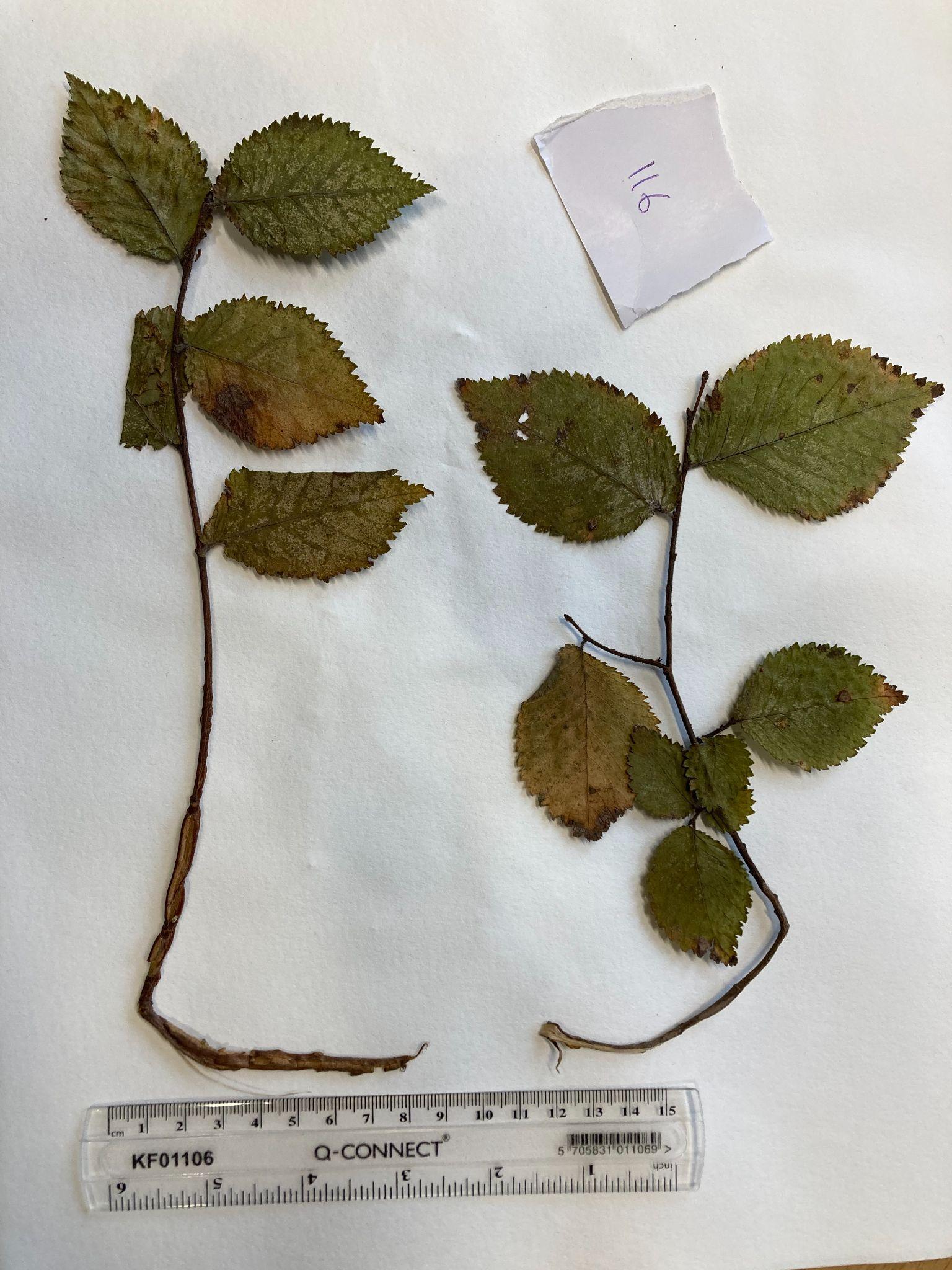

E262 *U. minor* Brighton 50.861409, 0.064348

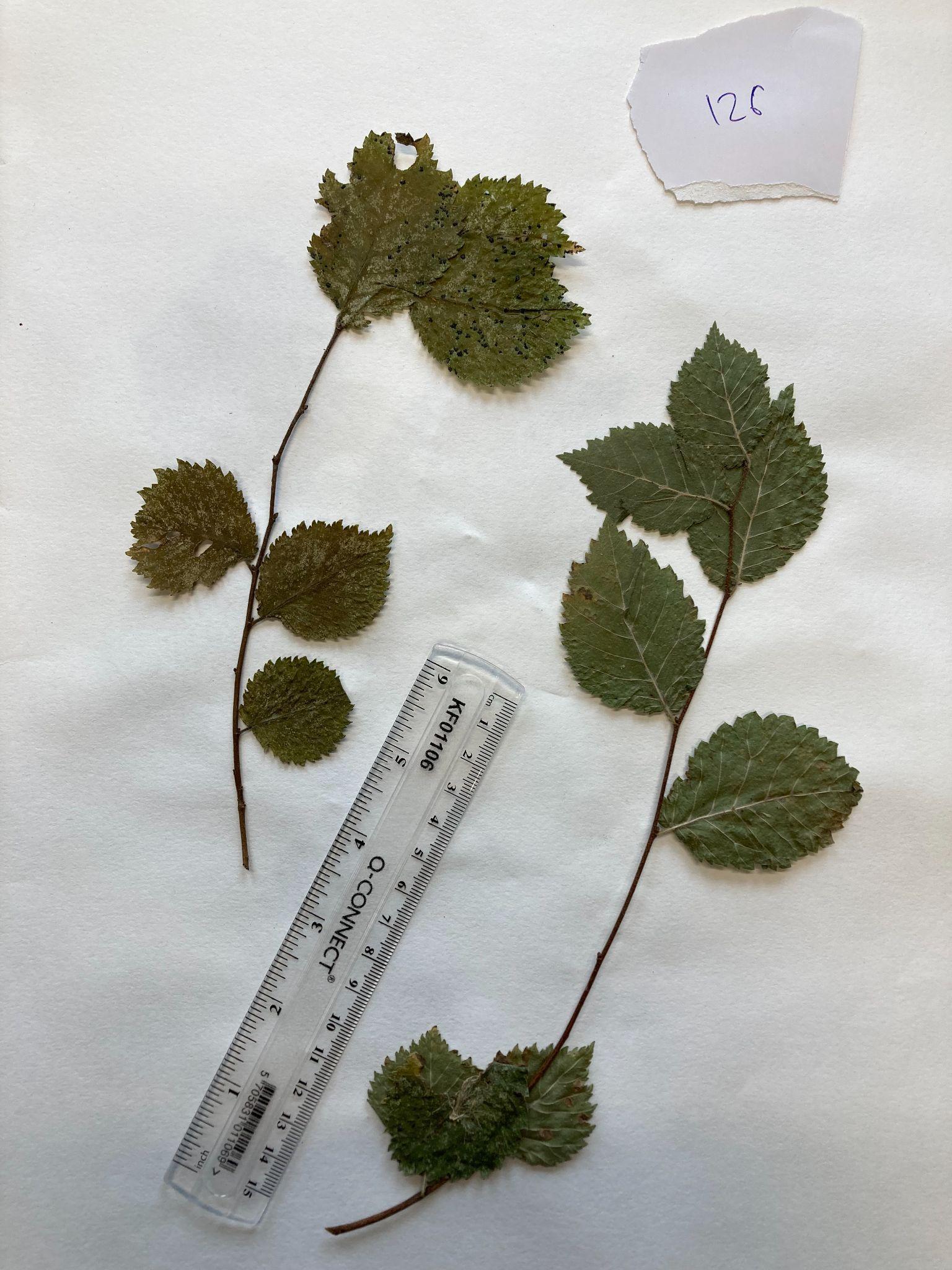
E263 *U. minor* Brighton 50.823809, 0.182489

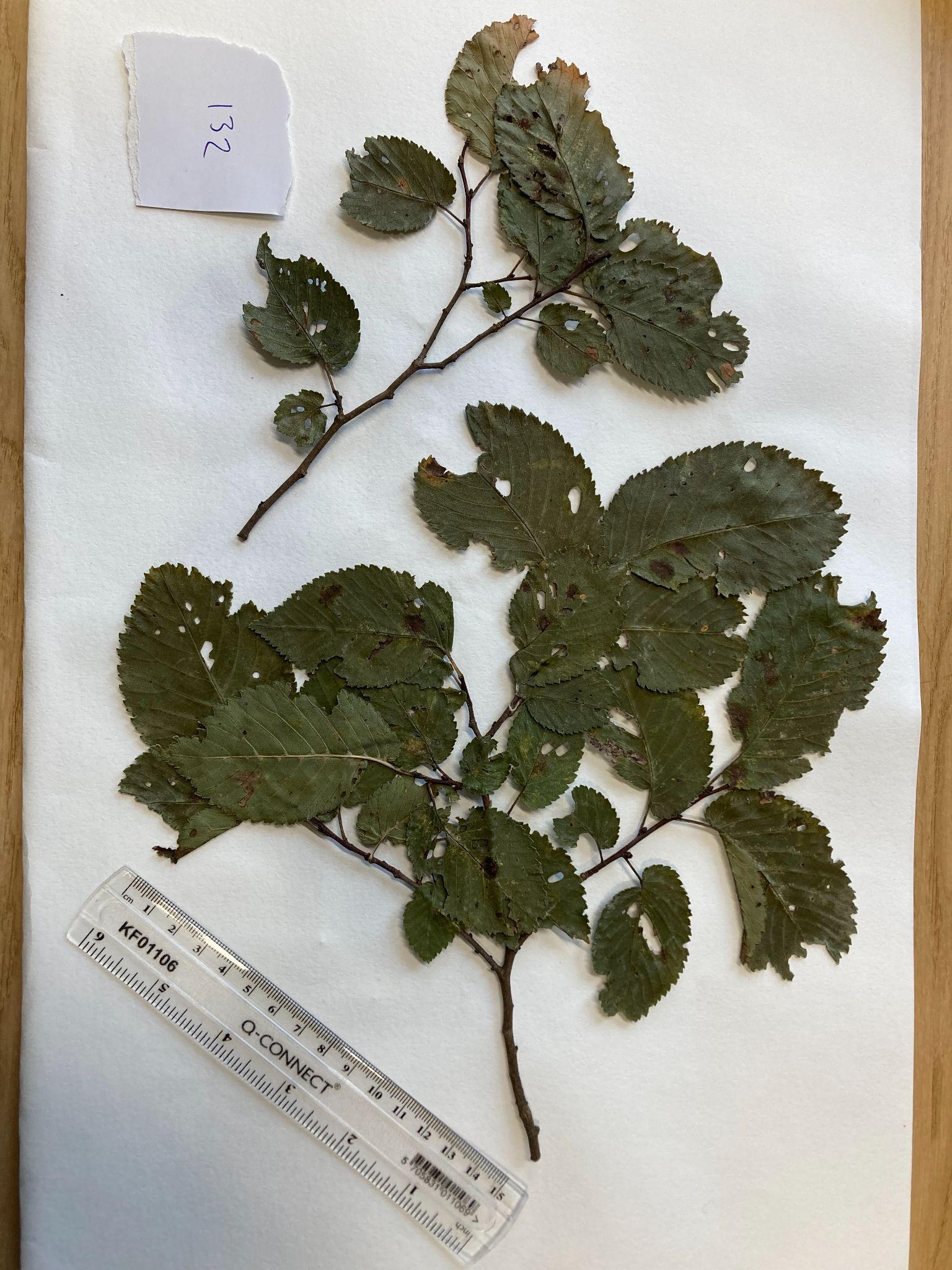
E264 *U. minor* Suffolk 52.104956, 1.1039543

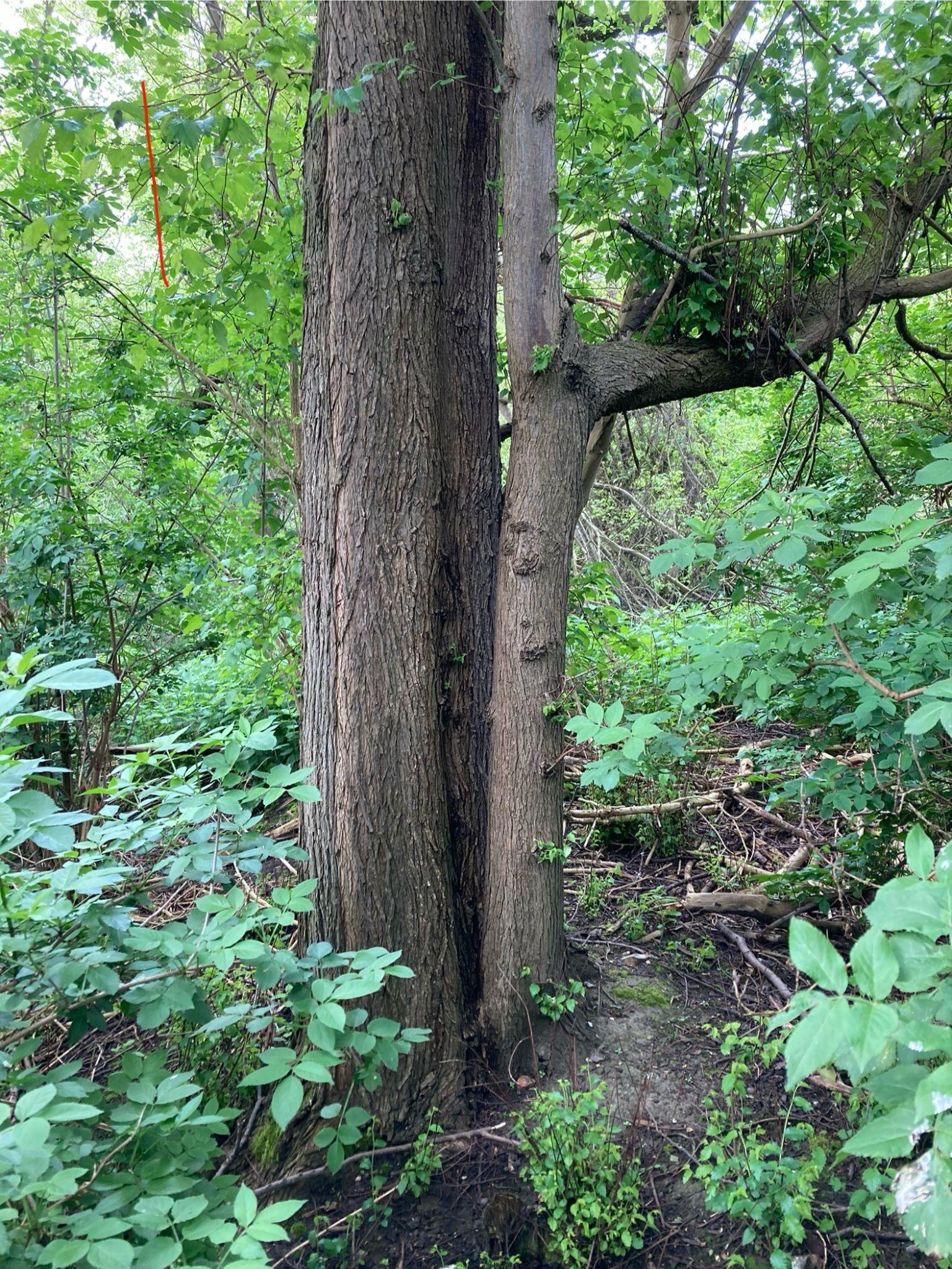

E288 *U. minor* Tonge Mill, tree 1 51°20'20.6"N, 0°46'26.2"E (Summer 2024)

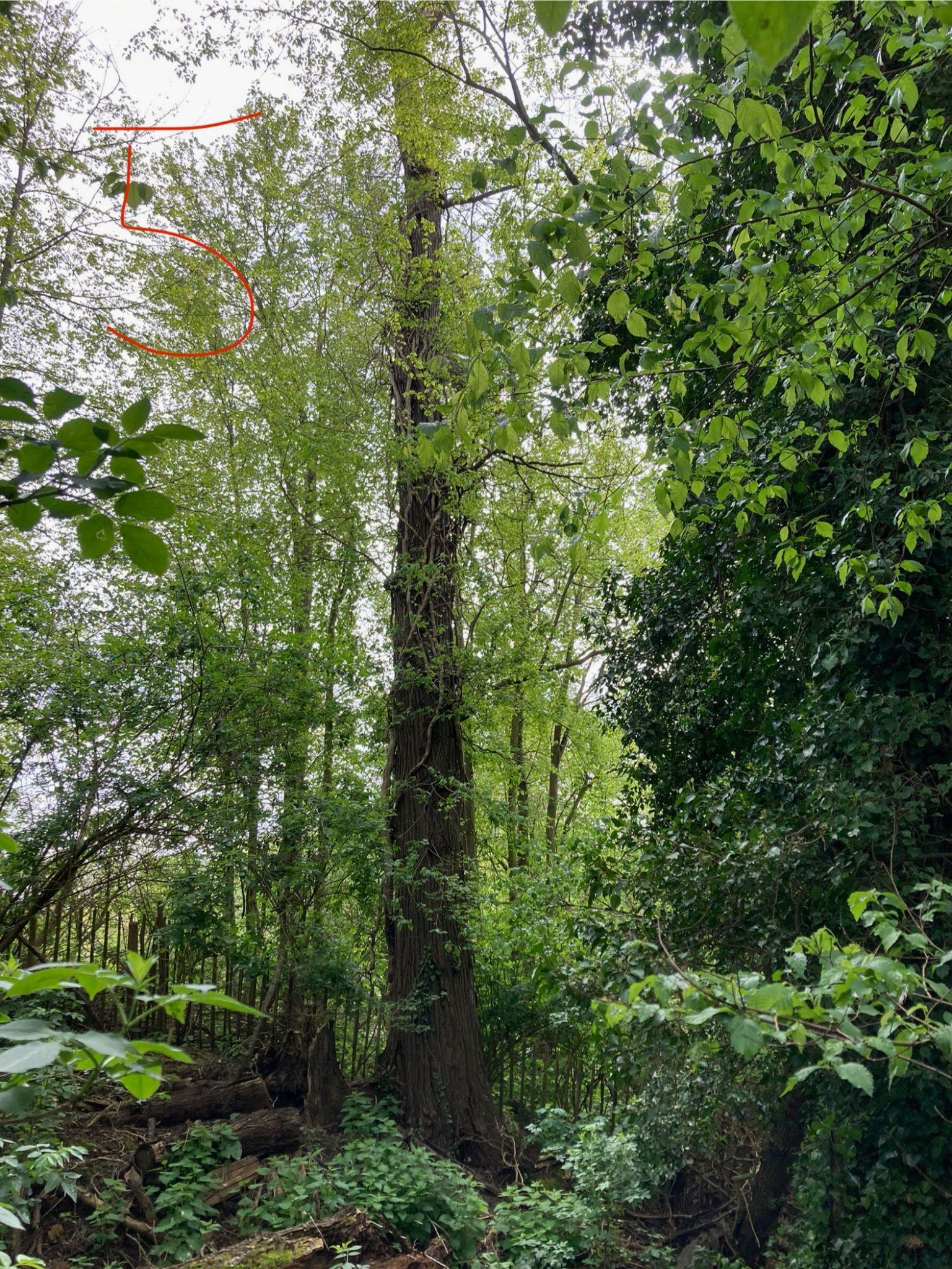

E289 *U. minor* Tonge Mill, tree 5, 51°20'19.1"N, 0°46'25.3"E (Summer 2024)

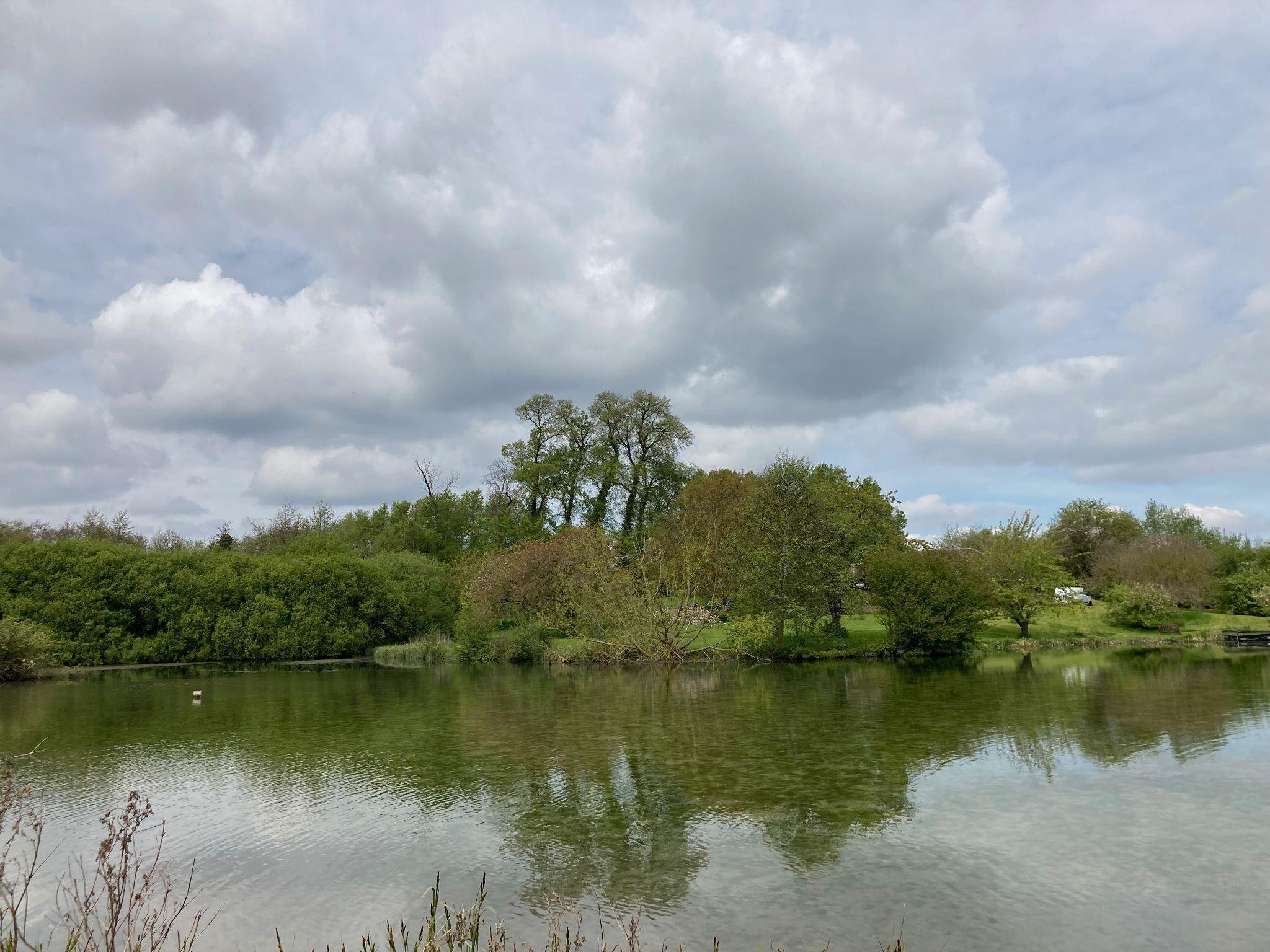

*U. minor* stand at Tonge Mill containing E288 and E289 (tall central trees) 51°20'19.1"N, 0°46'25.3"E (Summer 2024)

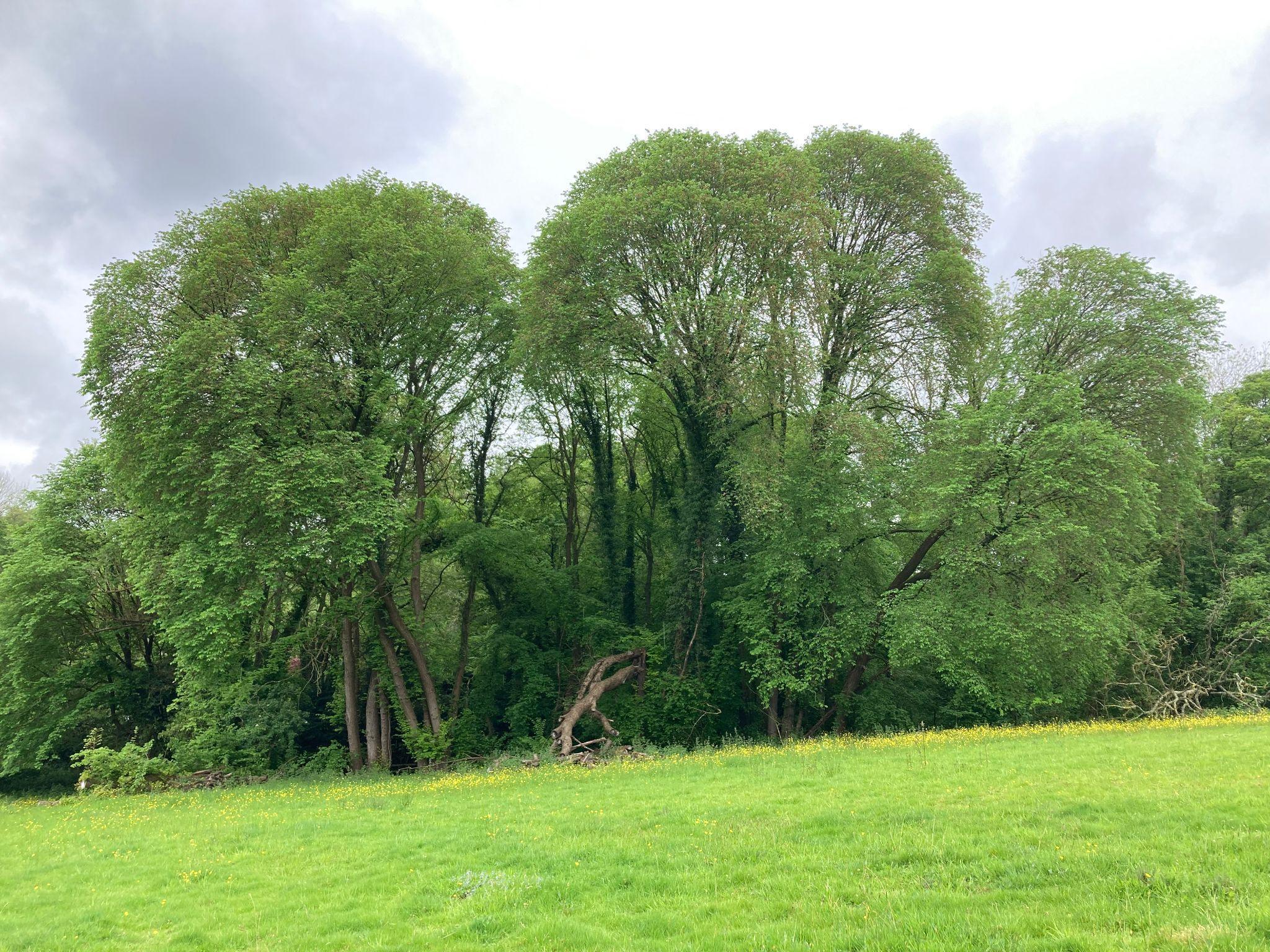

E290 *U. × hollandica s.l.* Amhurst Mill, Kent 51°09'50.7"N, 0°19'58.9"E (Summer 2024)

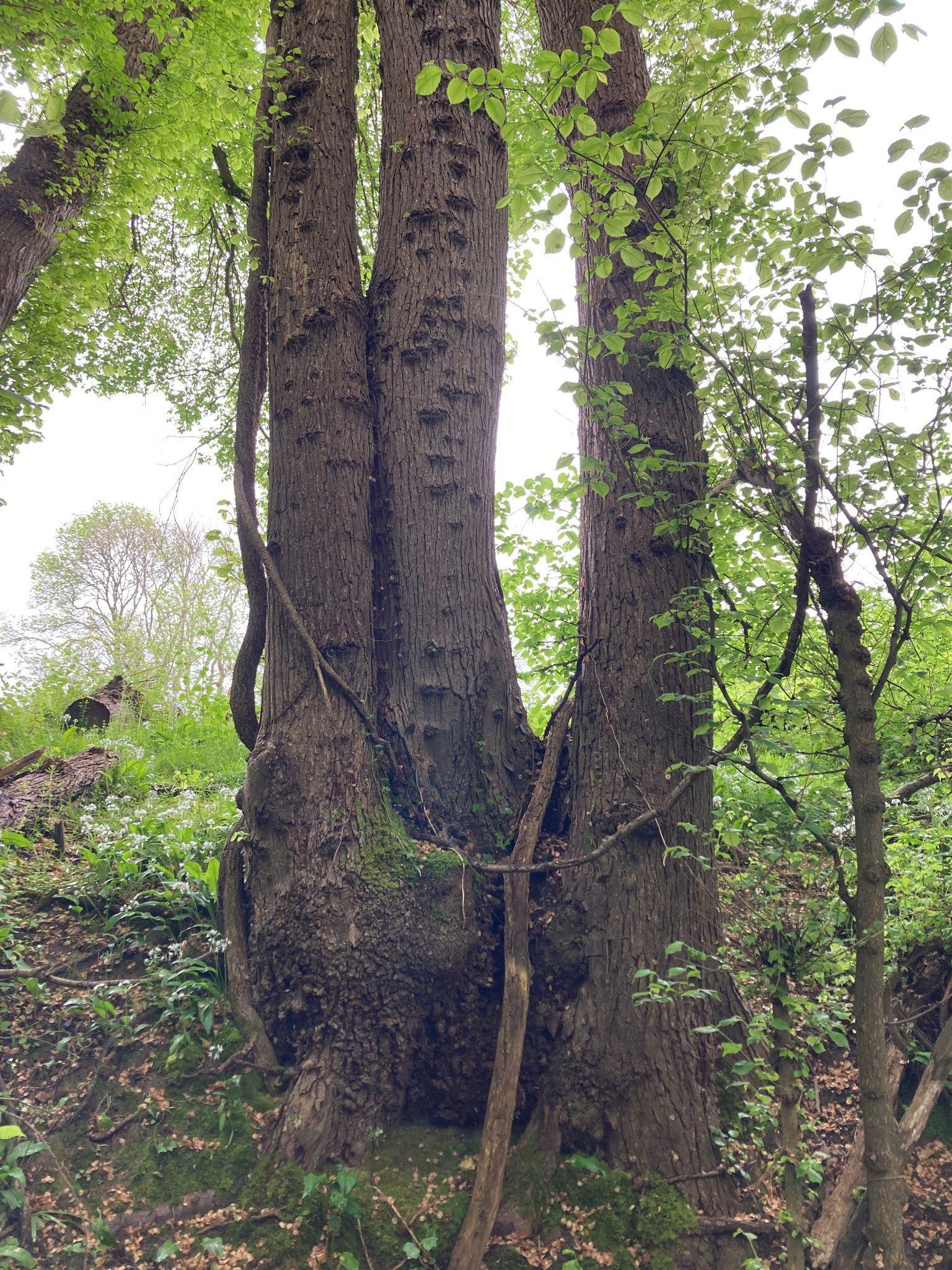

E290 *U. × hollandica s.l.* Amhurst Mill, Kent 51°09'50.7"N, 0°19'58.9"E (Summer 2024)

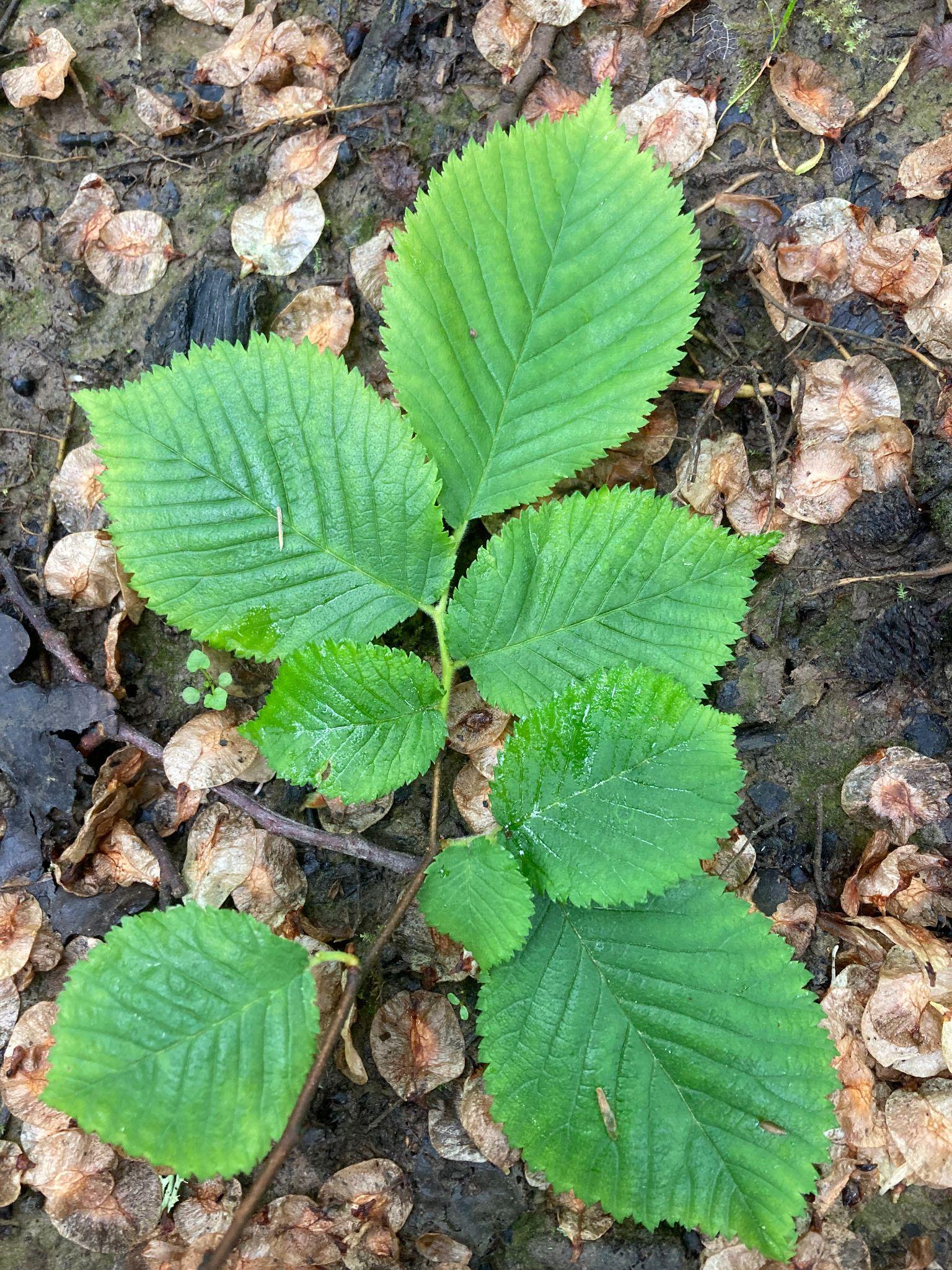

E290 *U. × hollandica s.l.* Amhurst Mill, Kent 51°09'50.7"N, 0°19'58.9"E (Summer 2024)

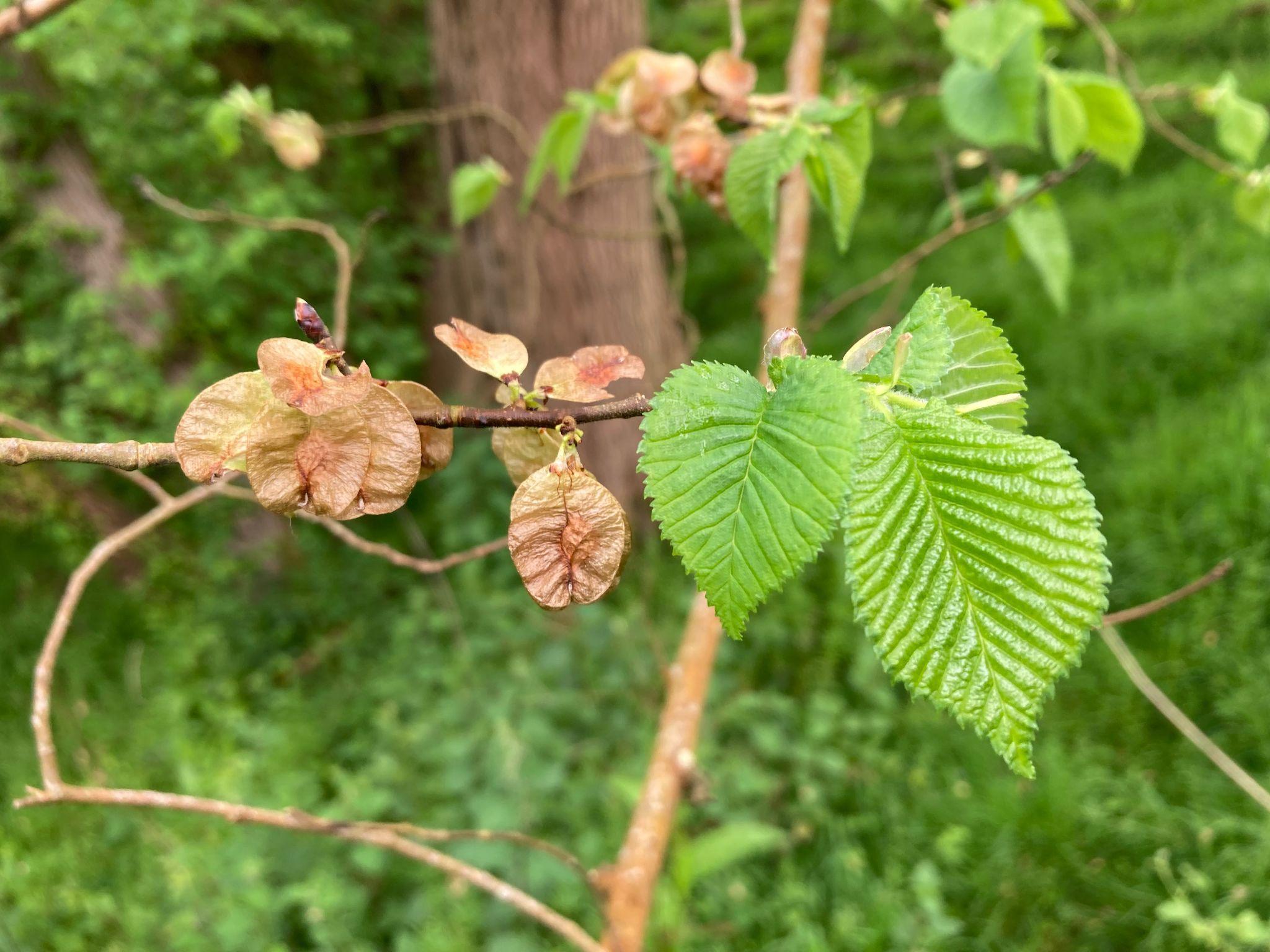

E291 *U. × hollandica s.l.* Amhurst “Amphitheatre”, Kent 51°09'51.9"N, 0°20'02.2"E (Summer 2024)

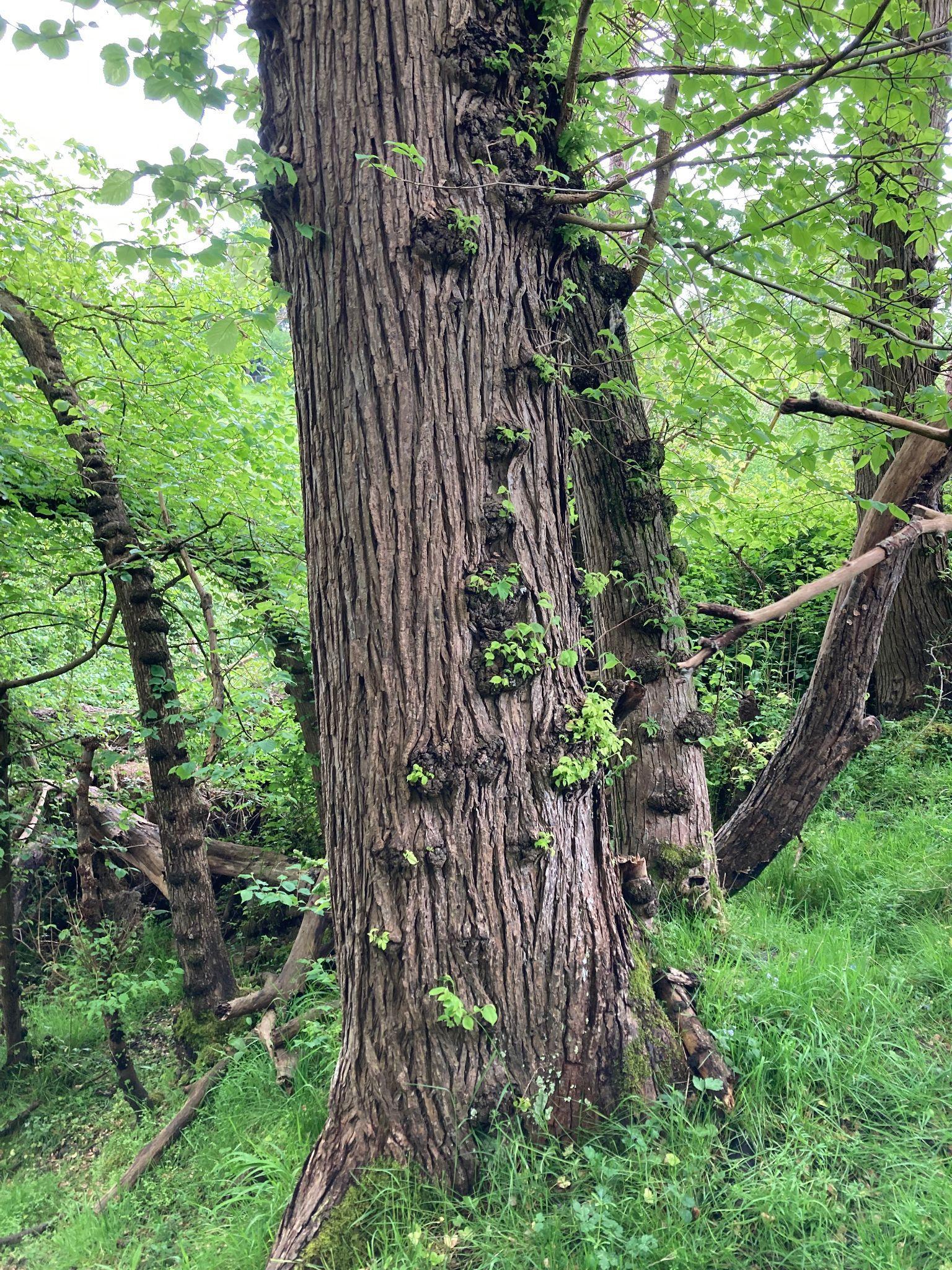

E291 *U. × hollandica s.l.* Amhurst “Amphitheatre”, Kent. 51°09'51.9"N, 0°20'02.2"E (Summer 2024)

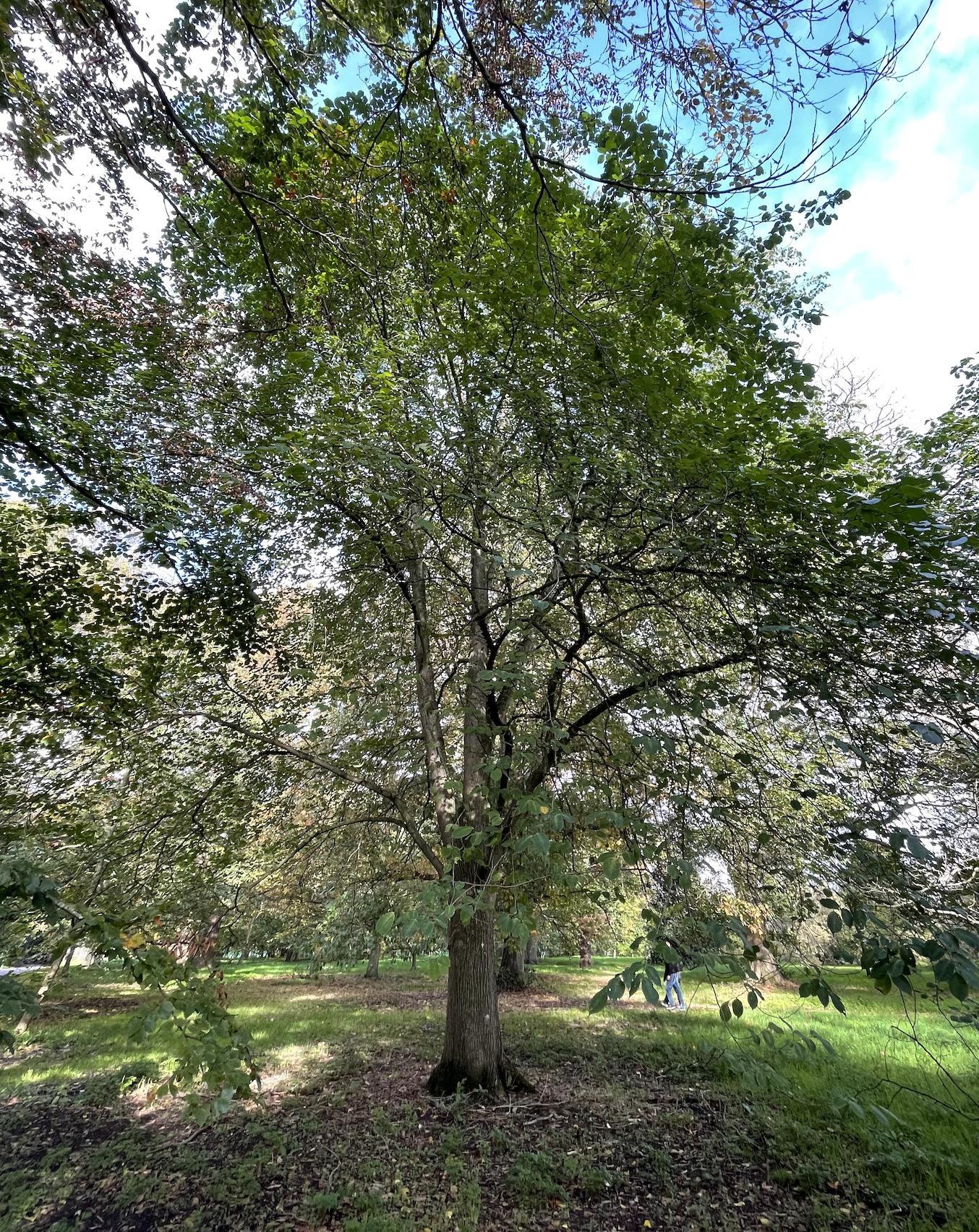

E148, *Ulmus glabra*, Kew Gardens 1959-23702*1

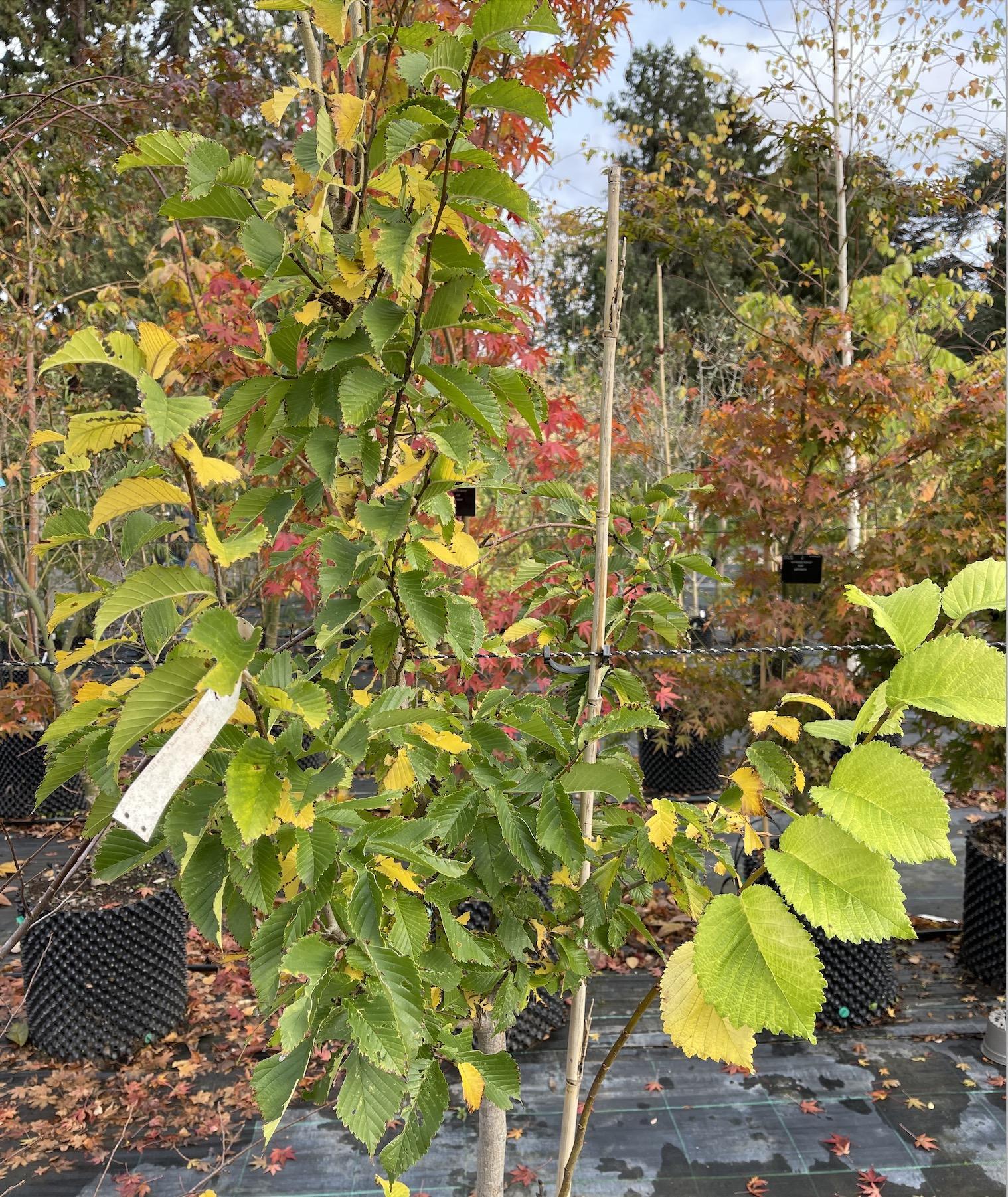

E158, *Ulmus*, San Zanobi (Plantyn' × *U. pumila*.) Kew Gardens (2019-2712*2)

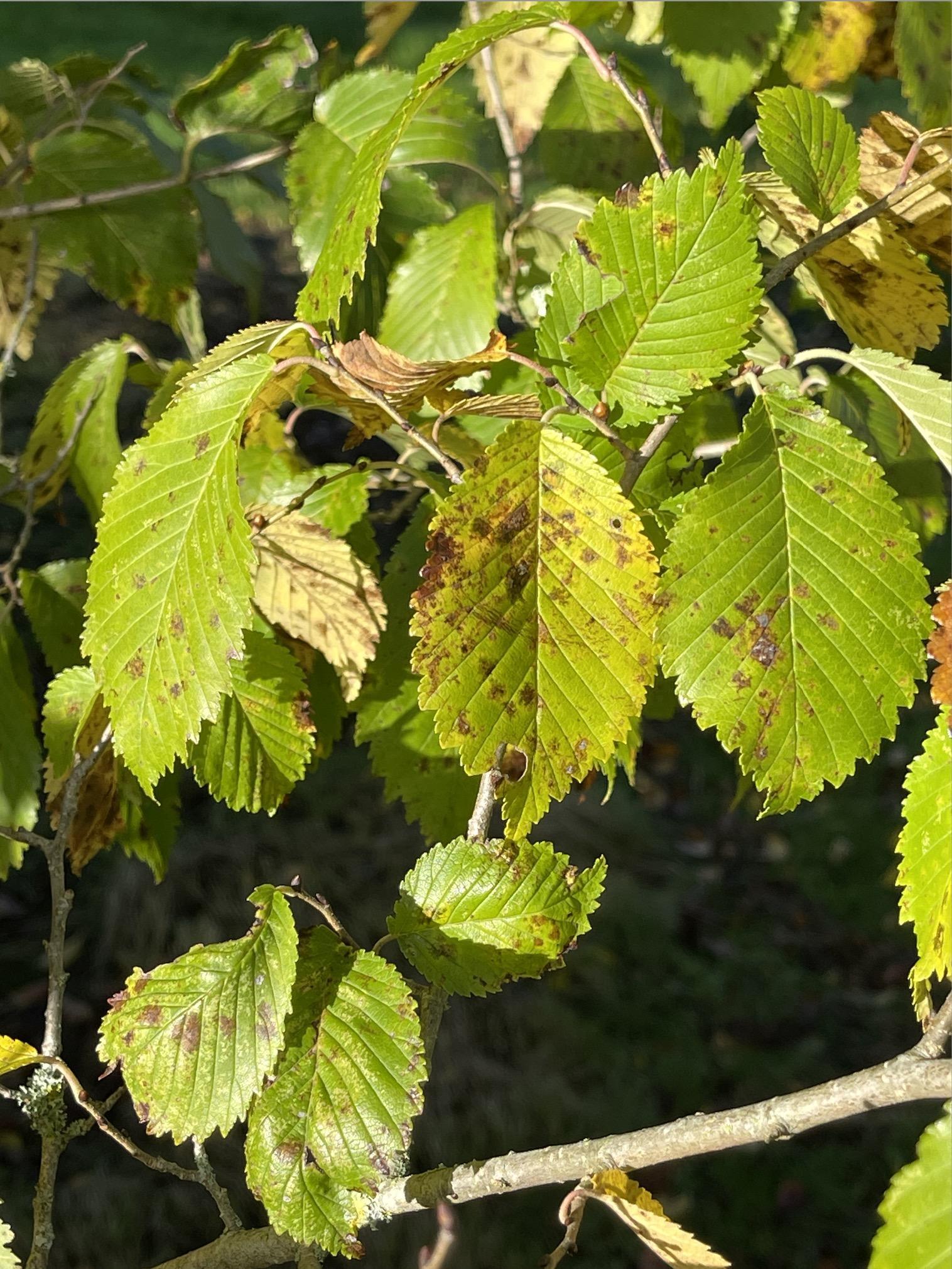

E091, *Ulmus*, FL610 (1038 [1038 is 799 *×* 410] *×* *U. japonica*), Sir Harold Hillier Gardens, 2007.0259*A

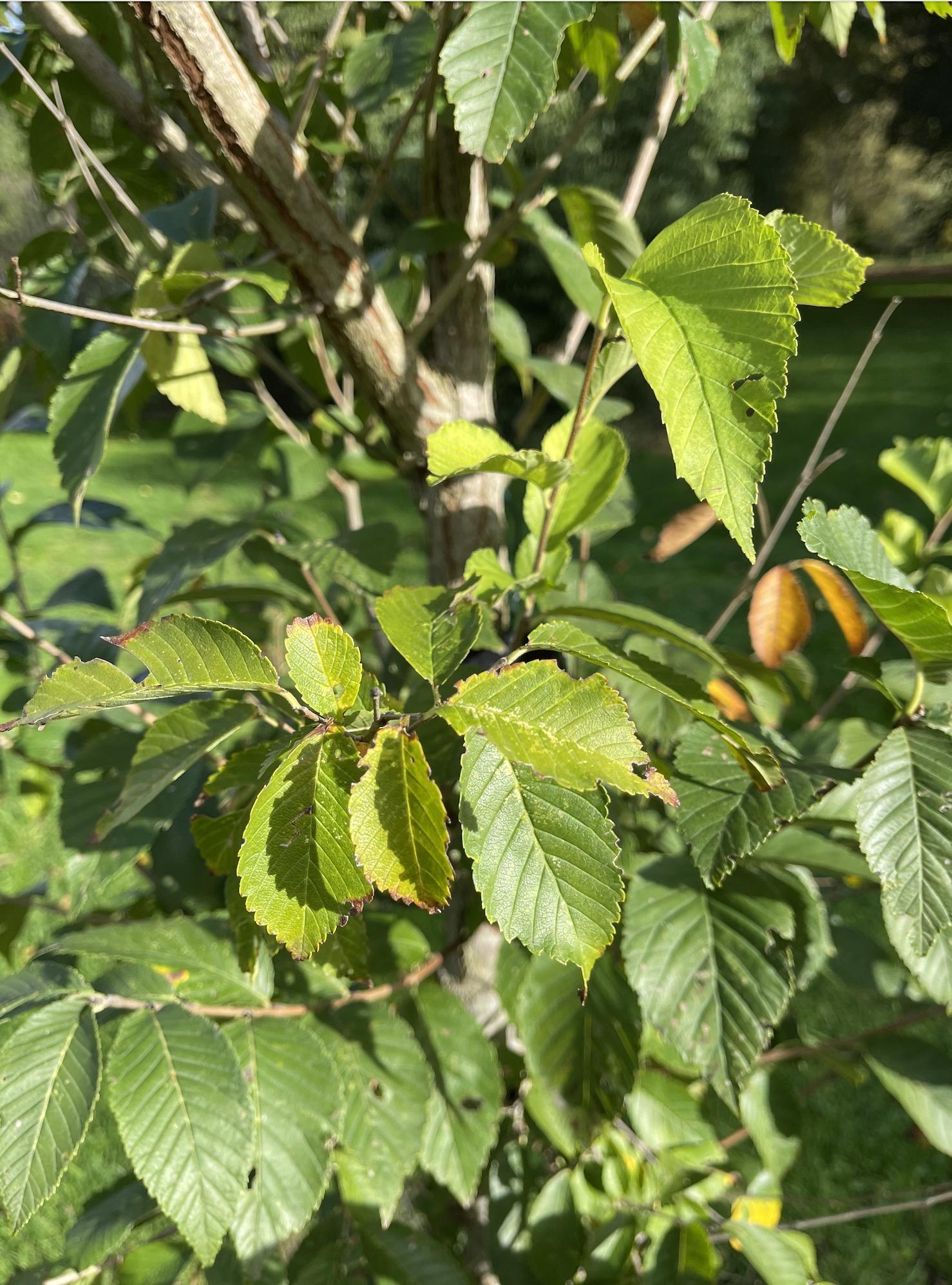

E154, *Ulmus* New Horizon (*U. davidiana* var. *japonica* × *U. pumila*), Kew Gardens (2020-203*1)

#### Fig. S2. Principal component analysis of whole-genome sequence data from all 204 sequenced elm accessions. Panels show principal components (a) 3 and 4, (b) 5 and 6, (c) 7 and 8, and (d) 9 and 10. Colours and group definitions follow Fig. 1.

**(a)**

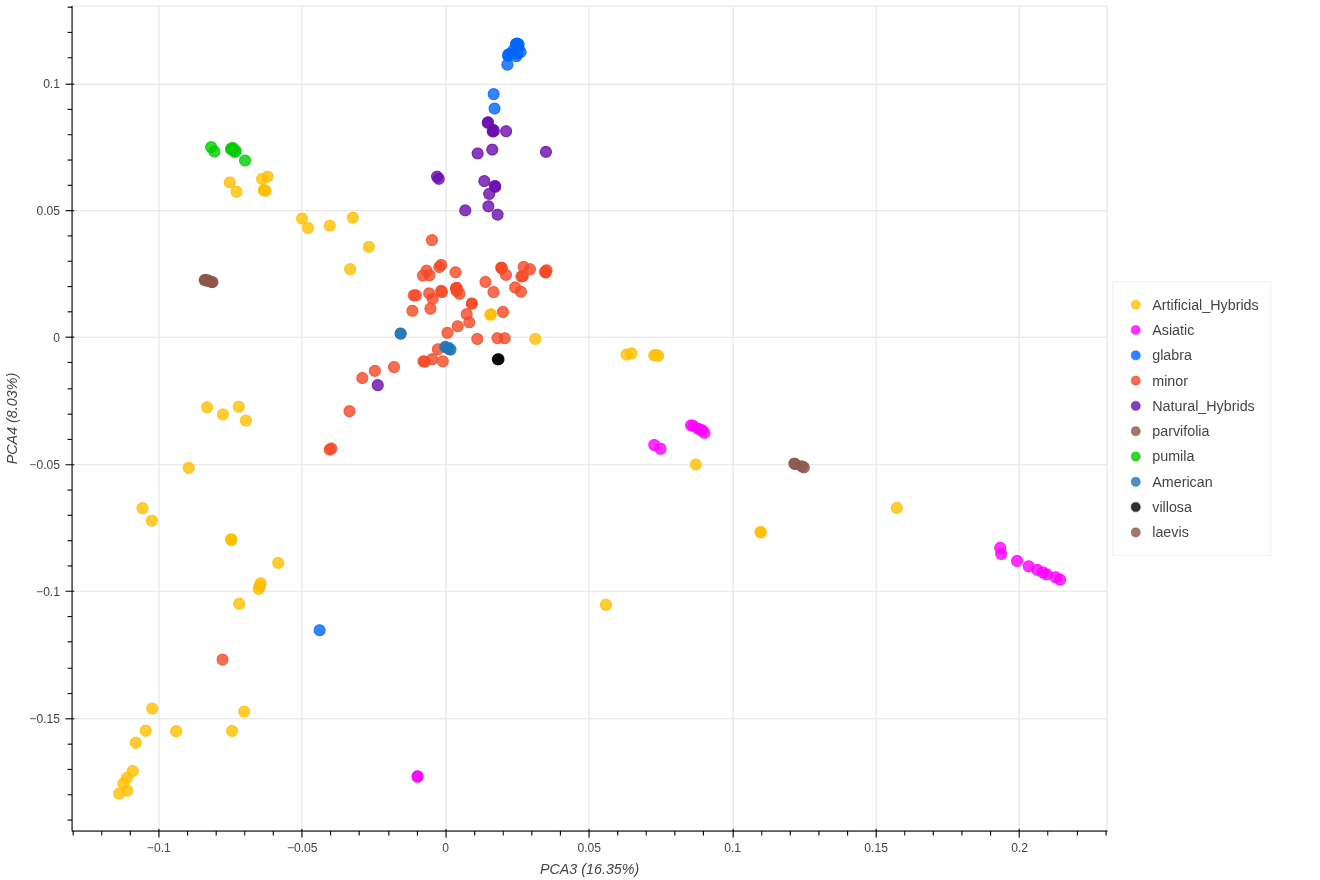

Principal components 3 and 4 are shown.

**(b)**

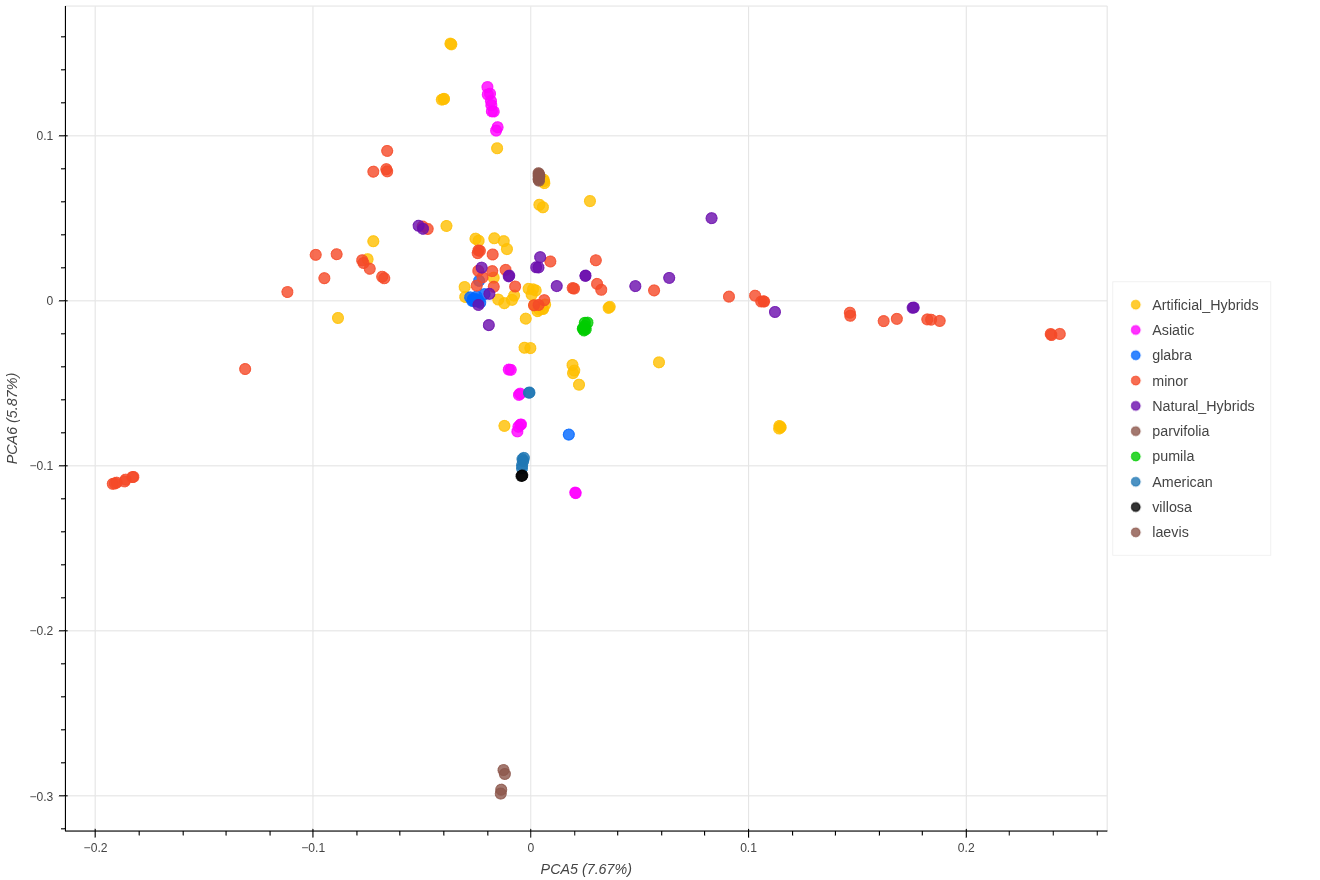

Principal components 5 and 6 are shown.

**(c)**

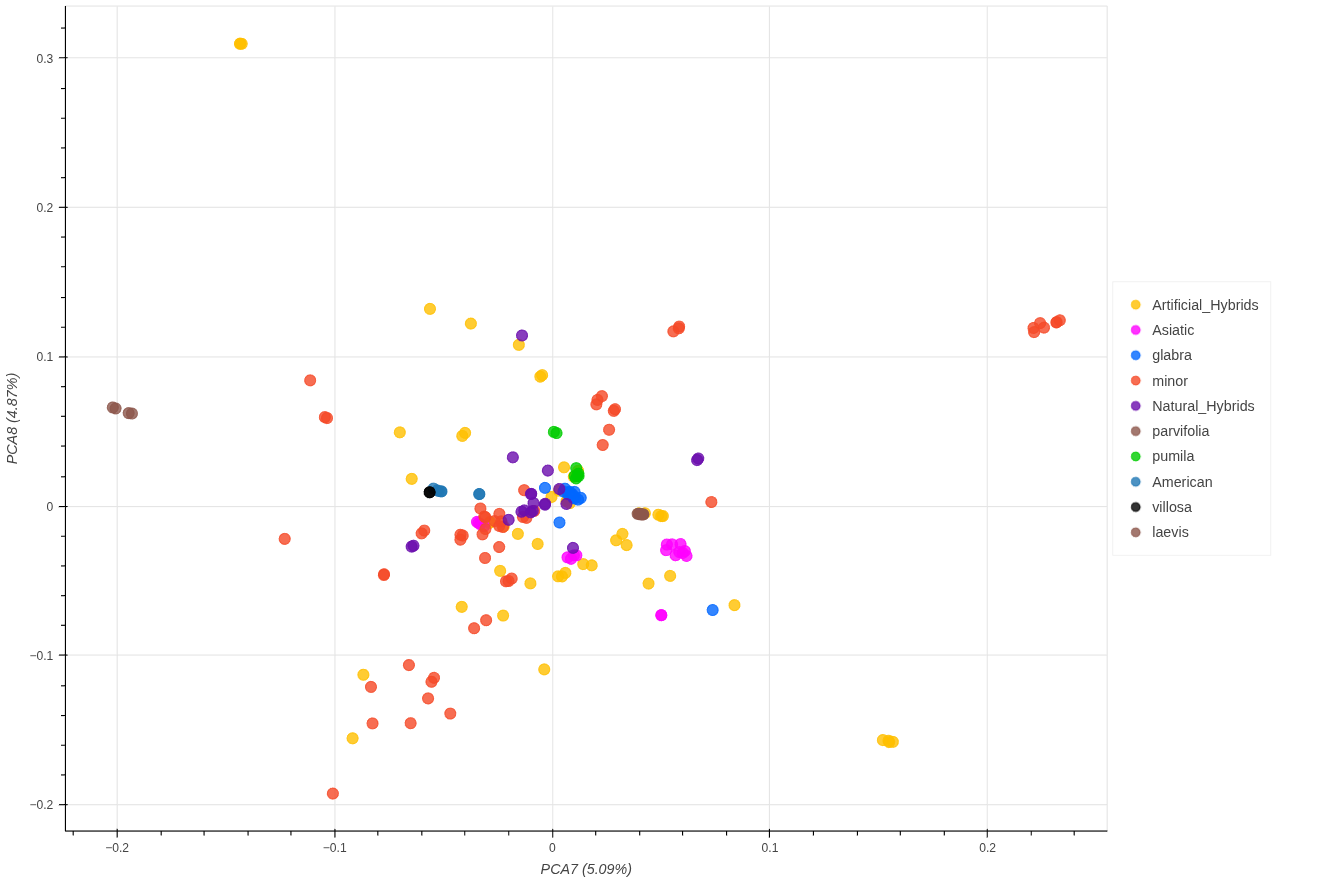

Principal components 7 and 8 are shown.

**(d)**

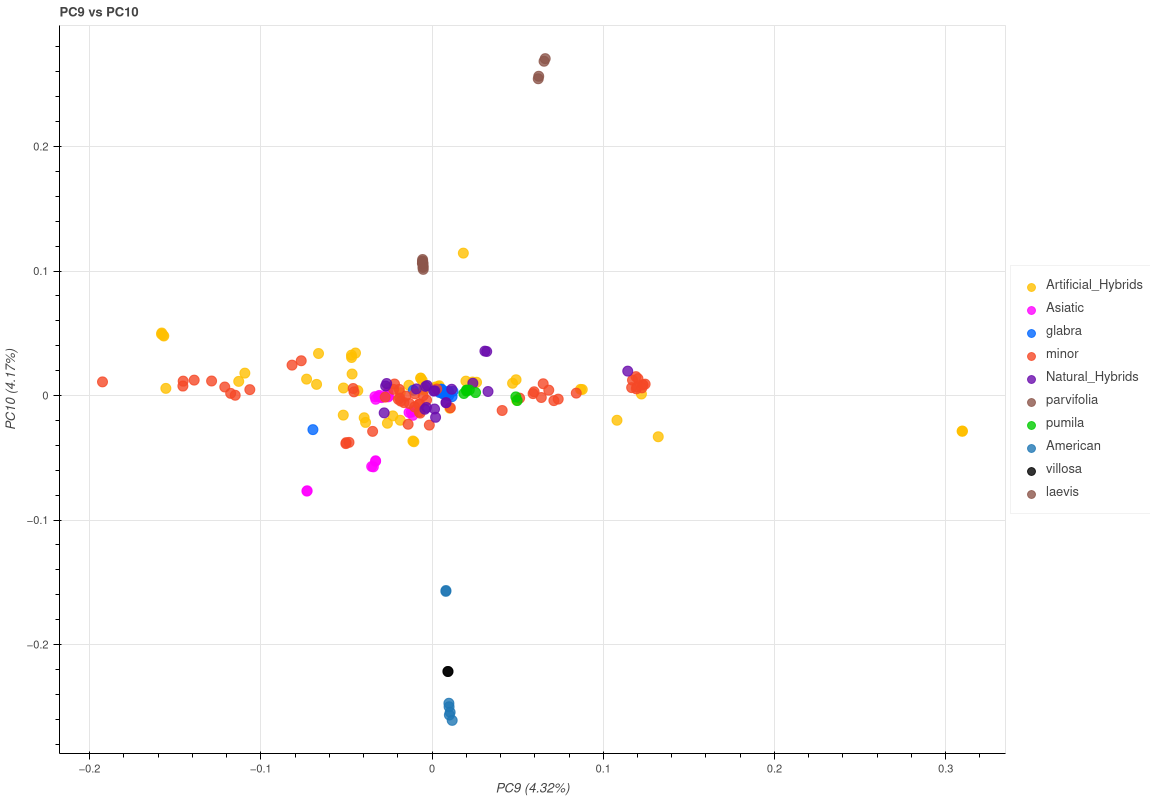

Principal components 9 and 10 are shown.

#### **Fig. S3**. Principal component analysis of whole-genome sequence data from 180 accessions belonging to *Ulmus* subgenus *Ulmus*. Panels show principal components (a) 3 and 4, (b) 5 and 6, (c) 7 and 8, and (d) 9 and 10. Colours and group definitions follow Fig. 2.

## **(a)**

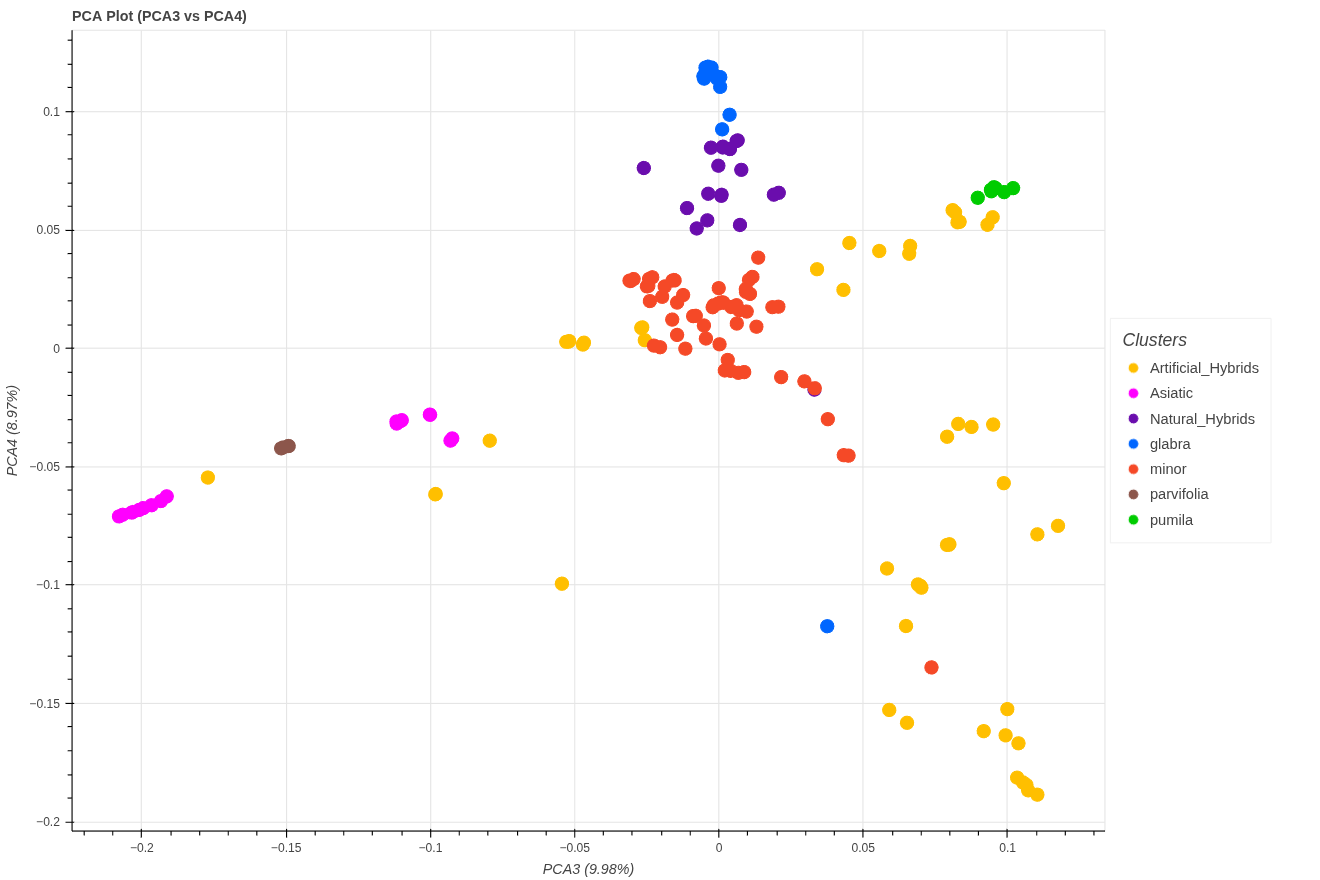

Principal components 3 and 4 are shown.

### **(b)**

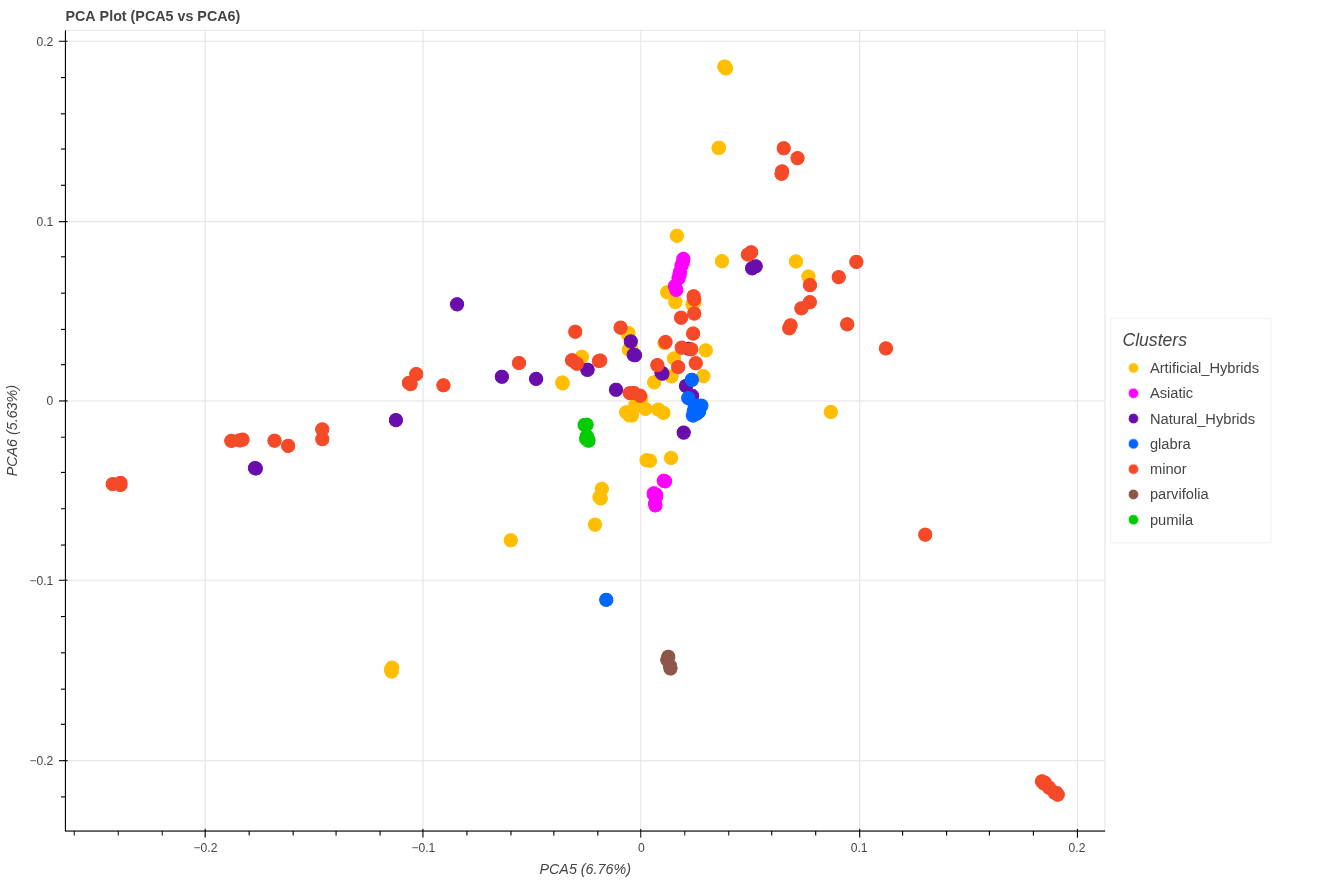

Principal components 5 and 6 are shown.

**(c)**

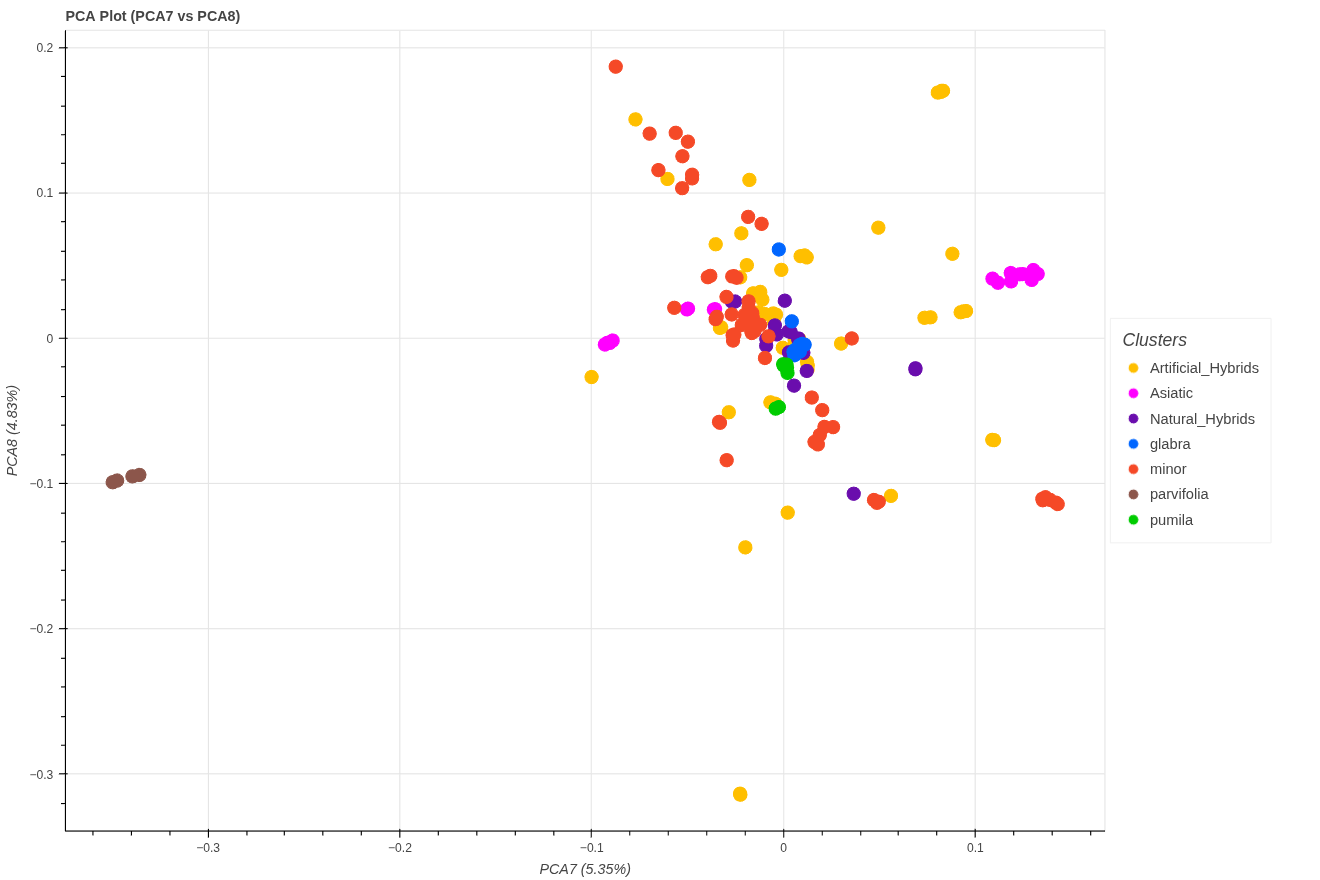

Principal components 7 and 8 are shown.

(d)

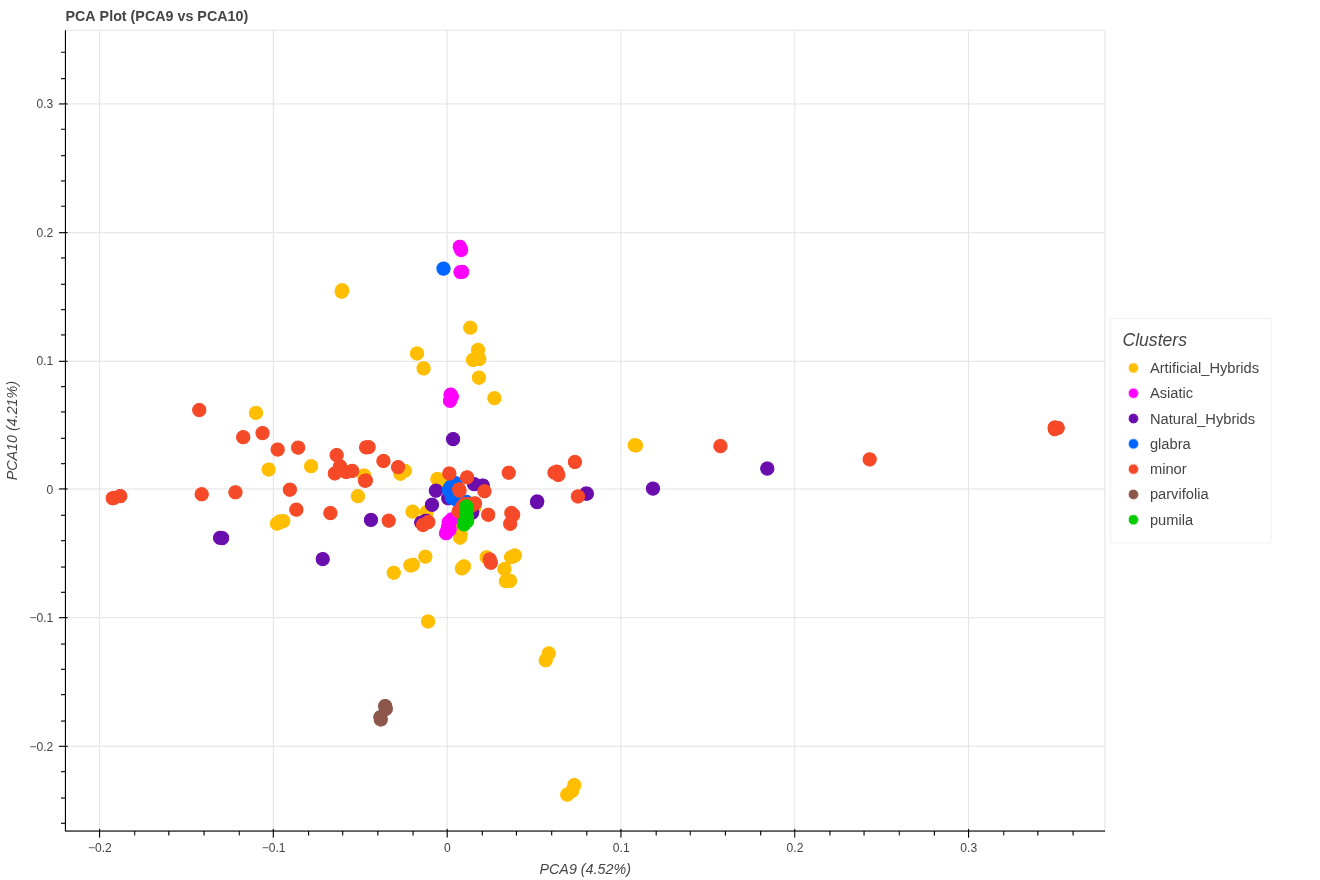

Principal components 9 and 10 are shown.

#### **Fig. S4** Cross-validation results and ADMIXTURE plots for the Elms-120 dataset, comprising mainly natural species and accessions not identified as complex artificial hybrids. The best K of this analysis was K=5, which is presented in the main text (Fig. 4a).

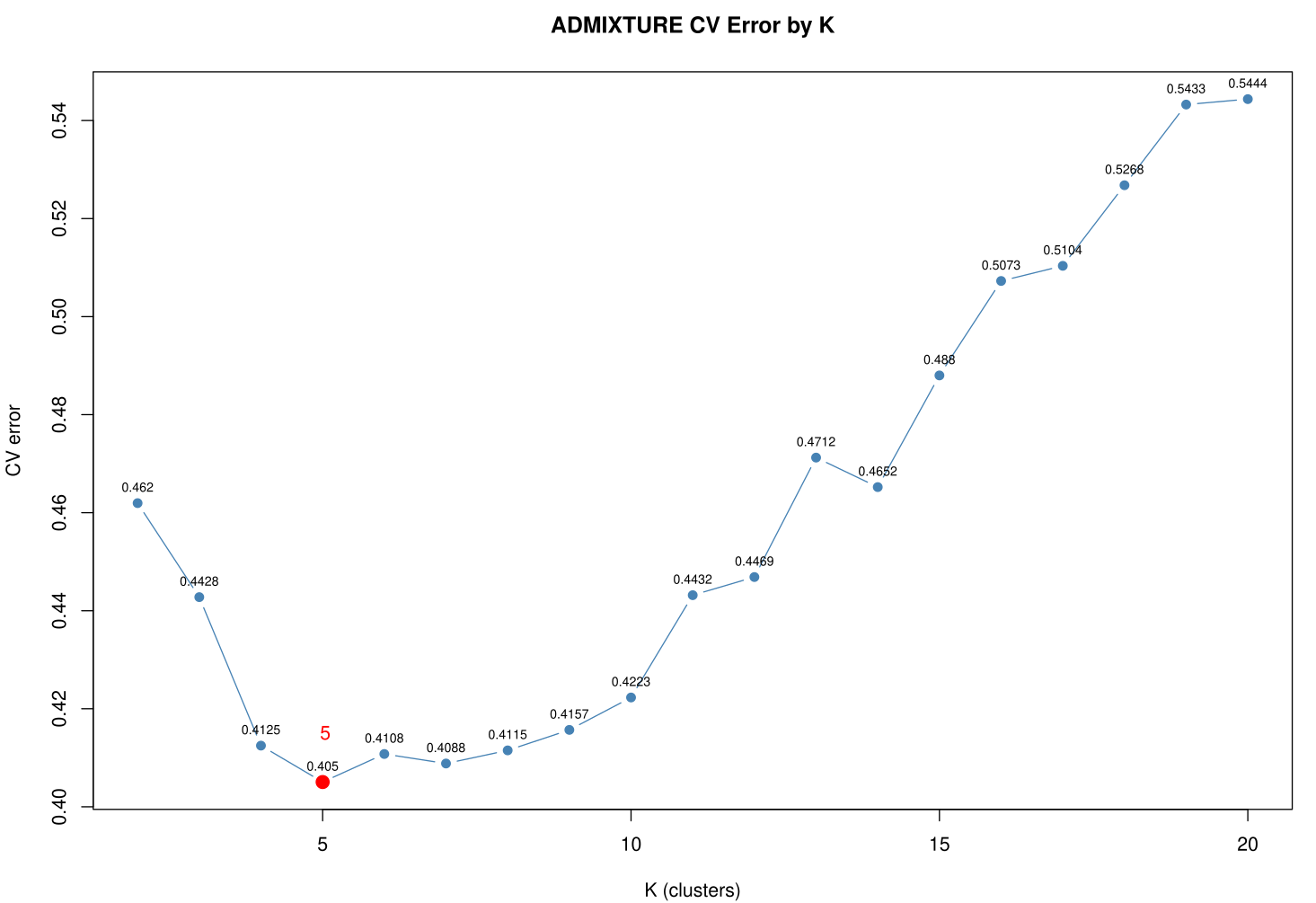

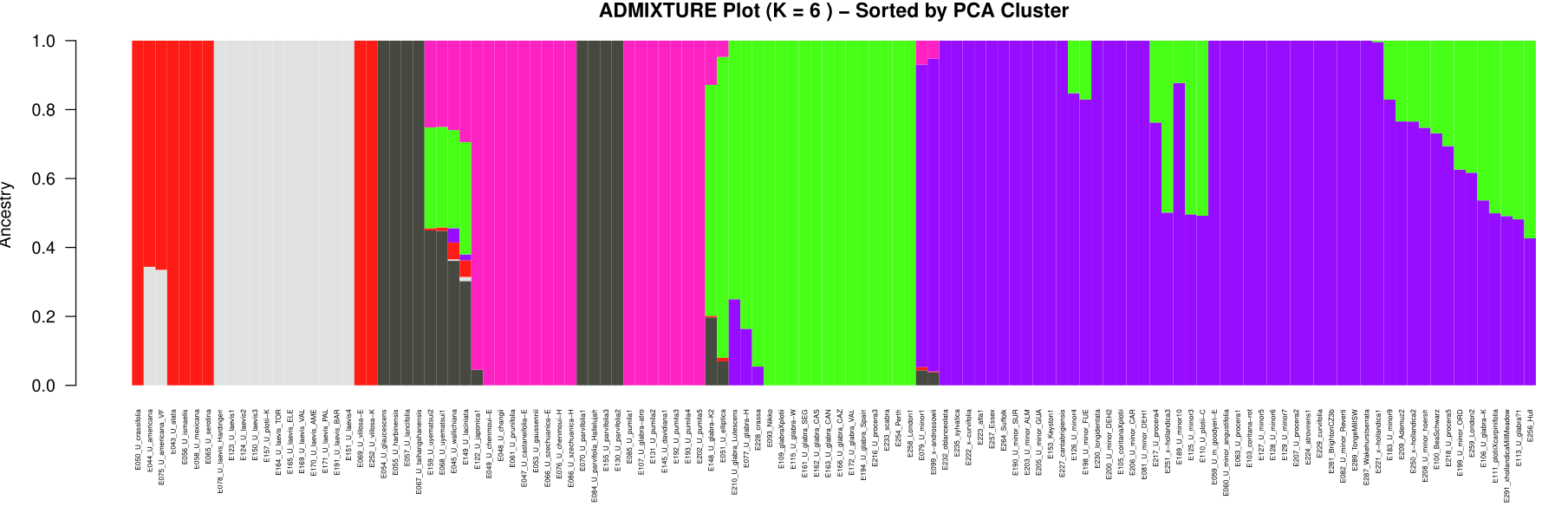

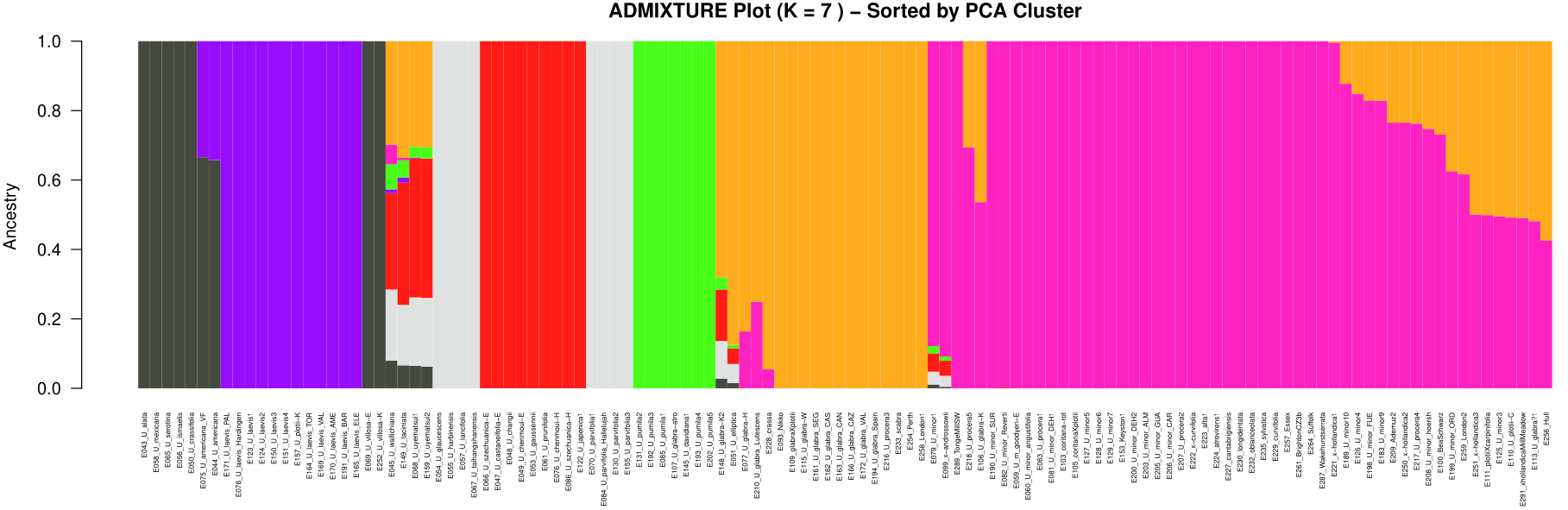

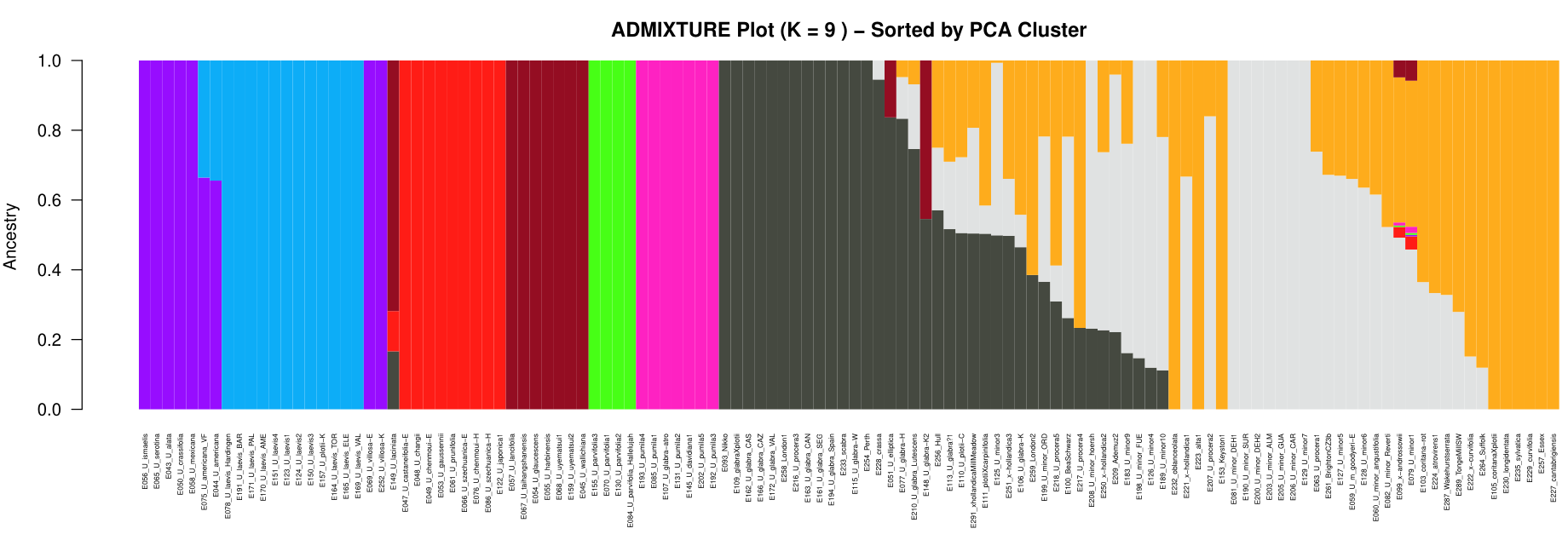

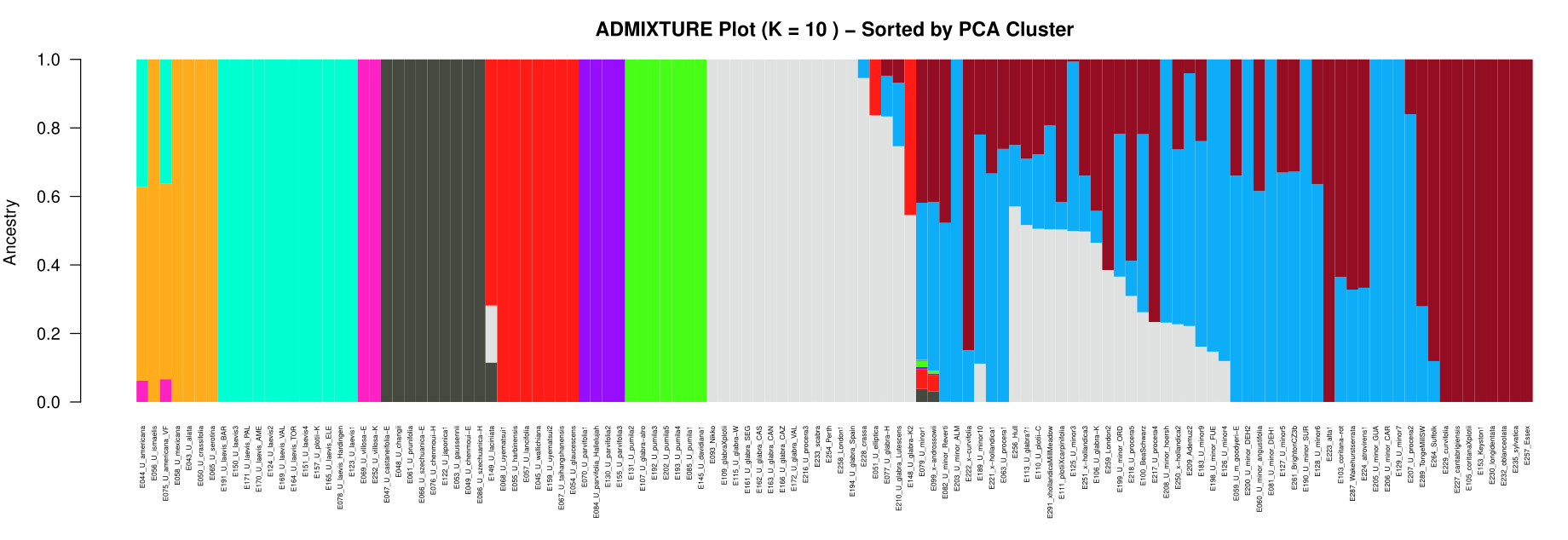

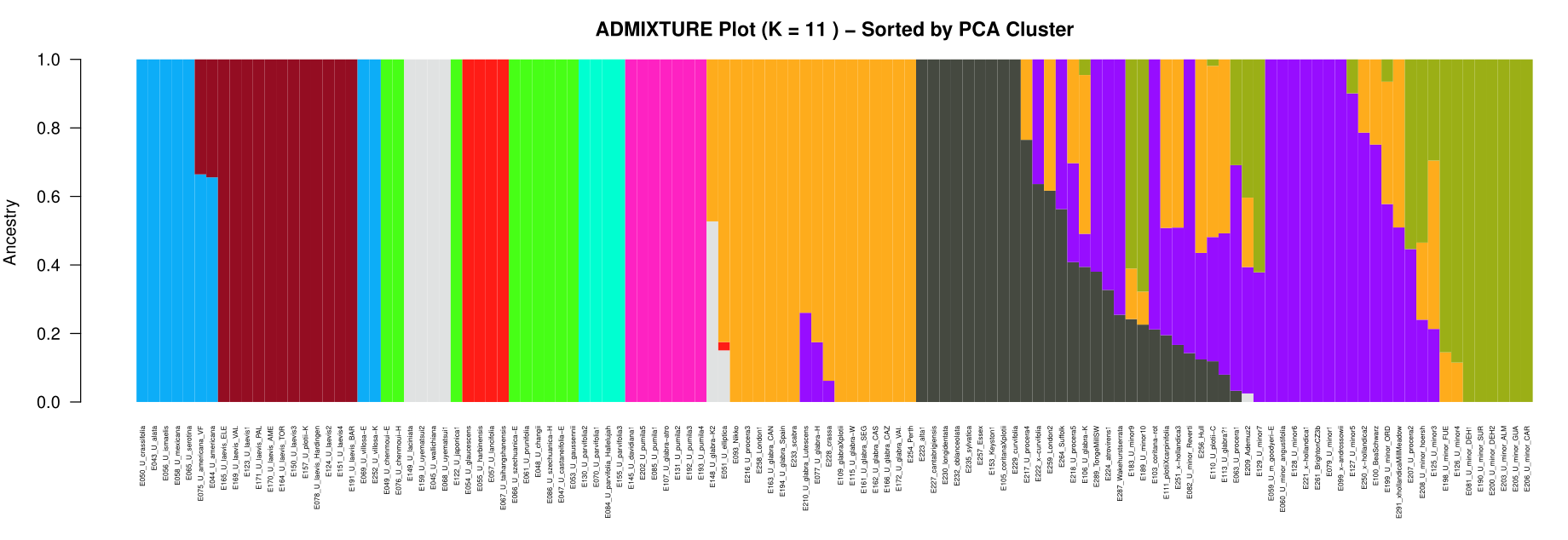

#### **Fig. S5** Cross-validation results and ADMIXTURE plots for the Elms-132 dataset, comprising the Elms-120 accessions plus one representative from each selected clonal group of complex hybrids. The best K of this analysis was K=6 based on cross-validations (CV) from K=2 to K=20 which is presented in the main text (Fig. 5a).

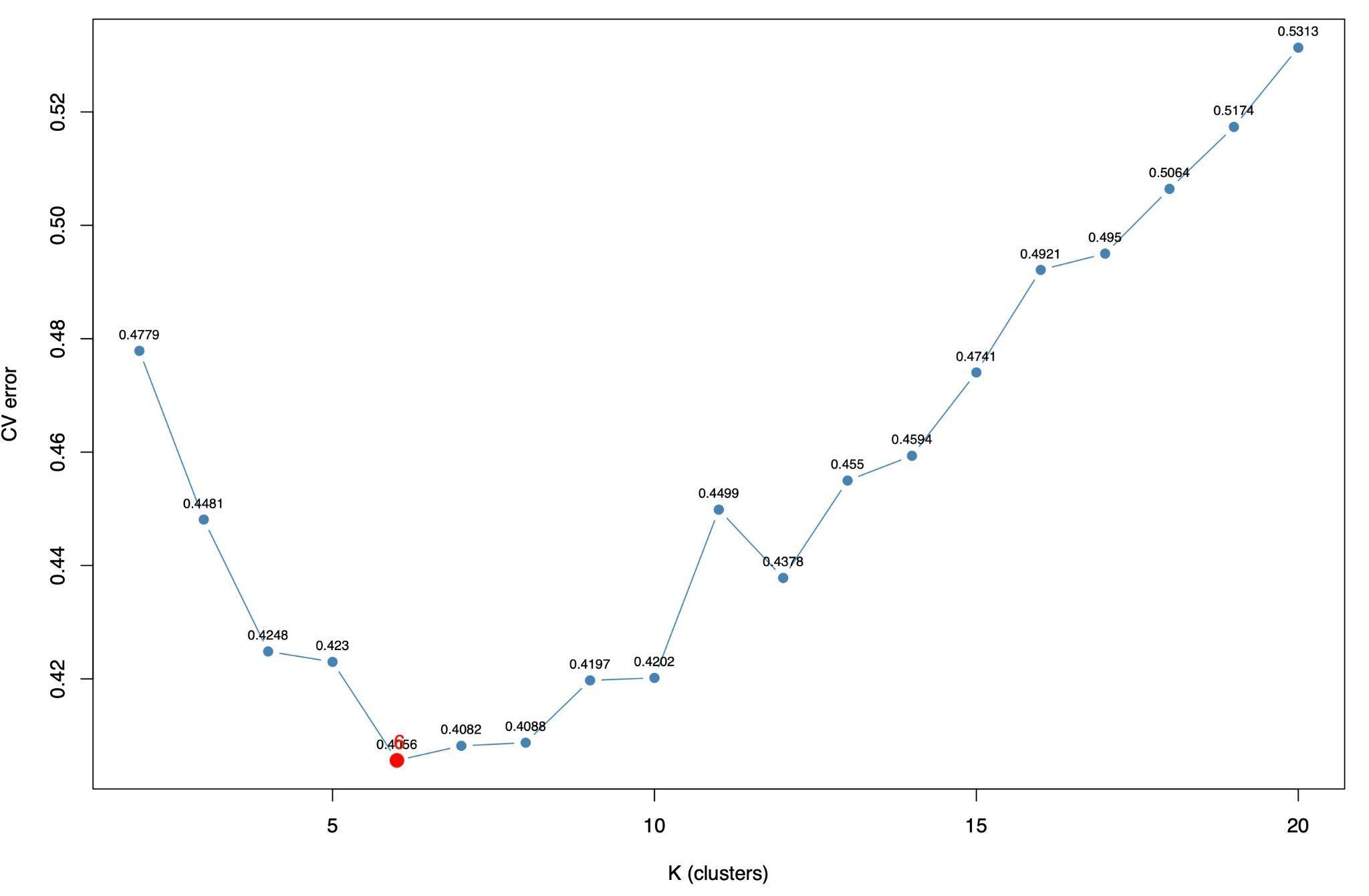

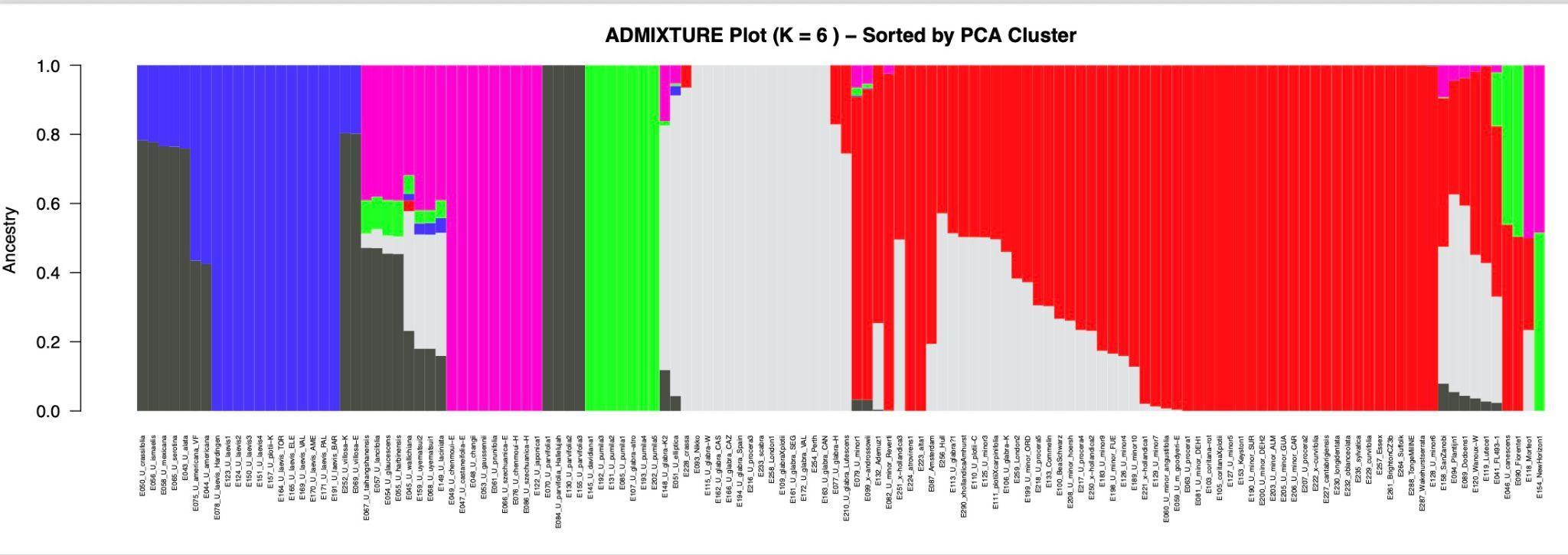

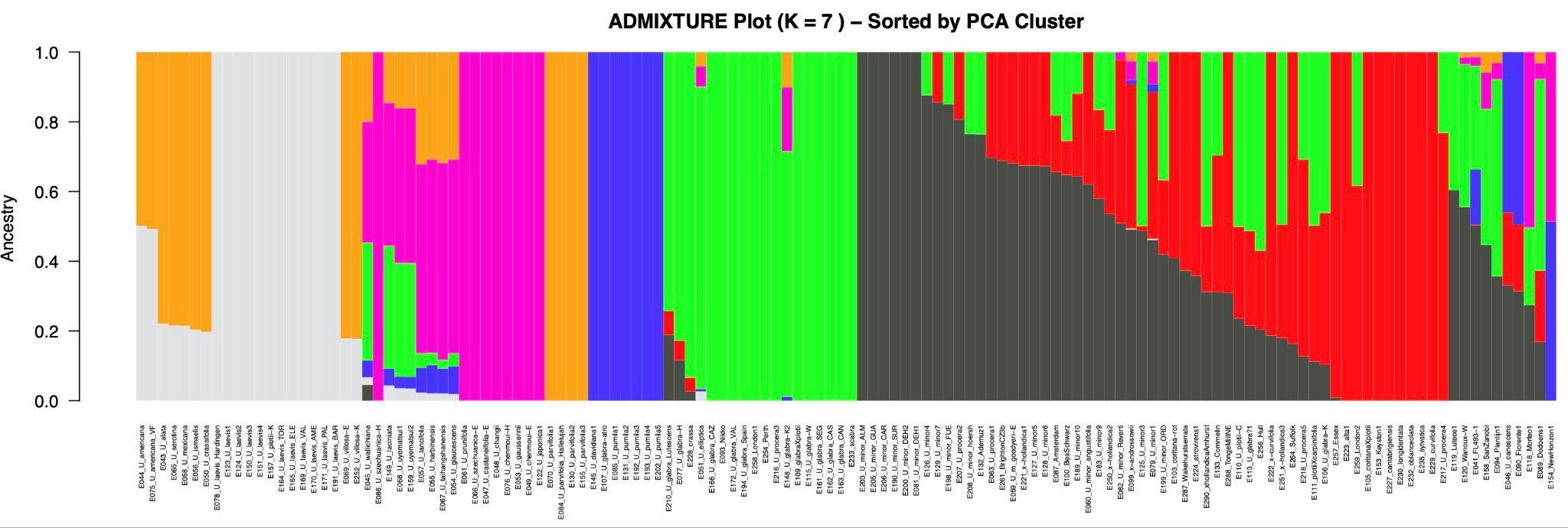

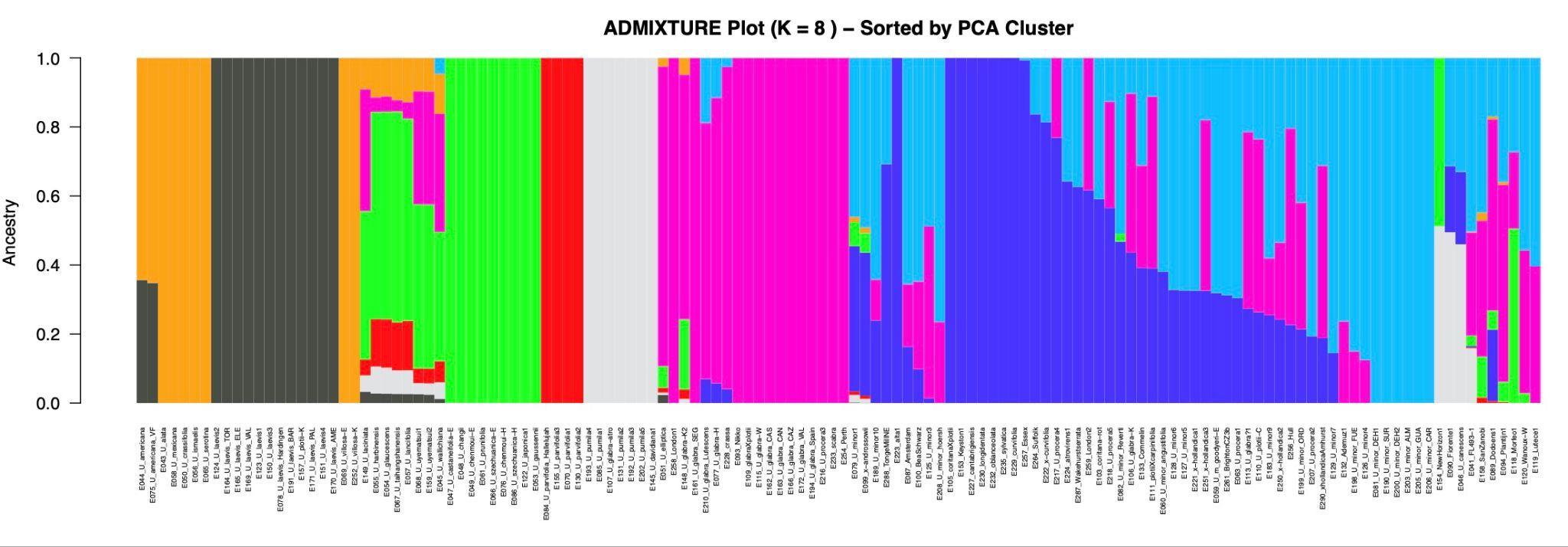

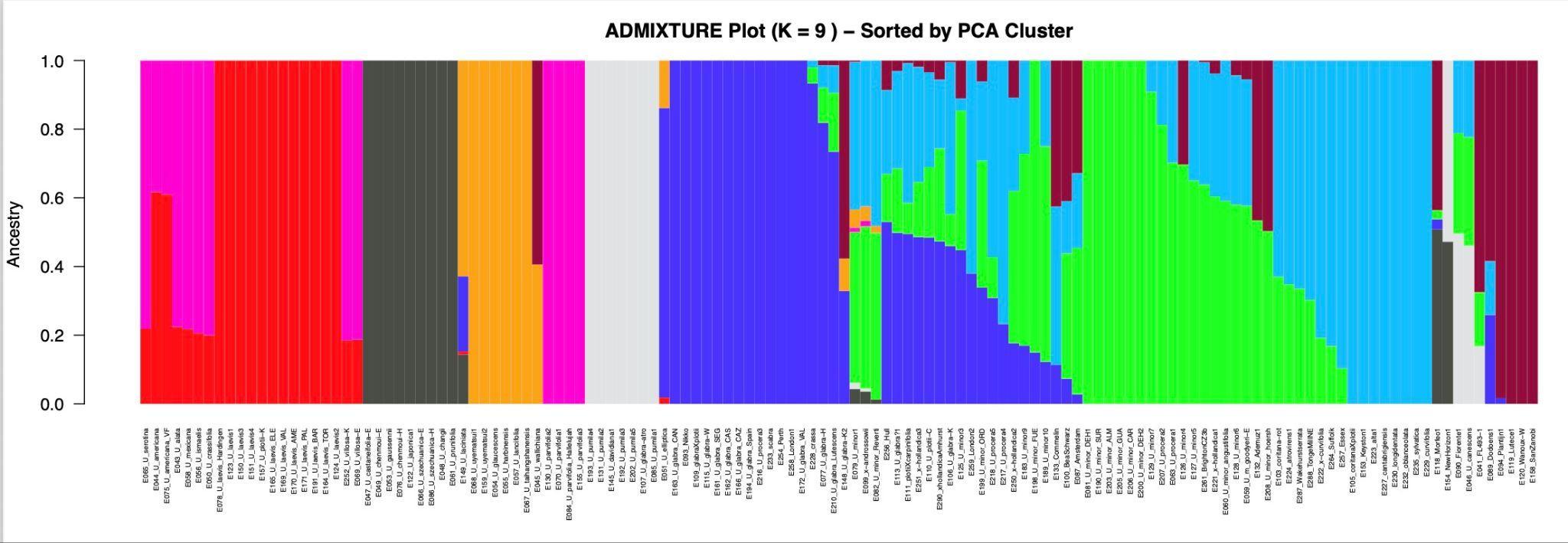

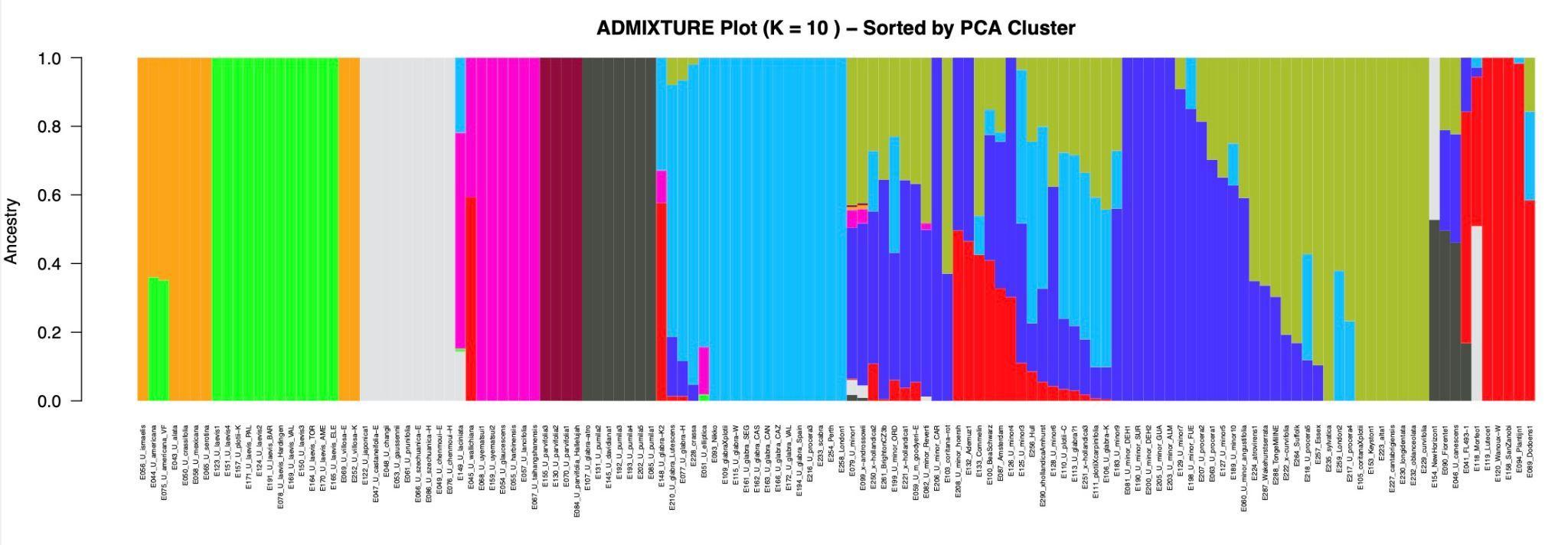

#### **Fig. S6** Genome-wide differentiation (*F_ST_*) between (a) 30 samples of *U. minor* and 13 samples of *U. laevis*; (b) 30 samples of *U. minor* and 15 samples of *U. glabra*; (c) 30 samples of *U. minor* and 21 samples with Sell and Murrell microspecies names: *U. alta, U. atrovirens, U. cantabrigiensis, U. carpinifolia, U. coritana, U. crassa, U. curvifolia, U. longidentata, U. oblanceolata, U. procera, U. scabra, U. serratifrons, U. sylvatica, U. × curvifolia*.

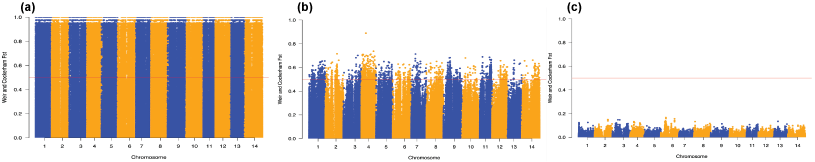

#### **Fig. S7** NeighborNet phylogenetic network inferred from the alignment of 4,651 RADseq loci described for Fig. 3. The dataset comprised 81 accessions newly sequenced in this study and 95 accessions from Whittemore et al. (2021).

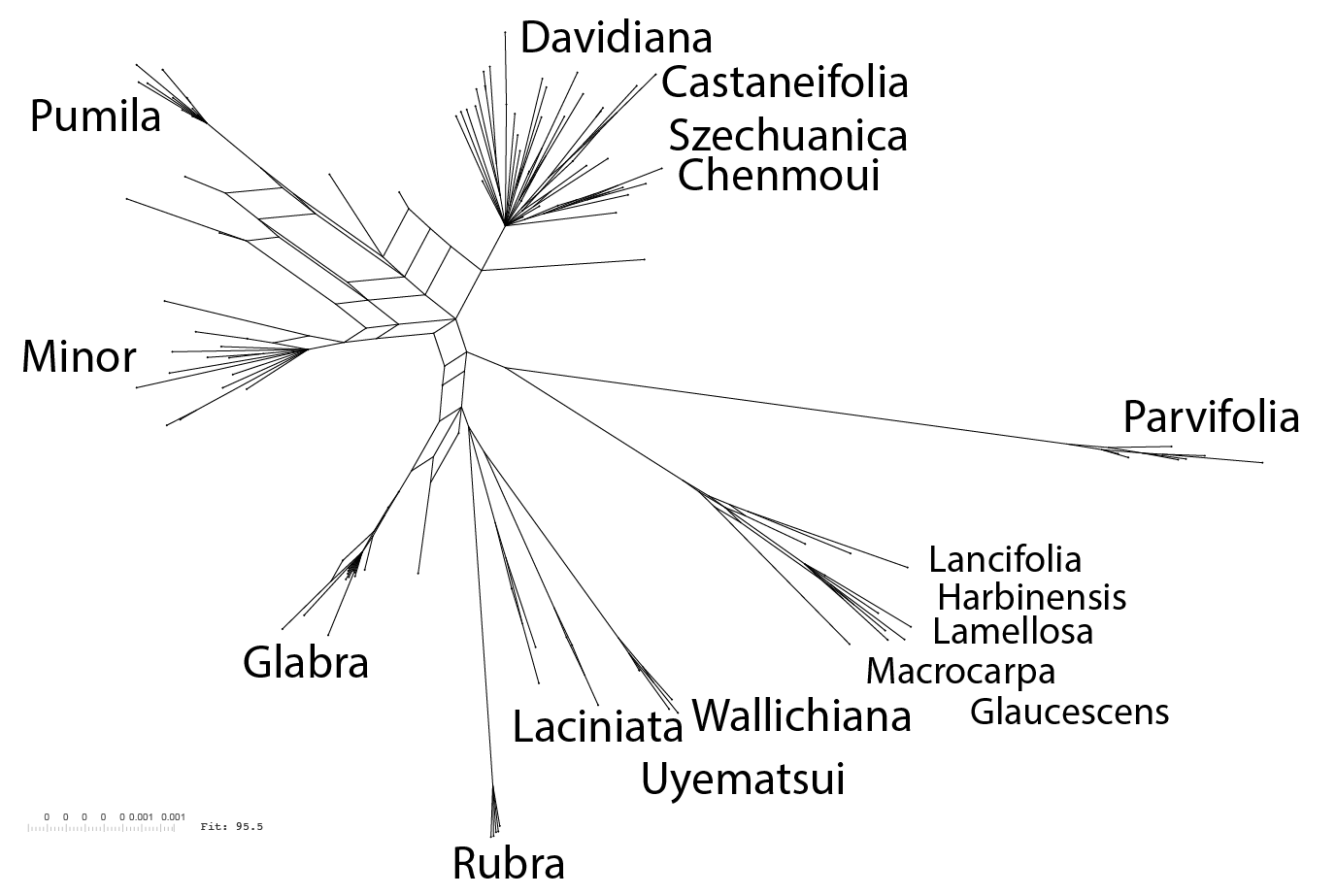

#### **Fig. S8** Nucleotide diversity (𝜋) between 14 individuals of *U. minor* and 14 individuals of *U. glabra*, calculated using VCFtools. (a) Paired window comparison of 𝜋, with *U. glabra* on the x-axis and *U. minor* on the y-axis. Each point represents a paired 1-Mb genomic window. The dashed diagonal indicates equal 𝜋 between species. (b) Distribution of windowed 𝜋 values for the two species. (c) Chromosome-level mean windowed 𝜋 across the 14 elm chromosomes. (d) Difference in windowed 𝜋 between species across 14 chromosomes.

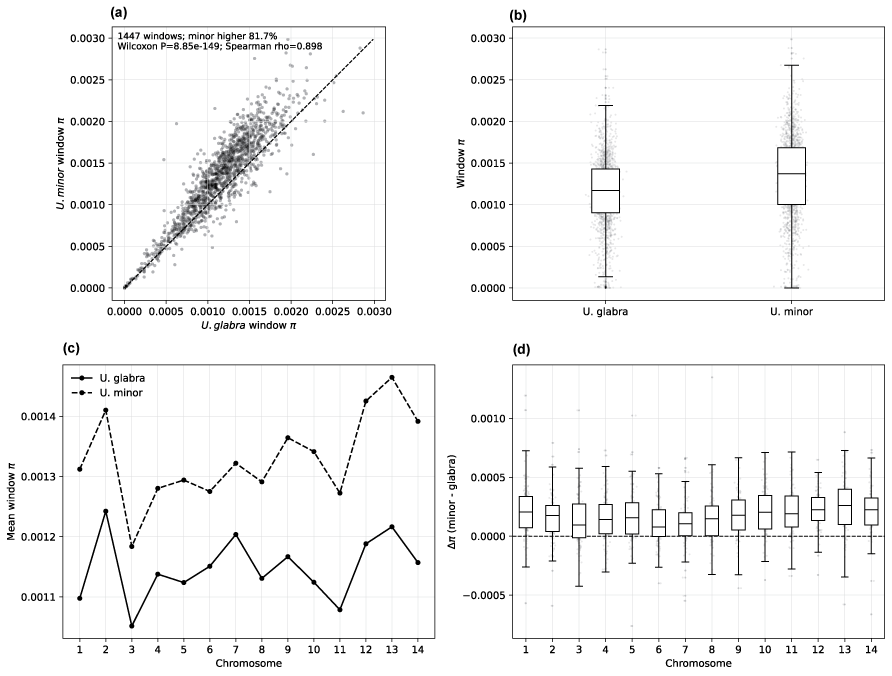

#### **Fig. S9** Genome-wide distribution of SNP density in elm groups made using the CMplot R package. (a) 14 individuals of *U. minor* (7,018,578 SNPs); (b) 14 individuals of *U. glabra* (6,485,481); (c) 17 hybrids between *U. minor* and *U. glabra* (9,160,987); (d) 12 hybrids from the Dutch programme: ‘Dodoens’, ‘Fagel’, ‘Lutece’, ‘Plantijn’, and ‘Wanoux’ (6,067,714); (e) 32 complex hybrids (i.e. *U. canescens*, ‘Fiorente’, FL493, FL610, ‘Lobel’, ‘Den Haag’, ‘Morfeo’, ‘Patriot’, ‘Plinio’, ‘San Zanobi’, and Spain hybrids) (13,656,802); (f) 12 individuals of *U. laevis* (7,246,264); (g) four individuals of *U. parvifolia* (4,249,641). Colour indicates SNP density in 1-Mb windows normalised by the total number of SNPs in each group.

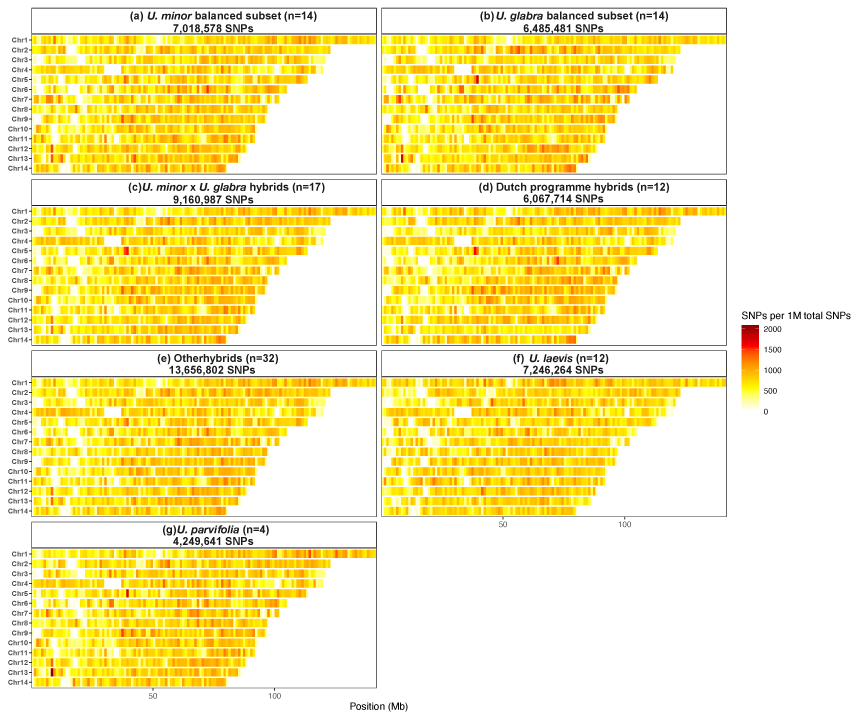
